# A Protein Antibiotic Inhibits the BAM Complex to Kill Without Cell Entry

**DOI:** 10.1101/2025.09.18.677229

**Authors:** Fabian Munder, Matthew Johnson, Imogen Samuels, Laura McCaughey, Oleksii Zdorevskyi, Chunxiao Wang, Ashleigh Kropp, Lauren Zavan, Erin P. Price, Derek S. Sarovich, Swati Varshney, Christopher McDevitt, Hari Venugopal, Vivek Sharma, Matthew T. Doyle, Francesca Short, Debnath Ghosal, James P. R. Connolly, Gavin J. Knott, Rhys Grinter

**Author notes:** These authors contributed equally (Matthew Johnson, Imogen Samuels, Laura McCaughey, Oleksii Zdorevskyi).

## Abstract

Many antibiotics are ineffective against Gram-negative pathogens such as *Pseudomonas aeruginosa* because they cannot penetrate the bacterial outer membrane. Here, we show that protein antibiotics called L-type pyocins kill *P. aeruginosa* by inhibiting the β-barrel assembly machinery (BAM) complex at the cell surface, halting outer-membrane protein assembly. Using single-particle cryo-electron microscopy, we show that L-type pyocins bind a surface-exposed region of BamA and deploy a C-terminal peptide that competitively inhibits the BAM complex, demonstrating that cell entry is not required for antibiotic activity. We combine genetics, multi-omics and cryo-electron tomography to show that BAM complex inhibition by L-type pyocins or the peptide antibiotic darobactin triggers a multifaceted transcriptomic, proteomic and morphological response. Despite this, BAM inhibition ultimately leads to a catastrophic loss of membrane integrity and cell death. These results validate BAM as a target for antibiotics that do not enter the cell and define an engineerable system for their development.

## Introduction

*Pseudomonas aeruginosa* is a significant Gram-negative bacterial pathogen that poses a substantial threat to human health, particularly in individuals with compromised immune systems or those with chronic lung conditions, such as bronchiectasis or chronic obstructive pulmonary disease (COPD)^1–3^. This pathogen causes a wide range of opportunistic infections, from ventilator-associated pneumonia to ear, burn, bloodstream and urinary tract infections, and is notoriously difficult to treat^1^. This difficulty stems in part from its outer membrane, which acts as a barrier to many antibiotics, conferring a high level of intrinsic resistance^4,5^. In addition, *P. aeruginosa* readily acquires resistance through horizontal gene transfer and mutation, using mechanisms such as efflux pumps, antibiotic-modifying enzymes, and alterations in membrane permeability^1,5^. The World Health Organization classifies *P. aeruginosa* as a “high-priority pathogen” for the development of new therapeutics^6^. Despite the pressing need, few new antibiotic classes with efficacy against this organism have been developed^1^. To date, only two essential outer membrane protein targets have been identified where inhibition leads to bacterial death: the LptDE complex, responsible for inserting lipopolysaccharide (LPS) into the outer leaflet of the outer membrane, and the β-barrel assembly machinery (BAM) complex, which mediates the folding and insertion of integral outer membrane proteins (OMPs)^7–11^. These complexes are increasingly attracting attention as targets for next-generation antibiotics^7^.

L-type pyocins are protein antibiotics produced by *P. aeruginosa* to eliminate closely related competitor strains and species^12–14^. These molecules display potent and specific antibacterial activity and have shown therapeutic promise in preclinical infection models, including the clearance of lethal *P. aeruginosa* lung infections in mice^15^. L-type pyocins recognize specific molecules on the surface of their target cell, achieving per-molecule potencies several orders of magnitude higher than small-molecule antibiotics^12,13^. Their specificity also enables targeted killing of *P. aeruginosa* without off-target effects to the host microbiota, making them attractive candidates for biologic therapies^14^.

Structurally, L-type pyocins are composed of N- and C-terminal β-prism lectin domains connected to an unstructured C-terminal peptide. The C-terminal β-prism domain contains conserved motifs that bind to D-rhamnose residues in the common polysaccharide antigen (CPA) of *P. aeruginosa* LPS, which serves as the primary cell surface receptor^12,16^. Strains lacking the CPA O-antigen are more tolerant to L-type pyocins, underscoring the importance of CPA binding for initial recognition^12,17^. However, CPA binding alone cannot explain the lethal activity of these proteins. Importantly, the deletion of three final residues of the C-terminal peptide from this family of protein antibiotics abolishes bactericidal function, suggesting that this region plays a critical but poorly understood role in L-pyocin function^16^.

Recent evidence implicates the BAM complex as a cell surface receptor for L-type pyocins, and potentially as their cytotoxic target. The BAM complex catalyzes the folding and insertion of outer membrane proteins (OMPs), with β-strand 1 of the BamA β-barrel binding an incoming OMP’s C-terminal β-strand to nucleate and guide β-barrel formation^18–21^. The BamA β-barrel consists of periplasmic turns and extracellular loops, with the sequence of loop 6 correlating with a strain’s susceptibility to L-type pyocins^13^. This observation points to an interaction between L-type pyocins and the BAM complex, yet its mechanistic basis and downstream consequences remain unresolved. Despite their promise as antimicrobials, the engineering and application of L-type pyocins have been hindered by a lack of understanding of their cytotoxic mechanism and the downstream consequences of intoxication.

In this study, we comprehensively dissect the L-type pyocin mechanism of action. We expand the L-type pyocin toolkit and establish the molecular determinants of activity across a panel of clinical *P. aeruginosa* isolates. We isolate the BAM complex from *P. aeruginosa* and determine the structural and mechanistic basis of L-type pyocin-BamA interaction using single-particle cryo-electron microscopy (cryo-EM) and molecular dynamics simulations. To determine the cellular consequences of L-type pyocin intoxication, we combine transposon-directed insertion site sequencing (TraDIS), transcriptomics, proteomics, and cryo-electron tomography (cryo-ET). We find that L-type pyocins deploy their C-terminal peptide to bind the BAM complex at β-strand 1, resulting in inhibition of outer membrane protein assembly, induction of envelope stress responses, and ultimately, bacterial cell death, resulting from a loss of integrity of the outer membrane. This process makes *P. aeruginosa* cells susceptible to co-treatment with existing antibiotics that are otherwise ineffective. These findings validate the BAM complex as a key vulnerability in Gram-negative bacteria that can be targeted by antibiotics that do not enter the bacterial cell and demonstrate the therapeutic potential of L-type pyocins targeting outer membrane biogenesis.

## Main

### L-type pyocins are potent inhibitors of *P. aeruginosa*

Previous work assessed the inhibition of *P. aeruginosa* by L-type pyocins in a binary growth/no-growth assay, with minimum inhibitory concentrations (MICs) as low as 6.9 nM reported^17^. To gain a more nuanced picture of L-type pyocin activity, we performed *P. aeruginosa* PAO1 growth curves in liquid culture in the presence of Pyocin L1 (HG422551), ranging from 3.27 µM to 0.033 nM. Pyocin L1 affected the growth of *P. aeruginosa* PAO1 at concentrations higher than 0.33 nM, with a half-maximal inhibitory concentration (IC_50_) of 0.85 nM (**Fig. 1A,B**). At higher concentrations, growth slowed 2 hours post-inoculation, arrested by ∼6 hours, and subsequently declined. An optical density of at least 0.75 was obtained even at higher Pyocin L1 concentrations, which may explain the lower IC_50_ calculated by our experiments, compared to the earlier binary inhibition assays (**Fig. 1B**). These data indicate that L-type pyocins can be at least 1000 times more potent, on a per molecule basis, than antibiotics commonly used to treat infection by *P. aeruginosa*^22–25^. In addition to Pyocin L1, previous reports identified Pyocin L2 and L3, which share 86% and 30% sequence identity with Pyocin L1, and exhibit distinct but overlapping spectra of activity^12,17^. To expand our repertoire of L-type pyocins for development as antibiotics targeting *P. aeruginosa*, we searched available microbial genomes, identifying four additional L-type pyocins, which we designated Pyocin L2b, L3b, L4 and L5. These new L-type pyocins share between 28-96% sequence identity, with ‘b’ designating >90% identity to an existing protein (**Fig. 1C,D**).

**Figure 1:**
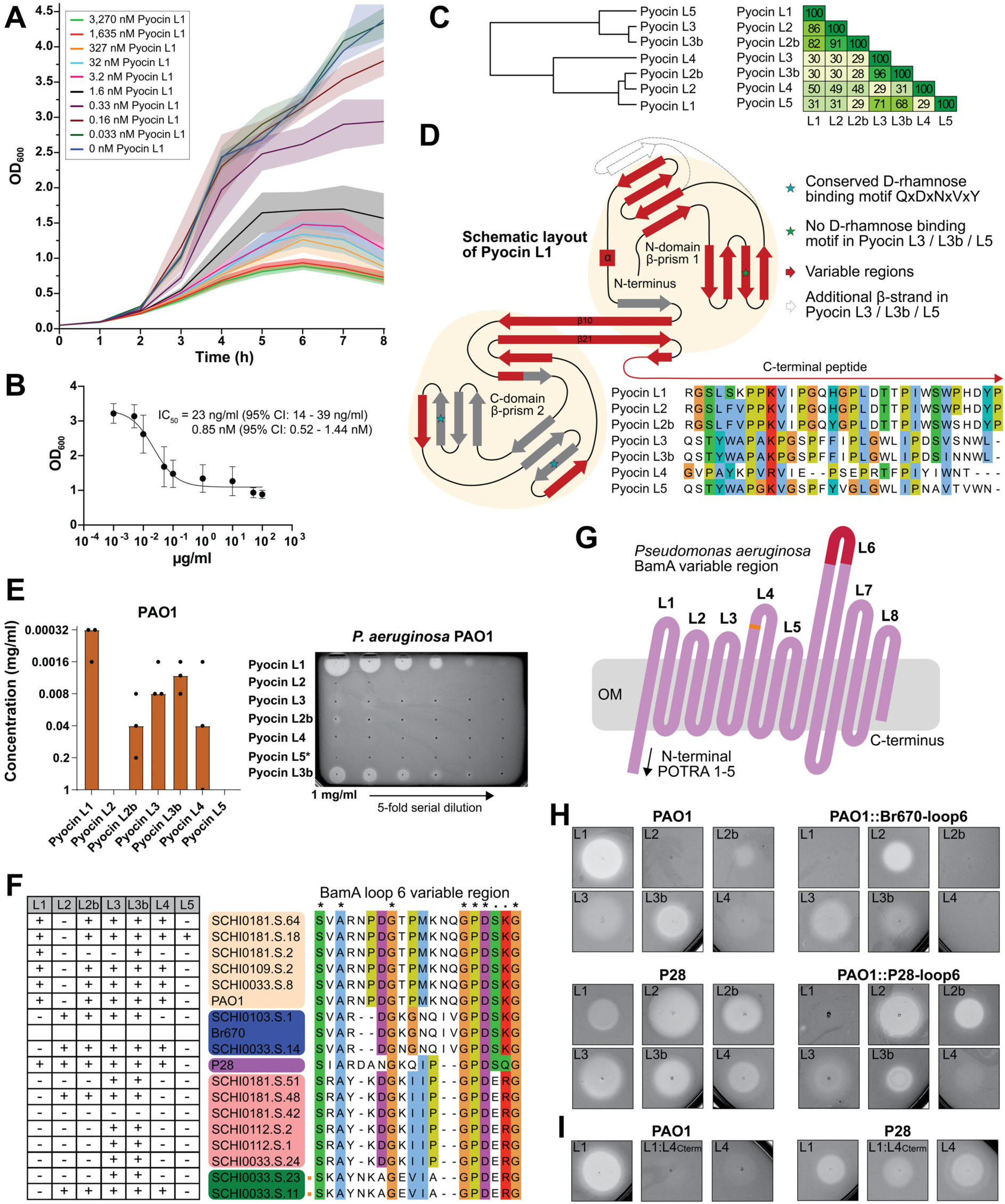
Discovery and functional characterisation of novel L-type pyocins. (A) Growth curve of *P. aeruginosa* PAO1 with Pyocin L1 treatment at various concentrations, SEM is shown as a shaded area (n=5). (B) IC^50^ plot of Pyocin L1-treated *P. aeruginosa* cultures, based on OD_600_ 6 hours post-treatment. Error bars show SEM (n=5). (C) A tree showing L-type pyocin phylogeny and a percentage sequence identity matrix. (D) A schematic of the Pyocin L1 structure and sequence alignment of the C-terminal peptide of Pyocin L1 to L5. Red regions have high sequence variability, and grey regions are conserved. Stars indicate rhamnose-binding motifs, and dotted lines indicate an additional β-strand for Pyocin L3, L3b, and L5. (E) A plot of results from plate-based inhibition screens of Pyocin L1-L5 against *P. aeruginosa* PAO1 (n=3) (left), and example inhibition assay plate (right). (F) An overview of the susceptibility of diverse clinical *P. aeruginosa* strains to Pyocins L1-5, and a sequence alignment of their BamA loop 6 region (See **Figure S2**, for detailed data on the inhibition of these strains). (G) A schematic of *P. aeruginosa* BamA. The main region of sequence variation at loop 6 is shown in red, with a minor region in loop 4 shown in orange. (H) Plate-based inhibition screens with Pyocin L1-L4 against PAO1, P28, *P. aeruginosa* PAO1::P28-loop6 and PAO1::Br670-loop6. (I) A plate-based inhibition screen of *P. aeruginosa* PAO1 and P28 with Pyocin L1, the chimeric of Pyocin L1 and the last 24 C-terminal residues of Pyocin L4, and Pyocin L4.

### L-type pyocins inhibit diverse pathogenic *P. aeruginosa* strains

We recombinantly produced our seven L-type pyocins and performed plate-based growth inhibition assays against 17 *P. aeruginosa* strains, including 15 recent clinical isolates^26^. Previous reports indicate that a C-terminal affinity tag can impact the activity of L-type pyocins^17,27,28^. Accordingly, they were purified using an N-terminal poly-histidine tag, except Pyocin L5, which could only be purified using a C-terminal His-tag (**Fig. S1A**). The *P. aeruginosa* isolates displayed variable susceptibility to the L-type pyocins in our panel. Sixteen of the seventeen strains were susceptible to at least one L-type pyocin, with the majority potently inhibited by multiple proteins (**Fig. 1E,F, Fig. S2**). Pyocin L5 was the least effective L-type pyocin, with weak activity observed only against one bronchiectasis-derived *P. aeruginosa* strain (SCHI0181.S.18). However, given that its C-terminal His-tag may affect its activity, this may not represent its true killing spectrum.

Previous work demonstrated a correlation between L-type pyocin susceptibility and the amino acid sequence of extracellular loop 6 of BamA (**Fig. 1G**)^13,29–32^. We identified five BamA loop 6 types in *P. aeruginosa* strains, which correlated with the L-type pyocin susceptibility (**Fig. 1F, Fig. S2**). However, this correlation is not absolute, with both susceptibility and sensitivity varying among strains sharing the same BamA loop 6 sequences (**Fig. 1F, Fig. S2**). To assess BamA loop 6 diversity in *P. aeruginosa*, we analysed a collection of 557 clinical and environmental isolates (**Supplementary Dataset S1**). The *bamA* gene was highly conserved (97.9% nucleic acid and 98.6% amino acid pairwise identity) with variation predominantly confined to loop 6 (**Fig. S3A,B**). This suggests this region is under strong diversifying selection pressure. To identify variations in loop 6, the 557 *P. aeruginosa* BamA sequences were analysed for unique amino acid sequences in this region. We identified eleven loop 6 sequence types, including the six represented in the isolates in our susceptibility screen, which correspond to 427 (76%) of the strains analysed (**Fig. S3C**). Notably, one of the nine loop 6 variants was represented by a shorter sequence that was present in 116 strains (21%) and was not represented in the screened isolates. Thus, the L-type pyocins we identified target the majority of BamA loop 6 sequence variants across a broad panel of *P. aeruginosa* clinical isolates.

Based on the importance of BamA loop 6 for L-type pyocin susceptibility, we substituted loop 6 in *P. aeruginosa* PAO1 with that of *P. aeruginosa* strains P28 and Br670, by allelic exchange (**Fig. 1H**). In plate-based growth inhibition assays, loop 6 substitution altered the L-type pyocin susceptibility of the *P. aeruginosa* PAO1::P28-loop6 and PAO1::Br670-loop6 strains towards that of loop 6 donor strains (**Fig. 1H**). Like *P. aeruginosa* P28 and Br670, both substitution strains were susceptible to Pyocin L2. Pyocin L3b, which is active against wild-type *P. aeruginosa* PAO1 and P28, had reduced activity against PAO1::P28-loop6 and PAO1::Br670-loop6. Interestingly, *P. aeruginosa* PAO1::P28-loop6 is resistant to Pyocin L1, even though Pyocin L1 is active against both *P. aeruginosa* PAO1 and P28 (**Fig. 1H**). Pyocin L1 activity towards *P. aeruginosa* P28 is weaker than towards PAO1, suggesting that the P28 loop 6 sequence is not a strong determinant of Pyocin L1 activity, and that other factors contribute towards P28 susceptibility relative to PAO1. These results strongly support the correlation between loop 6 sequence and L-type pyocin susceptibility but also show that loop 6 is not the exclusive determinant of activity.

In addition to tandem β-lectin domains, L-type pyocins also possess a C-terminal peptide rich in proline and aromatic residues^12,16^. The sequence of this region varies considerably among the L-type pyocins in our study in both in amino acid composition and length (**Fig. 1D**). This region is critical for the activity of other members of this family, with its deletion abolishing the activity of the related protein LlpA^16^. To assess the importance of the C-terminal peptide for L-type pyocin activity, we created a chimeric version of Pyocin L1 with the C-terminal peptide (last 24 residues) of Pyocin L4, which we designated Pyocin L1:L4_cterm_. Plate-based inhibition assays showed that Pyocin L1:L4_cterm_ inhibits the growth of *P. aeruginosa* PAO1 and P28, but with lower efficacy than Pyocin L1 or L4. Unlike Pyocin L1, which strongly inhibits *P. aeruginosa* PAO1, Pyocin L1:L4_cterm_ is similar in potency to Pyocin L4, which only weakly inhibits this strain. Alternatively, Pyocin L1:L4_cterm_ inhibition of *P. aeruginosa* P28 is more like both Pyocin L1 and L4, which both moderately inhibit this strain, demonstrating that different C-terminal peptides affect *P. aeruginosa* strains in different ways (**Fig. 1I**).

### The BAM complex is the target of L-type pyocins

To test whether the correlation between BamA loop 6 and L-type pyocin susceptibility results from a direct interaction between these proteins^13^, we chromosomally introduced a 2xStrep-tag at the N-terminus of BamA of *P. aeruginosa* PAO1 and natively isolated it in detergent micelles. All five BAM subunits were present in this complex (BamA-E), as well as a significant quantity of the LPS insertase LptD (**Fig. 2A**). We utilized cryo-EM to resolve the structure of the *P. aeruginosa* PAO1 BAM complex (BAM_PAO1_) to 2.75 Å, in an outward open conformation (**Fig. 2B,C, Fig. S4, Fig. S5, Table S1, Movie S1**).

**Figure 2:**
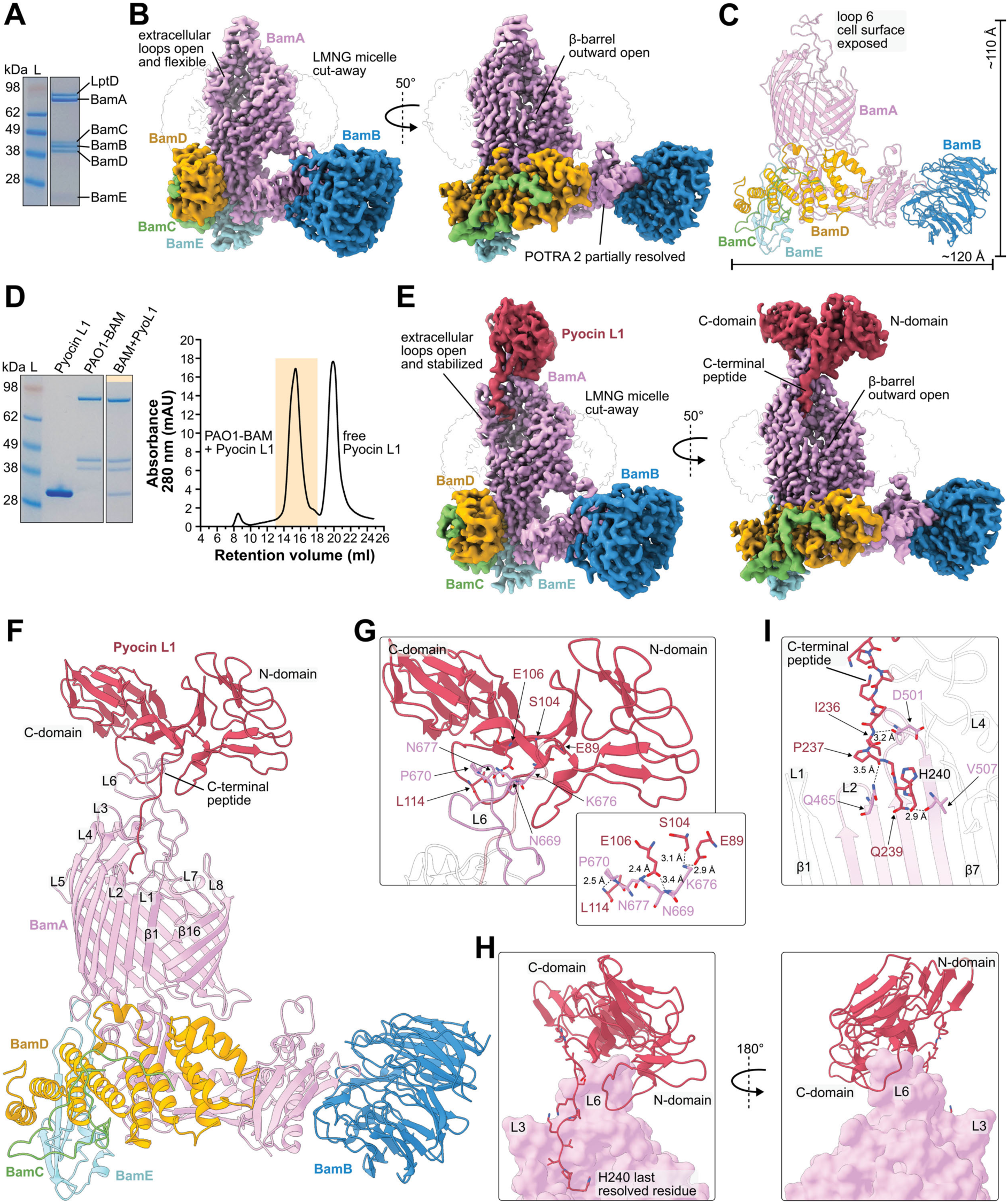
Pyocin L1 binds to loop 6 of BAM from P. aeruginosa PAO1. (A) A Coomassie-stained SDS-PAGE gel of the natively purified BAM complex from *P. aeruginosa*. (B) A 2.75 Å Coulomb potential map of *P. aeruginosa* BAM with a LMNG micelle cut-away. (C) A cartoon representation of the apo BAM structure (pdb: 9PXG). (D) A Coomassie-stained SDS-PAGE gel of overexpressed BAM from *P. aeruginosa* PAO1and Pyocin L1-His6 (left) and the corresponding size exclusion chromatogram (right). (E) A 2.84 Å Coulomb potential map of the BAM_PAO1_-Pyocin L1 in a pre-inhibition complex. (F) A cartoon representation of the PAO1-BAM-Pyocin L1 pre-inhibition complex structure (pdb: 9PXI). (G) A zoomed view of the binding interface between BamA loop 6 and Pyocin L1, shown as a cartoon with key electrostatic interactions shown as sticks. (H) Pyocin L1 (red cartoon) in complex with BamA (pink surface representation), showing interactions between Pyocin L1 and BamA loop 6, and deployment of the Pyocin L1 C-terminal peptide into the lumen of BamA. (I) A close-up view of the interface between the BamA β-strands 3-6 and the Pyocin L1 C-terminal peptide, in the interior of the BamA barrel.

To determine if BAM_PAO1_ and Pyocin L1 interact with high affinity, we combined these proteins at a ∼1:1 molar ratio and performed size exclusion chromatography demonstrating these species form a stable complex (**Fig. 2D**). We performed cryo-EM on this sample, resolving the structure of the BAM_PAO1_-Pyocin L1 complex to a global resolution of 2.84 Å (**Fig. S6, Table S1, Movie S2**). In comparison to the cryo-EM maps of BAM_PAO1_ alone, additional density is present in these maps corresponding to the two Pyocin L1 β-lectin domains, and part of the C-terminal peptide (**Fig. 2E**). This density allowed us to model the majority of the Pyocin L1 polypeptide chain (3-240 of 256 amino acids) in complex with BAM_PAO1_ (**Fig. 2F**). Consistent with Pyocin L1 susceptibility profiles, this structure demonstrated that Pyocin L1 interacts extensively with extracellular loop 6 of BamA, clasping this exposed loop between its two β-lectin domains (**Fig. 2G**).

In the x-ray structure of free Pyocin L1, the protein’s C-terminal peptide is bound in a cleft between the protein’s two β-lectin domains^12,16^. Strikingly, in the BAM_PAO1_-Pyocin L1 structure, the C-terminal peptide dissociates from the body of Pyocin L1 and is deployed into the lumen of the BamA β-barrel (**Fig. 2H**). The release of the peptide is driven by twisting of the Pyocin L1 β-lectin domains, likely resulting from loop 6 binding (**Fig. 2G**). The C-terminal peptide interacts with BamA extracellular loops 2, 3, and 4, stabilizing them in an outward open conformation (**Fig. 2F,I**). No density was observed for the final 16 C-terminal Pyocin L1 residues downstream of His-240, with this region positioned to occupy the lumen of the BamA barrel. To assess the dynamics of this region, we modelled it as an unstructured peptide within the lumen of the BamA β-barrel in an outer membrane lipid bilayer and performed 3 x 1µs molecular dynamics (MD) simulations. Consistent with the lack of density in our cryo-EM maps, the final 16 C-terminal Pyocin L1 residues were mobile during these simulations, adopting various conformations relative to the BamA β-barrel (**Movie S3**). The importance of the C-terminal peptide for Pyocin L1 activity suggests that its deployment into the BamA β-barrel is important for its cytotoxic mechanism, possibly through direct inhibition of the BAM complex. However, the lack of stable contacts between this region and BamA suggests that the BAM_PAO1_-Pyocin L1 structure does not represent a final inhibitory state. Unlike our activity assays, the Pyocin L1 construct used for this structure possessed a C-terminal His6-tag for affinity purification. C-terminally tagged Pyocin L1 exhibits comparable cytotoxic activity to the N-terminally tagged protein (**Fig. S1B**). However, the location of this tag may prevent the C-terminal peptide from fully associating with BamA in the detergent-solubilized BAM complex.

To determine the structure of BAM in complex with an N-terminally tagged L-pyocin, we combined N-terminally His6-tagged Pyocin L2 with the BAM complex from *P. aeruginosa* P28. Pyocin L2 and L1 share 86% sequence identity, and the sequence of their C-terminal peptides is identical, allowing for a direct comparison of this region of the structures. We confirmed that Pyocin L2 and BAM_P28_ form a stable complex via analytical-SEC and solved the structure of the complex via Cryo-EM to a global resolution of 3.18 Å (**Fig. 3A,B, Fig. S7, Table S1, Movie S4**). In this structure, Pyocin L2 binds BamA loop 6 in the same way as Pyocin L1 (**Fig. 3C**). However, sequence differences between Pyocin L1 and L2, and loop 6 of the respective BamA molecules, mean the molecular contacts that maintain this interface differ significantly (**Fig. 3D**). Like the BAM_PAO1_-Pyocin L1 structure, the C-terminal peptide of Pyocin L2 is deployed into the BamA lumen, where it stabilizes the extracellular loops 1-4 (**Fig. 3E**). Strikingly, in the BAM_P28_-Pyocin L2 structure, the Pyocin L2 C-terminal peptide is fully resolved, with the final 16 amino acids extending to bind to β-strand 1 of BamA (**Fig. 3C,F,G Fig. S8**). Amino acids 248–256 of Pyocin L2 bind to BamA β-strand 1 via β-augmentation, mimicking the interaction of an OMP terminal β-strand, with His253 occupying the same position as the conserved C-terminal aromatic residue of the OMP β-signal^19,32^. To assess the stability of the BAM_P28_-Pyocin L2 interaction, we performed 3 x 1µs MD simulations of the complex in an outer membrane bilayer. In these simulations and those of the pre-inhibition state, Pyocin L1 and L2 maintain stable interactions with loop 6 (5.6 ±1.6 and 5 ±1.5 H-bonds maintained over the simulations, respectively), while the C-terminal peptide of Pyocin L2 maintains a stable interaction with BamA β-strand 1. We observed no large-scale conformational changes, indicating the complex is stable in the membrane environment (**Fig. 2D**, **Fig. 3A**, **Movie S3 and S5**).

**Figure 3:**
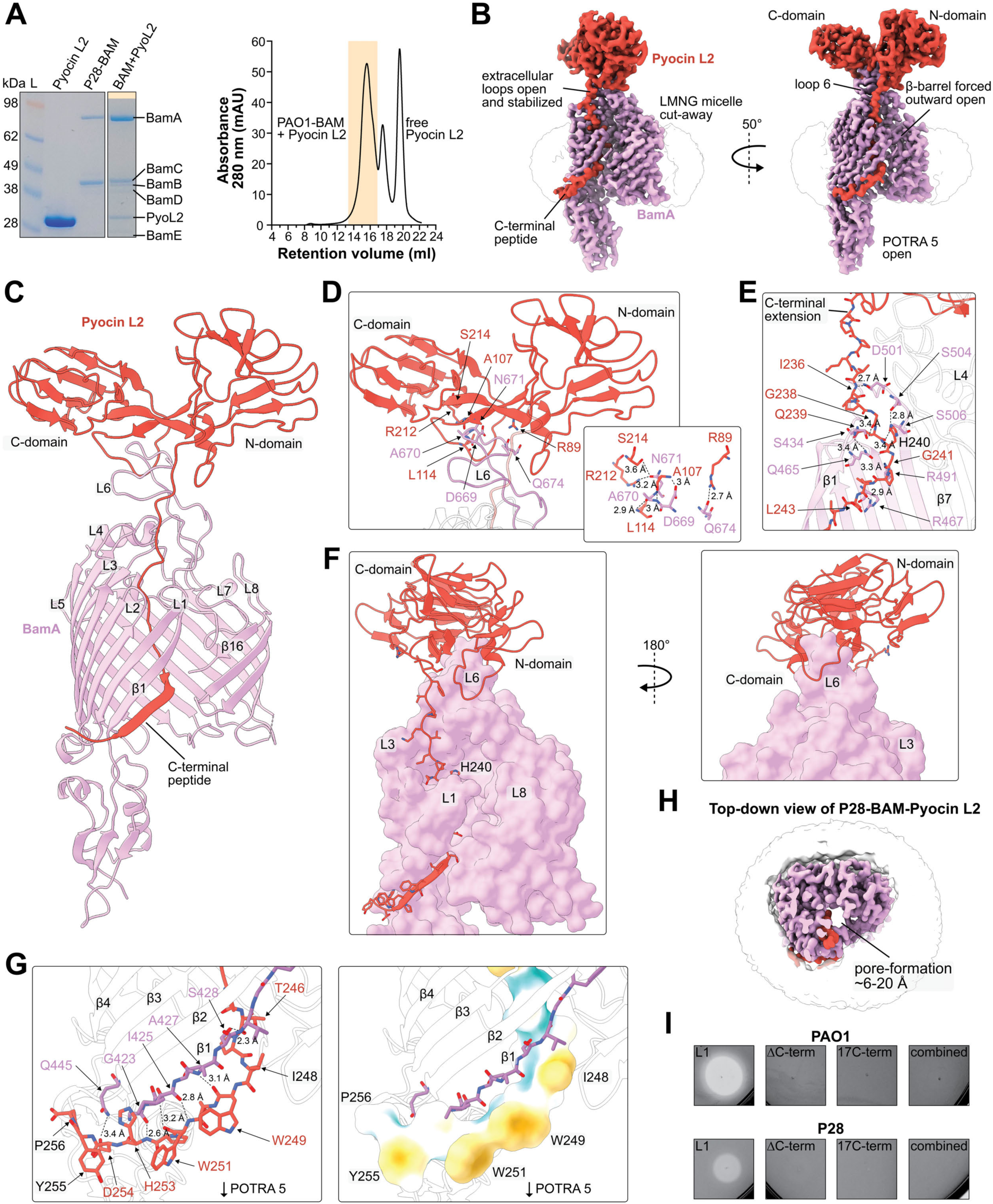
Pyocin L2 inhibits outer membrane protein assembly by blocking the β-barrel of BAM. (A) A Coomassie-stained SDS-PAGE gel of BAM from *P. aeruginosa* P28in complex with N-His6-Pyocin L2 (left) and the corresponding size exclusion chromatogram (right). (B) The 3.18 Å locally refined Coulomb potential map of BAM_P28_ (BamA-POTRA 5) in complex with His6-Pyocin L2 in a full inhibition complex. (C) Cartoon representation of the P28-BAM-Pyocin L2 full inhibition structure (pdb: 9PXJ). (D) A close-up view of the primary binding interface between BamA loop 6 and Pyocin L2, shown as a cartoon, with a stick representation of residues that form key electrostatic interactions. (E) A zoomed view of the interface between the BamA β-strands 3-6 and the Pyocin L2 C-terminal peptide, in the interior of the BamA barrel. (F) Pyocin L2 (red cartoon) in complex with BamA (pink surface representation), showing interactions between Pyocin L2 and BamA loop 6, and deployment of the Pyocin L2 C-terminal peptide into the lumen of BamA. (G) close-up of the binding interface of the Pyocin L2 C-terminal peptide at β-strand 1 of the BamA with electrostatic interactions (left) and hydrophobic surface representation (right). (H) A top-down view of the BAM_P28_-Pyocin L2 Coulomb potential maps with Pyocin L2 density outside the BamA barrel removed, showing Pyocin L2 binding to BamA creates an OM-spanning pore. (I) A plate-based inhibition assay of *P. aeruginosa* PAO1 and P28 with Pyocin L1, Pyocin L1ΔCterm (last 17 C-terminal residues deleted), a peptide of the 17 C-terminal residues of Pyocin L1, and the combination of Pyocin L1ΔC-term and 17aaC-term peptide.

The BAM complex adopts distinct extracellular loop-open (outward open) and loop-closed (inward open) states, where either the extracellular loops or POTRA 5 of BamA occlude its transmembrane β-barrel^29–33^. In the outward open state observed in our apo-BAM_PAO1_ and BAM_PAO1_-Pyocin L1 structures, BamA β-strand 1 is not accessible for peptide binding from the periplasm, due to contacts between the β-barrel N/C-termini and the presence of POTRA 5 (**Fig. 2C,F, Fig. S5**). Instead, functional BamA accepts nascent outer membrane proteins at β-strand 1 in the inward open conformation before transition to the open conformation for membrane insertion^21,32–34^. In the BAM_P28_-Pyocin L2 complex, BAM adopts an inward open conformation, where the BamA β-barrel is not occluded by POTRA 5 or the extracellular loops, leading to the formation of a continuous pore, ranging from ∼6-20 Å in diameter that has not been observed for other BamA-targeting antibiotics (**Fig. 3H**)^35–38^. A lack of interactions between POTRA 5 and the BamA β-barrel in this structure means that the POTRA domains and associated BamB-E subunits of this complex are mobile, only allowing modelling of Pyocin L2 and the BamA β-barrel with POTRA 5 (**Fig. 3C**).

The interaction between the C-terminal peptide of Pyocin L1 and BamA β-strand 1 is similar to the binding of BamA bamabactins family peptides (darobactin, dynobactin, and xenorceptide)^35–38^. Bamabactins are a recently identified family of macrocyclic peptide antibiotics that inhibit the BAM complex by competitively blocking β-strand 1 (**Fig. S8**). Based on this similarity, it is highly probable that L-type pyocins are also competitive inhibitors of OMP binding by the BAM complex. To test whether the C-terminal peptide in isolation is cytotoxic to *P. aeruginosa* cells, we synthesised the 17 C-terminal Pyocin L1 residue as a peptide and tested it against *P. aeruginosa* PAO1 and P28 in plate-based growth inhibition assays, where this peptide was not toxic even at a high concentration (10 mM) (**Fig. 3I**). We then exposed both strains to a Pyocin L1 C-terminal deletion construct (Pyocin L1_ΔCterm_) showing that the β-lectin domains are not active without the C-terminal peptide, even though they likely retain the capacity to bind BamA loop 6. We then applied both the isolated C-terminal peptide and Pyocin L1_ΔCterm_ concurrently, which also did not inhibit the growth of these strains (**Fig. 3I**). This indicates that while the Pyocin L1 C-terminal peptide is critical for toxicity, it requires the β-lectin domains for its delivery to the lumen of BamA.

### L-type pyocins inhibit the BAM complex via a multistage process

Our structural analysis shows that L-type pyocins bind to loop 6 of BamA and deploy their C-terminal peptide into the BamA lumen to inhibit the BAM complex by binding β-strand 1 (**Fig. 4A-C**). Entry of the C-terminal peptide is only possible when BamA is in the outward open conformation (**Fig. 4D,E**). However, β-strand 1 of BamA is only accessible in the inward open conformation, where entry of the C-terminal peptide into BamA is blocked. This mutual exclusivity indicates a two-step inhibition mechanism, where initial binding and C-terminal peptide deployment occurs in the outward open state, creating a primed or pre-inhibited BAM complex. Full inhibition then requires transition to the inward open state, which exposes β-strand 1 (**Movie S6**). Notably, our MD simulations of the BAM_PA01_-Pyocin L1 complex did not show spontaneous transition from the outward open to inward open state (**Movies S3** and **S5**), this conformational change may result from the delivery of nascent outer membrane proteins for assembly rather than being triggered solely by interactions with Pyocin L1 (**Fig. 4F**)^21,32–34,39^. Our MD simulations of the fully inhibited state provide context for the positioning of the complex in the outer membrane bilayer (**Fig. 4G,H**). The position of Pyocin L1 is compatible with binding D-rhamnose moieties of CPA via its C-terminal β-lectin domain, which may further stabilise the complex^12^. Furthermore, the last three residues of the C-terminal peptide are flexible and extend into the periplasm, which may allow for recognition of L-type pyocin intoxication by the bacterial cell.

**Figure 4:**
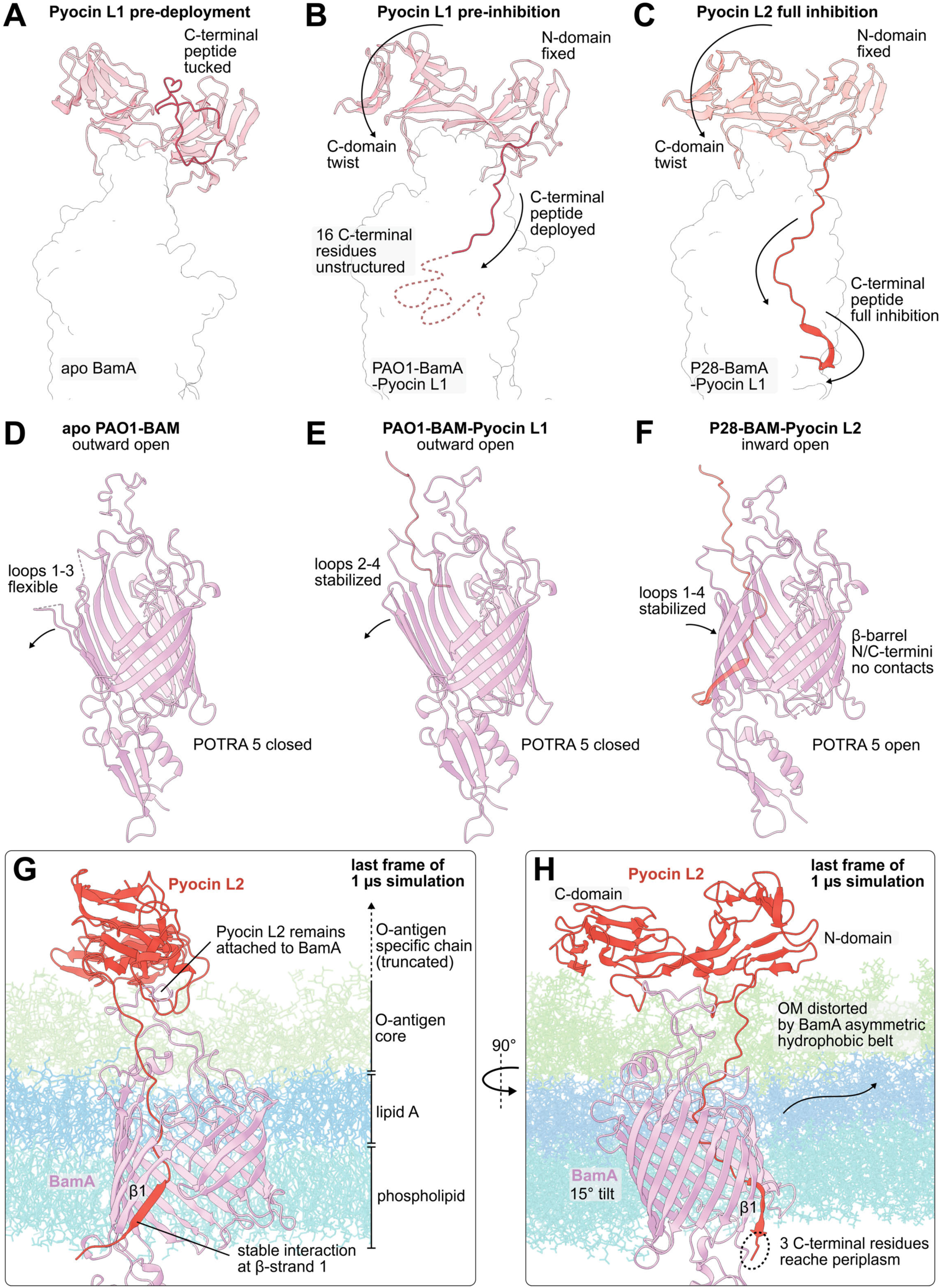
L-type pyocins and BAM undergo extensive conformational changes to form a stable complex. (A) Side-view of BamA (white surface) with unbound Pyocin L1docked to loop 6 by alignment with the N-domain of bound Pyocin L1. The C-terminal peptide (red) of Pyocin L1 is tucked between the N- and C-domains. (B) The pre-inhibition state of PAO1-BAM-Pyocin L1 (pdb: 9PXI) shows a twisting of the N/C-domains relative to each other. The C-terminal peptide is deployed into the BamA lumen, where about one-third binds to BamA and the remaining two-thirds remain flexible (dotted line). (C) The full inhibition state of P28-BAM-Pyocin L2 (pdb: 9PXJ) with the fully resolved C-terminal peptide binding β-strand 1of the BamA. (D) Side-view of apo PAO1-BAM (pdb: 9PXG) with POTRA 5 blocking access to the outward open BamA lumen and flexible loops 1-3. (E) Side-view of the PAO1-BAM-Pyocin L1 pre-inhibition complex with POTRA 5 blocking access to the BamA lumen with stabilized loops 2-4. (F) Side-view of the P28-BAM-Pyocin L2 full inhibition complex with POTRA 5 open, allowing access to the BamA lumen from the periplasmic side, and loops 1-4 stabilized. The β-barrel assumes an incomplete inward open conformation, as it cannot form contacts with the C-terminal side due to the presence of the Pyocin L2 C-terminal peptide. (G) MD simulation snapshot (1 µs frame) of the P28-BAM-Pyocin L2 full inhibition complex (POTRA 1-4 and BamB-E excluded) inside an outer membrane bilayer (O-antigen specific chain excluded). Pyocin L2 interactions with BamA loop 6 and β-strand 1 remain stable (see Movie S5). (H) The P28-BAM-Pyocin L2 simulation (90° rotated) shows that the N- and C-domains are in proximity to O-antigen (truncated). The last 3 residues of the C-terminal peptide are flexible and located in the periplasm, and BamA is tilted by ∼15°, causing minor distortion of the bilayer.

### Outer membrane homeostasis is critical for Pyocin L1 tolerance

To better understand the consequences of Pyocin L1 intoxication on *P. aeruginosa* cells, we used TraDIS to identify genes in *P. aeruginosa* PAO1 that are important for susceptibility or tolerance to Pyocin L1^40,41^. A library of *P. aeruginosa* PAO1 cells containing >450,000 random genomic transposon insertions, equivalent to an insertion every 15 bp, was cultured in the presence and absence of 14 nM (0.4 μg/mL) Pyocin L1, before the genomes of both populations were sequenced, and the location and abundance of transposon insertions were analyzed. Transposon insertions in genes that contribute to Pyocin L1 tolerance will show reduced abundance after Pyocin L1 treatment, while transposon insertions in genes involved in Pyocin L1 sensitivity will show increased abundance (**Table S2** and **Supplementary Dataset S2**). The majority of the genes that contribute to Pyocin L1 sensitivity are associated with the biosynthesis of the LPS CPA, which is consistent with Pyocin L1 binding non-essential CPA as a cell surface receptor (**Fig. 5A,C**)^12^. Other Pyocin L1 sensitizing genes include the cell stress-sensing two-component regulator *gacA/gacS*, and a limited number of genes associated with metabolic processes (**Fig. 5C, Fig. S9A,B, Table S2**)^42^. Interestingly, the genes PA5001 and PA5004, which encode putative glycosyltransferases that are predicted to be involved in LPS core biosynthesis, contribute to Pyocin L1 sensitivity, while PA5002 and PA5003 from the same operon, which encode proteins of unknown function, contribute to Pyocin L1 tolerance (**Fig. 5B,C,F**)^43^.

**Figure 5:**
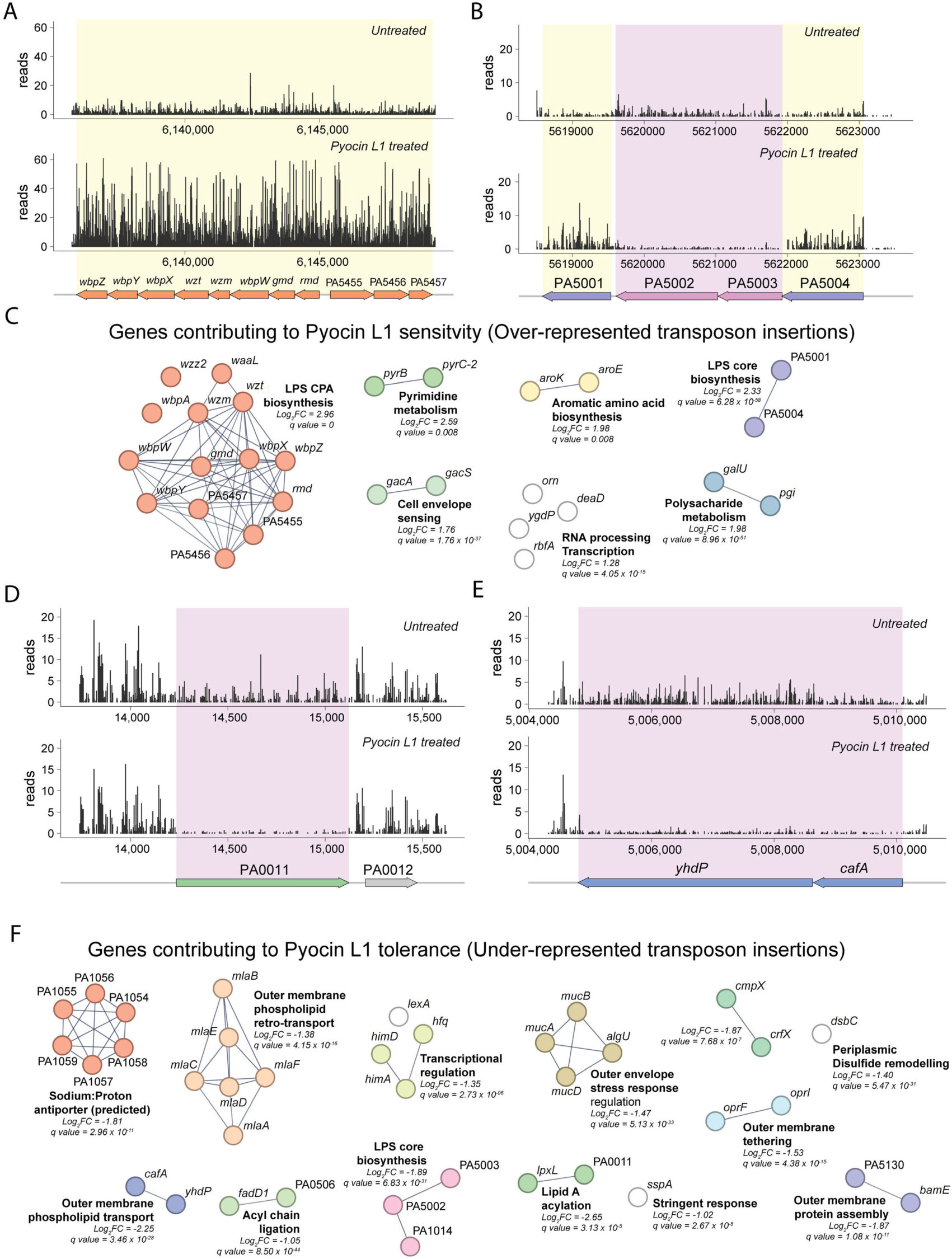
Identification of genes important for Pyocin L1 sensitivity and tolerance using TraDIS. (A) A TraDIS insertion plot of a region of the *P. aeruginosa* PAO1 genome, showing that Pyocin L1 treatment leads to the enrichment of transposon insertions in the gene cluster responsible for CPA O-antigen biosynthesis. (B) A TraDIS insertion plot showing that Pyocin L1 treatment causes a mixed insertion enrichment profile for the region of the *P. aeruginosa* PAO1 genome containing PA5001-PA5004. (C) STRING functional clustering analysis of genes with increased transposon insertion frequency after Pyocin L1 treatment, indicating a role in susceptibility. (D) and (E) TraDIS insertion plots showing that insertions in PA0011, *yhdP*, and *cafA* are under-represented after Pyocin L1 treatment, indicating they are important for Pyocin L1 tolerance. (F) STRING functional clustering analysis of genes with decreased transposon insertion frequency after Pyocin L1 treatment, indicating a role in tolerance. The frequency of transposon insertions is indicated by the size of the black ‘reads’ bar at that position. Regions with a higher frequency of transposon insertion in the Pyocin L1-treated cells are highlighted in yellow, while those with a lower frequency of insertion are highlighted in purple. Genes encoded in these regions are shown below the insertion plots.

Multiple genes encoding components of systems responsible for outer membrane assembly and homeostasis are important for Pyocin L1 tolerance (**Fig. 5D,E,F, Fig. S9C-F**, **Table S2**). These include the BAM complex component BamE, further reinforcing the BAM complex as the target for L-type pyocins (**Fig. S9D**). Both the AsmA-family protein YhdP, which plays an important role in transporting phospholipids to the outer membrane, and the MlaA-F system, which performs retrograde phospholipid transport critical for maintaining the asymmetry of the outer membrane bilayer, were important for Pyocin L1 tolerance (**Fig. 5E,F, Fig. S9C,F**)^44–46^. This indicates that disruption of both outer membrane phospholipid delivery and partitioning enhances *P. aeruginosa*’s susceptibility to Pyocin L1. In addition, OprF and OprL (equivalent to OmpA and Lpp from *Escherichia coli*) contribute to Pyocin L1 tolerance. These proteins tether the outer membrane to the peptidoglycan cell wall, helping to maintain its stability^47–49^. DsbC, which is required for proper disulphide bond formation in outer envelope proteins, also contributes to Pyocin L1 tolerance, as do the outer envelope stress response regulators MucA,B,D and AlgU (**Fig. S9F, Table S2**)^50–52^. Taken together, the prevalence of outer envelope proteins that contribute to both outer membrane maintenance and Pyocin L1 tolerance indicates that Pyocin L1 treatment places considerable stress on outer membrane homeostasis systems. This is consistent with our structural data and supports a mechanism of action for L-type pyocins that involves BAM complex inhibition, leading to outer membrane disruption.

### *P. aeruginosa* PAO1 responds rapidly to Pyocin L1 intoxication

To understand the cellular consequences of L-type pyocin intoxication, we performed dual transcriptomic (RNA sequencing) and proteomic profiling of *P. aeruginosa* PAO1 following Pyocin L1 exposure (**Fig. 6**). Transcriptomic responses were examined at 15- and 120 minutes post-treatment (∼1/2 and 4 cell divisions, respectively^53^), while proteomic changes were assessed at 300 minutes (∼10 cell divisions) to consider slower changes in the proteome and to capture later-stage consequences of Pyocin L1 intoxication. Transcriptional profiling 15 minutes post-treatment reveals a rapid and multifaceted response involving the differential expression of 771 genes (377 upregulated, 394 downregulated; Log_2_FC > 0.5; −log_10_FDR > 1.3) (**Fig. 6A, Supplementary Dataset S3**). These changes reflect both direct physiological disruptions caused by Pyocin L1 and its recognition as a signal of microbial antagonism. A hallmark of the early response is the strong downregulation of genes involved in translation (118 genes) and protein export (*secA,B,D,E,Y, yidC*). This is likely a result of cells sensing inhibition of the BAM complex by Pyocin L1, which stalls the insertion of outer membrane proteins (OMPs), leading to the repression of protein synthesis to mitigate cell envelope disruption. Concurrently, *P. aeruginosa* reprograms its metabolism, upregulating at least 33 genes involved in the catabolism of branched-chain amino acids and fatty acids, consistent with a shift from growth to non-replicative persistence (**Fig. 6B**). Selective changes in OMP gene expression occur, supporting a strategy to address envelope stress and community-level competition. A limited number of TonB-dependent transporters (TBDTs) are repressed, while some major porins (e.g., *oprB, oprD*) and proteins associated with cell envelope integrity (e.g. *oprL, oprF*) are upregulated, suggesting remodelling of the outer membrane to reduce vulnerability to competitor-produced bacteriocins, which often target TBDTs for cell entry^54^, and to maintain outer membrane homeostasis^55^. Notably, genes associated with the type VI secretion system (T6SS, ≥24 genes) are strongly induced, suggesting a response to Pyocin L1 as a hostile signal, triggering this contact-dependent defence mechanism^56,57^. This is accompanied by activation of phenazine biosynthesis, quorum sensing, and biofilm-associated genes, indicating the mounting of a coordinated, community-level defence (**Fig. 6B, Supplementary dataset S3**).

**Figure 6:**
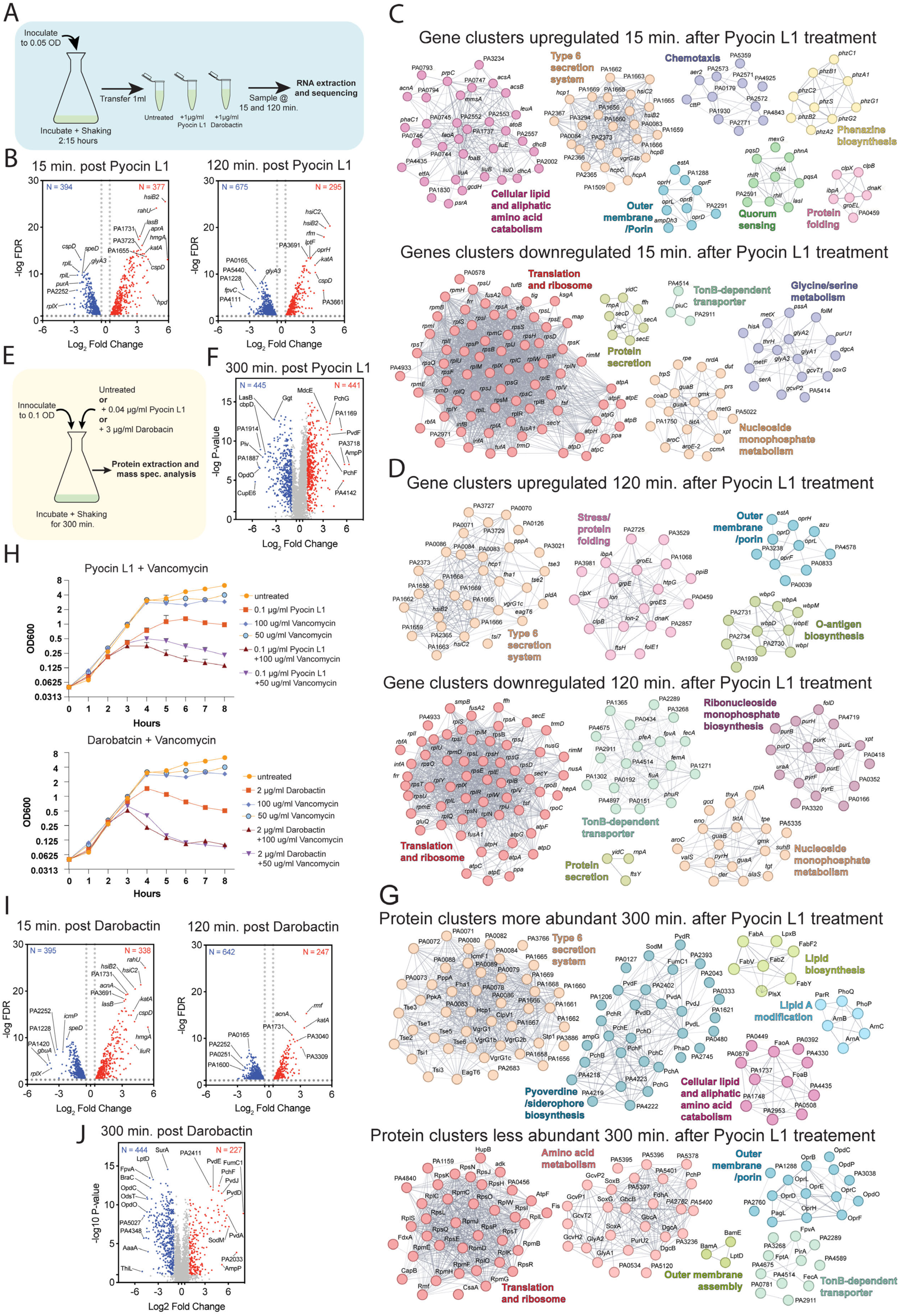
Transcriptomic and proteomic analysis indicate a rapid and targeted cellular response to Pyocin L1 and darobactin treatment. (A) A schematic of *P. aeruginosa* PAO1 treatment and sample preparation for transcriptomic analysis of the response to Pyocin L1 and darobactin. (B) Volcano plots showing genes differentially expressed by *P. aeruginosa* PAO1 in response to Pyocin L1 at 15- and 120-minutes post-treatment (Log_2_FC > ±0.5; −log_10_FDR > 1.3). (C) and (D) STRING functional clustering analysis of genes with increased and decreased expression 15 and 120 minutes after Pyocin L1 treatment, respectively. (E) A schematic of *P. aeruginosa* PAO1 treatment and sample preparation for proteomic analysis of the response to Pyocin L1 and darobactin. (F) A Volcano plot showing proteins with altered abundance in *P. aeruginosa* PAO1 in response to Pyocin L1 at 300 minutes post-treatment, proteins with increased and decreased abundance (Log_2_FC >1; −Log_10_Pvalue >1) are coloured red and blue, respectively. (G) STRING functional clustering analysis of proteins with increased and decreased abundance 300 minutes after Pyocin L1 treatment. (H) Liquid culture growth curves of *P. aeruginosa* PAO1 cells treated with Pyocin L1 or darobactin, in combination with Vancomycin. (I) Volcano plots showing genes significantly differentially expressed by *P. aeruginosa* PAO1 in response to darobactin at 15- and 120-minutes post-treatment (Log_2_FC > ±0.5; −log_10_FDR > 1.3). (J) A Volcano plot showing proteins with altered abundance in *P. aeruginosa* PAO1 in response to darobactin at 300 minutes post-treatment, coloured as in panel F.

By 120 minutes, the transcriptome shifts toward managing accumulating protein folding and envelope stress, with robust upregulation of chaperone systems (including *lon1/2, groESL, clpBX, dnaK*), likely a consequence of misfolded OMPs accumulating due to BAM inhibition (**Fig. 6A,C, Supplementary dataset S3**). Interestingly, significant upregulation of periplasmic chaperones (*surA, degP, skp,* etc.) was not observed, indicating that folding stress is not directly addressed in the periplasm of the cell envelope. At least 65 secretion signal peptide-encoding genes, including outer membrane porins, are upregulated, likely in an indirect attempt to restore outer membrane function and integrity. Conversely, there is reinforced repression of 18 TBDTs. Strong translational repression and some Sec system suppression (e.g., *secY*, *secE*, *yidC*) persist, showing a continued adaptive response to envelope stress. T6SS gene induction remained strong, but community-level signalling (quorum sensing, phenazine production) is dampened, suggesting a shift from cooperation to self-preservation (**Fig. A,C, Supplementary dataset S3**).

Proteomic analysis at 300 minutes post-treatment reveals significant changes in protein abundance during Pyocin L1 intoxication, with 986 proteins (441 increased or 445 decreased; Log_2_FC >1; −Log_10_Pvalue >1) showing altered abundance compared to untreated cells (**Fig. 6D, Supplementary dataset S4**). These changes reflect both a sustained stress response and systemic disruption of protein homeostasis and envelope integrity. Consistent with transcriptomic data, T6SS components were more abundant, and ribosomal proteins and translation factors were reduced, confirming prolonged growth suppression. Strikingly, 167 signal peptide-containing proteins were reduced in abundance, of which 37 were OMPs, including 11 TBDTs and 12 porins (**Fig. 6D,E**). In contrast to the transcriptomic data, OprF and OprL, which are critical for outer membrane tethering, were reduced in abundance, indicating an impaired ability to maintain outer envelope structure. Notably, the BAM components BamA and BamE, and the LPS insertase LptD, were reduced in abundance, indicating an impaired ability to incorporate OMPs and LPS into the outer membrane. Fatty acid biosynthesis-related proteins increased in abundance, possibly to compensate for failure to insert OMPs and LPS, by reinforcing the outer membrane with phospholipids in place of these key components. Despite transcriptional induction of chaperones, few were elevated at the protein level, suggesting an inability to effectively address folding stress. However, components of the Arn lipid A modification system (ArnA, ArnB, ArnC) were upregulated, indicating activation of resistance mechanisms against cationic antimicrobial peptides, consistent with a perceived breach in envelope integrity. Finally, a greatly increased abundance of pyoverdine biosynthesis and export proteins was observed, likely reflecting iron starvation, possibly due to TBDT repression and membrane transport dysfunction (**Fig. 6E, Supplementary dataset S4**).

Based on our omics data, which indicates that L-type pyocins compromise outer membrane integrity and the pore formed by the BAM_P28_-Pyocin L2 complex, we hypothesised that Pyocin L1 may sensitize *P. aeruginosa* to antibiotics normally too large to cross the outer membrane (**Fig. 3H**). To test this, we co-treated *P. aeruginosa* PAO1 with Vancomycin, at concentrations that are not normally cytotoxic to this bacterium, and either Pyocin L1 or darobactin. We saw a more rapid and pronounced decline in the optical density of cultures treated with these antibiotic combinations, indicating that both Pyocin L1 and darobactin compromise the outer membrane, allowing Vancomycin to target the cell wall in the periplasm (**Fig. 6H**). However, this effect only becomes pronounced 4 hours after antibiotic treatment, making it unclear whether it is related specifically to the pore formed by the BAM-antibiotic complex or a more general disruption of the outer membrane.

### The response of *P. aeruginosa* to Pyocin L1 and darobactin diverges over time

Given similar mechanisms of action of L-type pyocins and the bamabactins^7,35^, we compared the transcriptional and proteomic response of *P. aeruginosa* PAO1 to Pyocin L1 with that of darobactin. At 15 minutes post-treatment, the transcriptional responses to Pyocin L1 and darobactin were similar, with over half of the significantly upregulated and downregulated genes shared between the two conditions (**Fig. 6I, Fig. S10A, Supplementary dataset S3**). Both agents induced a common set of responses, including upregulation of genes involved in T6SS, catabolism of fatty acids and aliphatic amino acids, phenazine biosynthesis, and quorum sensing. Simultaneously, both treatments triggered broad downregulation of translation, ribosome biogenesis, protein export, and nucleotide biosynthesis, reflecting a rapid shift away from growth-related processes (**Fig. 6C, Fig. S10C**). Despite these similarities, there were notable differences. Pyocin L1 triggered stronger T6SS induction (24 vs 18 genes upregulated), increased expression of lipid oxidation genes, and modest upregulation of specific outer membrane porins. In contrast, darobactin treatment led to the induction of genes involved in oxidative phosphorylation, including components of respiratory complexes I and IV. Pyocin L1 also provoked a more pronounced repression of ribosomal genes, whereas darobactin more strongly downregulated genes associated with fatty acid and carboxylic acid metabolism. These distinctions suggest that while both antibiotics disrupt envelope homeostasis through BAM inhibition, they differ in how *P. aeruginosa* responds to the resulting stress (**Fig. S10C, Supplementary dataset S3**).

At 120 minutes post-treatment, the transcriptomic responses remained broadly similar, with over half of the upregulated genes and a substantial minority of downregulated genes shared between conditions (**Fig. 6I, Fig. S10B, Supplementary dataset S3**). Shared transcriptional programs included upregulation of cytosolic protein folding chaperones such as *groES, groEL, clpB, clpX, and dnaK*, as well as genes associated with iron metabolism, heme biosynthesis, and central carbon metabolism. Translation remained repressed under both treatments, although the effect was more pronounced in Pyocin L1-treated cells. A shared envelope stress signature was also evident, including strong downregulation of TonB-dependent transporters (TBDTs) and upregulation of key structural proteins such as OprF and OprL. However, treatment-specific differences became more pronounced over time. While Pyocin L1 sustained strong induction of the T6SS, this response was absent in darobactin-treated cells at 120 minutes, with eleven T6SS-related genes downregulated. The induction of oxidative phosphorylation genes observed in darobactin-treated cells was maintained, whereas no such upregulation was observed following Pyocin L1 exposure. These distinctions point to a divergence in long-term adaptation strategies: Pyocin L1 triggers persistent interbacterial aggression, while darobactin elicits a more inwardly focused metabolic realignment (**Fig. 6D,G, Fig. S10D, Supplementary dataset S3**).

Proteomic analysis at 300 minutes post-treatment revealed that the overlap in responses between Pyocin L1 and darobactin diminished at the protein level (**Fig. 6J, Fig. S10E,F, Supplementary data S3**). Only a minority of proteins with altered abundance were shared between the two treatments. Nonetheless, several core stress responses remained conserved, which is consistent with a shared cytotoxic mechanism. Both treatments led to a greatly increased abundance of proteins involved in pyoverdine biosynthesis. Lipid metabolism was similarly affected, with both treatments showing upregulation of fatty acid biosynthesis enzymes, including those involved in lipid A production. A prominent shared feature was the reduction in outer membrane protein abundance, including porins and TBDTs, indicative of impaired BAM function and outer membrane assembly. However, differences were again evident. Pyocin L1-treated cells showed a clear reduction in ribosomal and translation-associated proteins, consistent with transcriptional repression observed at 120 minutes post-treatment. In contrast, darobactin-treated cells did not display the same proteomic signature of translation inhibition, likely reflecting the weaker repression of these genes at the 120-minute timepoint. Moreover, darobactin-treated cells exhibited additional reductions in proteins involved in fatty acid metabolism and iron handling, including BfrB, SodB, and CcoN2, suggesting a more pronounced impact on metabolic homeostasis (**Fig. 6G,J, Fig. S10E,F, Supplementary dataset S4**). MD simulations show that the 3 C-terminal residues of Pyocin L2 reach the periplasm (**Fig. 4H**). It is plausible that *P. aeruginosa* has evolved an L-type pyocin recognition mechanism that dictates a different stress response compared to darobactin exposure, which is further supported by the apparent co-evolution of BamA loop 6 and L-type pyocins.

### Pyocin L1 treatment leads to cell envelope degradation

While *P. aeruginosa* rapidly responds to Pyocin L1 by radically remodelling its transcriptome and proteome, the cells ultimately succumb to a process likely driven by an extreme loss of outer envelope proteins. To assess the consequences of this process on cell morphology, we performed cryo-ET on Pyocin L1 and darobactin-treated cells. We explored different timepoints post-treatment, settling on 420 minutes. At earlier time points (e.g. 2 hours post-treatment), gross morphological changes were rarely evident (**Fig. 1A, Fig. S11**). Cryo-ET imaging of untreated cells and 3D segmentation exhibited typical cell morphology for early stationary phase, with an intact cell envelope, an abundance of ribosomes, a polar flagella and chemotaxis arrays (**Fig. 7A-E, Fig. S12A, Movie S7**). In contrast, Pyocin L1-treated cells exhibited significant loss of outer membrane integrity, characterized by large holes/gaps, or delamination leading to outer membrane blebbing and particle-filled vesicle release (**Fig. 7F-K, Fig. S12B, Fig. S13, Movie S8**). The phenotype of darobactin-treated cells was similar, with subtle differences, including multilayered intracellular vesicle formation and rupture of the outer and inner membranes, which was not observed in Pyocin L1-treated cells (**Fig. 7L-P, Fig. S12C, Fig. S13, Movie S9**). Consistent with the omics data, the ribosome content of Pyocin L1-treated cells was substantially lower than wild-type cells. Ribosome numbers were also reduced in darobactin-treated cells, but to a lesser extent than observed in Pyocin L1-treated cells. Also, consistent with our omics data, T6SSs were significantly more abundant in Pyocin L1-treated cells compared to either darobactin-treated or untreated cells, confirming that specific recognition of Pyocin L1 triggers this anti-competition response. (**Fig. 7, Fig. S12, Fig. S13, Movies S5-7**).

**Figure 7:**
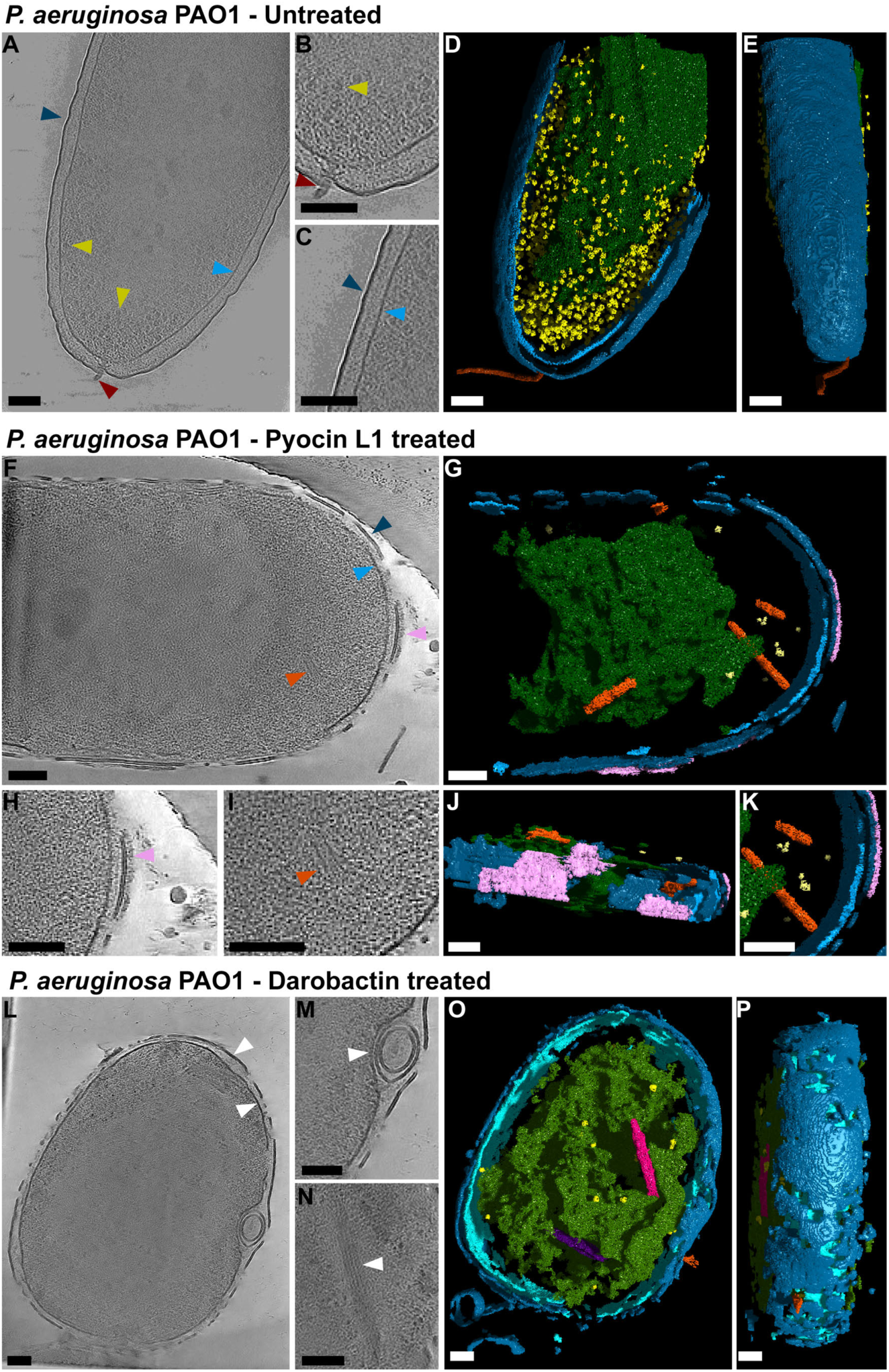
Analysis of antimicrobial treatment of *P. aeruginosa* cells by cryo-ET. (A) 2D-slice view (4 slices) through a 3D reconstructed tomogram of an untreated *P. aeruginosa* cell. (B, C) Enlarged 2D slice views of the cell depicted in (A). (D) Segmentation of an untreated *P. aeruginosa* cell is shown in (A) top down and rotated 90 degrees along the x-axis (E). (F) 2D-slice view (4 slices) through a 3D reconstructed tomogram of a *P. aeruginosa* cell treated for 7 hours with Pyocin antimicrobial. (G) Segmentation of the 3D tomogram shown in (F). (H,I) enlarged 2D slice views of the same cell shown in (F). (J) Side view of segmentation shown in (G) rotated 90 degrees about the y-axis. (K) Enlarged view of the segmentation shown in (G). (L) 2D-slice view of a *P. aeruginosa* cell treated with darobactin for 7 hours. (M,N) Enlarged views of the same volume are shown in (L). (O) Segmentation of the 3D volume shown in (L) from a top-down view and rotated 90 degrees along the x-axis. (P) Cellular features, highlighted by arrows or in segmentation, are coloured as follows: the outer membrane (dark blue), inner membrane (light blue), ribosomes (yellow), flagella (red), T6SS (orange), diffuse outer layer (pink), cytoplasmic filament 1 (pink), cytoplasmic filament 2 (purple). Scale bars represent 100 nm.

## Discussion

Our work reveals that L-type pyocins are potent, evolutionarily tailored protein antibiotics that kill *P. aeruginosa* by targeting the BAM complex, an essential system for the insertion of outer membrane proteins^32^. We demonstrate that L-type pyocins bind to loop 6 of BamA and insert their C-terminal peptide into BamA binding β-strand 1 and arresting BAM activity, which inhibits OMP assembly and compromises outer membrane homeostasis. This mechanism is remarkably like that of the bamabactin family peptides and represents a striking example of convergent evolution. Both families of molecules exploit the same vulnerability in Gram-negative bacterial physiology, reinforcing the validity of the BAM complex as an antibiotic target^7,13,35,37,58^. Crucially, this mechanism allows L-type pyocins to kill *P. aeruginosa* without crossing the outer membrane, abrogating one of the most formidable permeability barriers in biology, and making them immune to efflux pump-mediated resistance^59,60^. The extensive interactions between L-type pyocins and BamA account for their narrow spectrum of activity and underlie their remarkable potency, more than 50-fold greater than that of bamabactins^17,35–37^. This enhanced activity likely arises from a combination of high-affinity binding of BamA loop 6 and targeted delivery of the inhibitory peptide to the BamA lumen. Interestingly, while the majority of the BamA sequence is conserved in *P. aeruginosa* strains, loop 6 appears to under substantial selection pressure most likely attributable to targeting by L-type pyocins.

Beyond establishing the structural and mechanistic basis for L-type pyocin toxicity, we also comprehensively profile the cellular response to L-type pyocin exposure and BAM complex inhibition more generally. Transcriptomic analysis reveals that *P. aeruginosa* rapidly senses and responds to Pyocin L1, dramatically reprogramming gene expression in an attempt to mitigate the threat. This response is multifaceted: translation is sharply suppressed, which pre-emptively addresses the burden of mislocalized outer membrane proteins, and metabolic pathways are restructured to prepare for a shift from growth to survival. Upregulation of genes associated with the T6SS, biofilm formation, and quorum sensing indicates that the bacterium recognizes Pyocin L1 as a weapon in interbacterial competition, prompting reciprocal aggression and community-level defense. Two hours after Pyocin L1 treatment, before an observable decline in growth rate or overt morphological damage, the transcriptional response shifts. There is a marked increase in the expression of genes associated with protein folding and outer envelope stress, hallmarks of a failure to compensate for BAM inhibition^61,62^.

In the longer term, BAM inhibition by L-type pyocins triggers a collapse in outer membrane integrity. Our TraDIS data highlights the critical importance of cell envelope maintenance pathways, including LPS biosynthesis, outer membrane tethering and phospholipid transport and partitioning, in conferring greater tolerance to L-type pyocin attack. This is mirrored at the proteomic level with global reductions in outer membrane proteins, including essential BAM subunits and porins. Comparison of the cellular response to L-type pyocins and darobactin reveals both shared and distinct stress signatures. While both agents elicit a core BAM-inhibition response, their downstream effects diverge. L-type pyocins trigger sustained interbacterial aggression via T6SS^56,57^, whereas darobactin induces a metabolic shift indicative of oxidative stress adaptation. These differences also likely reflect the ecological origin of the molecules; L-type pyocins are recognized by *P. aeruginosa* as intraspecies weapons, while darobactin is produced by a phylogenetically distant organism that broadly targets BAM from many bacteria^14,35^.

Cryo-ET reveals the terminal consequences of L-type pyocin intoxication: structural disintegration of the outer membrane, loss of ribosomes, and deployment of the T6SS. These features illustrate the broad physiological collapse and response triggered by BAM inhibition, despite a concerted response to mitigate this damage. The morphological similarities between pyocin- and darobactin-treated cells further reinforce that, despite compensatory stress responses, BAM inhibition ultimately leads to lethal outer membrane disruption. The proximal cause of this disintegration is halted outer membrane protein insertion; however, the consequences are far-reaching. These include the loss of LptD-dependent LPS insertion, outer membrane tethering, and permeability via porins and TBDTs. The dramatic increase in siderophore biosynthesis at later time points may reflect iron starvation due to reduced siderophore import in cells before outer membrane disruption. The outer membrane’s biophysical properties are also likely altered; phospholipid content likely increases and compensates for the dearth of proteins and LPS, decreasing membrane rigidity and increasing fluidity, which may explain the extreme blebbing and vesicle formation observed in L-type pyocin and darobactin-treated cells.

Together, our findings establish L-type pyocins as precision-guided inhibitors of the BAM complex, with both therapeutic potential and ecological significance. These results broaden our understanding of how bacteria target essential cellular machineries and open new avenues for engineering proteins to kill Gram-negative pathogens by targeting the BAM complex without entering the bacterial cell.

## Methods

### Bacterial strains and culturing conditions

For cloning, *E. coli* DH5α was grown in Lysogeny Broth (LB) (10 g/l Tryptone, 5 g/l Yeast extract, 5 g/l NaCl, 15 g/l agar) and maintained with antibiotics where appropriate. Liquid cultures were incubated at 37°C with orbital shaking at 180 rpm. All *P. aeruginosa* strains could be grown on LB plates and in LB broth at 37°C and 180-200 rpm shaking. The clinical isolates grew with a range of bright green to yellow pigments, and some tended to clump together but could be separated by resuspension with a 1 mL pipette.

For the overexpression of L-type pyocins, *E. coli* BL21(DE3)C41 was transformed with expression plasmids (**Table S3**). After 1 h of recovery, a starter culture was inoculated with 90% of the transformed *E. coli* and grown in LB broth with 50 µg/mL Kanamycin (pET28a, pET29b) or 100 µg/mL Ampicillin (pET23a) at 37°C overnight at 180 rpm on a rotary shaker. The remaining 10% of the transformation was plated on LB agar with the appropriate antibiotic. On the next day, the main cultures were inoculated to a start OD_600_ of 0.05 and grown until OD_600_ of 0.8-1 in Terrific Broth (TB) (12 g/l Tryptone, 24 g/l Yeast extract, 12.3 g/l K_2_HPO_4_, 2.52 g/l KH_2_PO_4_) at 37°C at 200 rpm. The cultures were cooled to 4°C and induced with 0.3 mM IPTG for expression for ∼22 h at 20°C at 200 rpm.

For BAM overexpression, *E. coli* BL21(DE3)C41 or *P. aeruginosa* PAO1 were transformed with expression plasmids and spread on LB plates with 100 µg/mL Trimethoprim and grown overnight at 37°C. The next day, a starter culture was inoculated with transformed cells and grown in LB broth with 100 µg/mL Trimethoprim at 37°C overnight at 180 rpm on a rotary shaker. The main cultures were inoculated to a start OD_600_ of 0.05 and grown until OD_600_ of 0.8-1 in LB broth with 100 µg/mL Trimethoprim at 37°C at 200 rpm. The cultures were cooled to 4°C and induced with 0.2% L-Rhamnose for expression for ∼24 h at 25°C at 200 rpm. Cells could be harvested and stored at −80°C for several weeks without loss of protein yield. For native purification of BAM, a glycerol stock of *P. aeruginosa* PAO1::2xStrep-BamA was streaked on LB plates and grown overnight at 37°C. The next day, a starter culture was inoculated with several colonies and grown in LB broth at 37°C overnight at 180 rpm. The main cultures were inoculated to a start OD_600_ of 0.05 and grown in LB broth at 37°C at 200 rpm for 15 h. It was important to use high-yielding culturing flasks to ensure good aeration and growth. The best yield was achieved when cultures were harvested while bright green.

### Growth curves and L-type pyocin-treatment

For *P. aeruginosa* PAO1 growth experiments, a glycerol stock was streaked on LB plates and grown overnight at 37°C. The next day, a culture was inoculated with several colonies and grown in LB broth at 37°C at 180 rpm on a rotary shaker for ∼20 h. The culture should reach the stationary phase with an OD_600_ of ∼4.8. The next day, a fresh aliquot of purified Pyocin or darobactin was thawed and centrifuged at maximum speed for 5 min at 4°C. Then, the Pyocin concentration was measured at a wavelength of 280 nm with a NanoDrop Spectrophotometer (Thermo Scientific). The growth curve cultures were inoculated to a start OD_600_ of 0.05 in 125 mL non-baffled Erlenmeyer flasks with 25 mL LB broth, and appropriate amounts of Pyocin, darobactin, and vancomycin were added. Cultures were grown at 37°C at 200 rpm and turned on at the start of the growth experiment. The time points were taken every hour for 8 h, making sure that cultures spent the least possible amount of time outside of the shaker.

### Solid-media overlay growth inhibition assays

For *P. aeruginosa* PAO1 growth experiments, a glycerol stock was streaked on LB plates and grown overnight at 37°C. The next day, cultures were inoculated in a 20-mL scintillation vial with several colonies and grown in 5 mL LB broth at 37°C at 180 rpm on a rotary shaker. The cultures should reach stationary phases and have a range of bright green to yellow pigments depending on the strains. Plates were prepared with 6 mL of LB agar for round plates and 20 mL for square plates and cooled until solid. For the *P. aeruginosa* strain overlay, 0.6% agar was dissolved in boiling water. For round plates, 6 mL of 0.6% agar was transferred to a scintillation vial, and when cool to the touch but not solid, 100 µL of freshly resuspended *P. aeruginosa* overnight culture was added and resuspended before being poured on top of the LB agar plate. For square plates, 12 mL of 0.6% agar was mixed with 200 µL of culture and overlayed in the same way. The agar-cell overlay had to cool completely to avoid condensation. Fresh aliquots of purified L-type pyocins were thawed and centrifuged at maximum speed for 5 min at 4°C. Then, the L-type pyocin concentrations were measured at a wavelength of 280 nm with a NanoDrop Spectrophotometer (Thermo Scientific). Each L-type pyocin was prepared in a 5-fold dilution series with 6 steps in room temperature buffer (50 mM Tris-HCl pH 7.4, 150 mM NaCl), resulting in 1 mg/mL, 0.2 mg/mL, 0.04 mg/mL, 0.008 mg/mL, 0.0016 mg/mL and 0.00032 mg/mL. With a multi-channel pipette, 1 µL samples were applied to the square plates, leaving enough room to prevent the inhibition zones from touching. Round plates were prepared with 1µL of 1 mg/mL Pyocins, except for Pyocin L1Δ17aaCterm, which was used at 85 mg/mL. The plates were incubated at 37°C for no more than 16 h to avoid overgrowth of low-concentration inhibition zones.

### BamA sequence analysis

To assess the presence and conservation of *bamA,* 572 publicly available *P. aeruginosa* genomes were screened using the BLASTN screening tool, Screen Assembly (v1.2.7)^63^, applying cut-offs of 80% identity and 80% reference length with 441 complete *bamA* sequences passing this threshold. PAO1 *bamA* was mapped to an additional 116 whole *P. aeruginosa* genomes using Geneious Prime (v2024.0.7) with default parameters. A total of 557 nucleotide and translated protein sequences were used for variation analysis by Clustal Omega (v1.2.4_1)^64^ and MUSCLE Alignment (v3.8.115)^65,66^ respectively with default parameters. Aligned unique nucleotide sequences were trimmed using Noisy (v1.5.12.1)^66,67^, and phylogenetic trees were constructed using the maximum likelihood approach with IQ-Tree incorporating 1,000 ultrafast bootstrap (v3.0.1)^68^ with no outgroup and visualized using iTol (v7.2.1)^69^. The conservation of translated protein sequences was mapped to the Alphafold3 model (to include the in cryo-EM structure unmodelled PTORA 1 domain) of BamA using Chimera X (1.9)^70^. Unique BamA loop 6 amino acid sequences were extracted from translated protein sequences, aligned, and visualised using Geneious Prime.

### Cloning

Constructs were generated with conventional restriction enzyme cloning and SLiCE cloning^71^. For PCR, Phusion High-Fidelity PCR Master Mix with GC buffer was used (ThermoFischer Scientific, F532L). DNA sequences were confirmed by Nanopore sequencing (Primordium Labs, formerly Plasmidsaurus).

### Allelic exchange

Allelic exchange was done to chromosomally 2xStrep-tag the BamA subunit of *P. aeruginosa* PAO1 for purification and to chromosomally exchange the variable loop 6 region of PAO1-BamA with the loop 6 of P28-BamA. The pEX18Gm constructs were cloned as described before, and the allelic exchange method was adapted and simplified from Hmleo et al. 2015^72^. To make electrocompetent *P. aeruginosa* PAO1, cells were streaked from a glycerol stock onto LB agar and grown at 37°C overnight (∼18h). Then, 5 mL of LB broth in a 20 mL scintillation vial was inoculated with a single colony and grown for 20 h at 42°C in an incubator without shaking. The next day, 4 mL of the culture was harvested in 4 sterile 1.7 mL reaction tubes by centrifugation at maximum speed at room temperature. The supernatant was discarded, and the pellet was resuspended in 1 mL of room temperature 1 mM MgSO_4_ and re-centrifuged. The supernatant was discarded, and the cell pellets were resuspended in another 1 mL of 1 mM MgSO_4_. Finally, the cells were centrifuged as before and combined in a total of 50 µL of 1 mM MgSO_4_.

On the same day, the electrocompetent cells were mixed with 4 µL of allelic exchange plasmid at a concentration of >2µg/µL and incubated at room temperature for 20 min. Electroporation was done at room temperature with a MicroPulser Electroporator (Bio-Rad) at 2.2 kV with a decaying exponential waveform (25 μF capacitor, 600 Ω). Then, 1 mL of LB broth was added to the cells before recovery for 3 h in an orbital shaker at 37°C and 200 rpm. The cells were centrifuged at 3000 x *g* for 5 min, after which they were resuspended in 100 µL of LB and spread on an LB plate containing 20 µg/mL Gentamycin. Within 20 h of growth at 37°C, large positive colonies appeared, but also much smaller negative colonies. A single colony was streaked on a no-salt LB agar plate containing 15% (w/v) sucrose and grown at 30°C overnight. The same colony could be used for an overnight culture to isolate gDNA and perform PCR to confirm the colony as merodiploid by PCR with primers binding in the chromosome and plasmid. From the plate, 8-10 colonies were each resuspended in 50 µL sterile phosphate-buffered saline, after which 3 µL of each colony was patch plated on LB agar, no-salt LB agar containing 15% (w/v) sucrose, and LB agar with 20 µg/mL Gentamycin. In this step, the cells are selected for double cross-over mutants, which means that the allelic exchange plasmid is excised from the chromosome, but the DNA sequence to be exchanged stays in the chromosome, having replaced the original DNA sequence. Positive colonies will grow on plates with LB agar and no-salt LB agar with 15% (w/v) sucrose, but not on LB agar with Gentamycin. The allelic exchange can be verified by isolating a colony’s gDNA and performing PCR with a primer binding in the chromosome and the new exchange DNA sequence.

### BAM purification

For BAM purification from overexpression cultures of *E. coli* BL21(DE3)C41, cultures were grown as described before. The protein yield was ∼30 µg/L of expression culture. Cell pellets were resuspended in lysis buffer (20 mM Tris-HCl pH 8, 150 mM NaCl, 2 mM MgCl_2_, 0.1 mg/mL lysozyme, 0.05 mg/mL DNase, 1x protease inhibitor) and homogenised with a glass homogeniser before lysis with a high-pressure cell disruptor. The lysate was centrifuged at 27,000 x *g* for 20 min at 4°C. Then, the supernatant was centrifuged at 100,000 x *g* for 1 h at 4°C to obtain the bacterial cell membranes. The supernatant was discarded, and the dark red membrane pellet was resuspended in 30 mL solubilization buffer (20 mM Tris-HCl pH 8, 150 mM NaCl, 1% N-dodecyl-β-D-maltoside (DDM), 300 µL biotin blocking buffer (BioLock, 2-0205-050)) with a glass homogeniser. The homogeniser membranes were solubilized at room temperature for 30 min on a slow orbital shaker. After solubilization, the membranes were centrifuged at 30,000 x *g* for 40 min, after which the supernatant was loaded twice on a Strep-Tactin XT 1-mL column (Cytiva) with an Äkta start and Äkta pure purification systems (Cytiva). The DDM detergent was exchanged with lauryl maltose neopentyl glycol (LMNG) detergent by applying 30 column volumes (CV) of detergent-exchange buffer (20 mM Tris-HCl pH 8, 150 mM NaCl, 0.025% Lauryl maltose neopentyl glycol (LMNG). Bound protein was eluted by hand with a syringe containing 10 mL elution buffer (20 mM Tris-HCl pH 8, 150 mM NaCl, 0.025% LMNG, 75 mM biotin) and collected in 1 mL fractions. After 4 mL, the column was incubated in elution buffer for 5 min on ice to optimise protein elution. The fractions were investigated on an SDS-PAGE gel with Coomassie-staining. Clean fractions were pooled and concentrated in a 15-mL concentrator (Amicon Ultra Centrifugal Filter, 30 kDa MWCO) to a final volume of 500 µL for injection on a Superose 6 10/300 Increase column for size exclusion chromatography (20 mM Tris-HCl pH 8, 150 mM NaCl, 0.01% LMNG). The BAM complex eluted at the expected retention volume, and appropriate fractions were concentrated to the highest possible concentrations to ∼200 µL volume to avoid protein loss by excessive use of the concentrator. Purified BAM could be stored in size exclusion buffer at −80°C without disassembly or degradation of the complex.

To remove LMNG from the sample, BAM was thawed and diluted in two times in 15 mL detergent-free buffer (20 mM Tris-HCl pH 8, 150 mM NaCl), then iteratively re-concentrated in a 15-mL concentrator and then 0.5-mL concentrator (Amicon Ultra Centrifugal Filters, 100 kDa MWCO). Because of the high dilution, monomeric LMNG would pass through the filter. The LMNG concentration could be diluted to below the critical micelle concentration because LMNG has a low dissociation rate when the concentration is below the critical micelle concentration. This created favourable detergent-free conditions for the usage of purified BAM.

For the native purification of BAM from *P. aeruginosa* PAO1, the aforementioned protocol was applied with only a few comments. The protein yield was ∼7 µg/L of bacterial cell culture. After lysate was centrifuged at 15,000 x *g* for 20 min at 4°C and depending on how clear the supernatant was, had to be centrifuged a second time at 27,000 x *g* for 10 min. Lastly, LptD was commonly co-purified, which was not the case for BAM purification from *E. coli* overexpression cultures.

### L-type pyocin purification

L-type pyocins were purified from overexpression cultures of *E. coli* BL21(DE3)C41 as described before. The protein yield ranged from 1.2-30 mg/L of expression culture, which was highly dependent on the pyocin and the location of the His6-tag, with N-terminal tags often yielding lower amounts of soluble pyocin. Cell pellets were resuspended in lysis buffer (20 mM Tris-HCl pH 7.4, 150 mM NaCl, 5 mM Imidazole, 2 mM MgCl_2_, 0.1 mg/mL lysozyme, 0.05 mg/mL DNase, 1x protease inhibitor) and homogenised with a glass homogeniser before lysis with a high-pressure cell disruptor. The lysate was centrifuged at 30,000 x *g* for 20 min at 4°C and loaded twice on a HisTrap 5mL column for immobilised metal affinity chromatography on Äkta start and Äkta pure purification systems (Cytiva). After loading, the column was washed with buffer (20 mM Tris-HCl pH 7.4, 500 mM NaCl, 5 mM Imidazole) until the UV absorption at 280 nm read close to 0. For elution, a step gradient (5%, 10%, 25%, 50%, and 100%) of Elution buffer (20 mM Tris-HCl pH 7.4, 150 mM NaCl, 1 M Imidazole) was applied, with pyocins eluting mostly at 250 mM Imidazole. Relevant elution fractions were investigated by SDS-PAGE and Coomassie-staining, and appropriate fractions were pooled. The protein was concentrated in a 15-mL concentrator (Amicon Ultra Centrifugal Filter, 30 kDa MWCO) to a final volume of 10 mL for subsequent injection on a Superdex 26/600 Increase 200pg column to perform size exclusion chromatography (20 mM Tris-HCl pH 7.4, 150 mM NaCl). L-type pyocins eluted at the expected retention volume, and appropriate fractions were concentrated to the highest possible concentrations before protein aggregation.

Pyocins could be stored in size exclusion buffer at −80°C without loss in potency or additional protein aggregation.

### SDS-PAGE analysis

Protein samples were run on a Bolt 4-12% SDS-PAGE gel according to the manufacturer’s instructions. Gel staining was done with AcquaStain Protein Gel Stain (Bulldog) and subsequently washed in distilled H_2_O. Protein samples were generally loaded by constant volume.

### Protein Identification by mass spectrometry

For mass spectrometry, the bands of the purified BAM complex were cut from an SDS-PAGE gel and given to the Monash Proteomics & Metabolomics Platform (Monash University). Protein identities were determined by mass spectrometry fingerprinting after trypsin digestion and compared to a *P. aeruginosa* PAO1 proteome database.

### Cryo-electron microscopy single particle imaging

Cryo-EM imaging was performed on BAM, which was initially purified in DDM and then detergent-exchanged to LMNG with subsequent removal of excess LMNG as described before^73^. For grid preparation, UltrAuFoil gold grids (Quantifoil GmbH) were glow discharged at 15 mA for 180 s with a PELCO easiGlow (Ted Pella Inc.) in air. A 3 µL sample was applied on glow-discharged grids at concentrations of 2.8 mg/mL (12.5 µM PAO1-BAM), 2.6 mg/mL (10.2 µM PAO1-BAM-Pyocin L1), 5 mg/mL (19.5 µM P28-BAM-Pyocin L3), and 5.75 mg/mL (22.6 µM P28-BAM-Pyocin L2). The excess protein solution was blotted off using a blot force of −9 to −12, a blot time of 3 s, and a blot total of 1 with a Vitrobot Mark IV, FEI (Thermo Fisher Scientific), which was maintained at 100% humidity and 4°C. Grids were screened for sample quality, homogeneity, and ice thickness on a Talos Arctica microscope. Imaging was performed at the Ramaciotti Centre for cryo-electron microscopy at Monash University on a G-4 Titan Krios (Thermo Fisher) at 300 kV with a Falcon 4i direct electron detector (apo BAM and PAO1-BAM-Pyocin L1) and at the Ian Holmes Imaging Centre at The University of Melbourne on a Titan Krios at 300 kV with a K3 direct detector and Gatan phase plate (P28-BAM-Pyocin L3) and Falcon 4i direct detector (P28-BAM-Pyocin L2). The fast data collection strategy EPU (Thermo Fischer Scientific), enabled with aberration-free-image-shift (AFIS) alignments, was used for automated data collection. Zero-loss energy-filtered images were acquired at a nominal 105,000x magnification in electron counting mode with a pixel size of 0.75 Å (Monash University) for apo PAO1-BAM and PAO1-BAM-Pyocin L1, at 105,000x magnification in EFTEM mode with a pixel size of 0.833 Å (P28-BAM-Pyocin L3) and at 96,000x magnification in electron counting mode with a pixel size of 0.808 Å (P28-BAM-Pyocin L2) at The University of Melbourne.

For apo PAO1-BAM, 8,372 movies were collected with a total dose of 60 e^−^/Å^2^ accumulated over 5.22 s exposure time at a dose rate of 8.518 e^−^ pixel^−1^ s^−1^ fractionated into 60 frames (**Figure S4**). For image processing, EPU optic groups were determined, and the resultant movies were dose-weighted and motion-corrected using CCSF Motioncor2 1.4.7 to output averages^74^. The non-dose-weighted averages were used for estimating the contrast transfer function (CTF) parameter using CTFFIND 4.1.8^75^ in RELION 5.0^76^. 1,717,815 Particles were picked with TOPAZ particle picker^77^, which had previously been trained on a smaller BAM dataset and extracted with a box size of 320 pixels and binned to 1.5 Å/pix. These particles were imported to cryoSPARC 4.5.3^78^ and 2D-classified, retaining 195,821 particles. Ab initio reconstructions followed by non-uniform refinements resulted in a Coulomb potential map at 3.11 Å with BAM features and LMNG micelle (162,000 particles). These particles were converted to the RELION-compatible file format using the pyem^79^ script and used to train a new TOPAZ picking model, which resulted in the picking of 3,831,721 particles extracted at 320 pixels binned to 1.5 Å/pix. Multiple rounds of 2D-classification, Ab-initio reconstruction, and non-uniform refinements in cryoSPARC resulted in 299,109 particles that were re-extracted at 0.75 Å/pix. 129,206 particles were subjected to Bayesian Polishing in RELION 5.0^76,80^. These particles were subjected to a final round of ab-initio reconstruction and non-uniform refinement, which yielded a Coulomb potential map at a global resolution of 2.75 Å with 121,910 particles (gold standard FSC 0.143 criteria). AlphaFold3 models of the *P. aeruginosa* PAO1 subunits were roughly fit into the map to create 3 (BamA-POTRA 5-3, BamB, BamC/D/E) in ChimeraX 1.8^81^. These maps were expanded with soft edges using empirically determined thresholds with a custom bash script (Dr. Matthew Belousoff) and used for local refinement to increase map quality.

For PAO1-BAM-Pyocin L1, 11,513 movies were collected with a 60 e^−^/Å^2^ accumulated over 5.22 s exposure time at a dose rate of 8.518 e^−^ pixel^−1^ s^−1^ fractionated into 60 frames (**Figure S6**). For image processing, EPU optic groups were determined, and the resultant movies were dose-weighted and motion-corrected using CCSF Motioncor2 1.4.7 to output averages. The non-dose-weighted averages were used for estimating the contrast transfer function (CTF) parameter using CTFFIND 4.1.8 in RELION 5.0. 1,165,588 particles were picked with TOPAZ particle picker, which had previously been trained on a smaller PAO1-BAM dataset and extracted with a box size of 320 pixels and binned to 1.5 Å/pix. These particles were imported to cryoSPARC 4.5.3 and 2D-classified, retaining 132,991 particles. Ab-into reconstruction with 2 classes (no similarity) resulted in one class (85,140 particles) that showed BAM subunits, LMNG micelle, and additional density corresponding to Pyocin L1. A non-uniform refinement followed by local refinement with the cryoSPARC-generated mask resulted in a Coulomb potential map at a global resolution of 3.32 Å. These particles were converted to the RELION-compatible file format using the pyem script and used to train a new TOPAZ picking model, which resulted in the picking of 3,141,875 particles extracted at 320 pixels binned to 1.5 Å/pix. Multiple rounds of 2D-classification, Ab-initio reconstruction, and non-uniform refinements in cryoSPARC resulted in 162,354 particles that were re-extracted at 0.75 Å/pix. These particles were subjected to Bayesian polishing in RELION 5.0. In cryoSPARC, an ab initio reconstruction with 2 classes (no similarity) resulted in 1 incomplete map and one complete map with 110,600 particles, which was used for non-uniform refinement, which yielded a map at a global resolution of 2.84 Å (gold standard FSC 0.143 criteria). AlphaFold3 models of the *P. aeruginosa* PAO1 subunits and the crystal structure of Pyocin L1 (pdb: 4LE7) were roughly fit into the map to create 3 (BamA-Pyocin L1, Bam-BamA-POTRA 3 B, BamC/D/E) in ChimeraX 1.8. These maps were expanded with soft edges using empirically determined thresholds with a custom bash script (Dr. Matthew Belousoff) and used for local refinement, which significantly increased map quality, especially for the BamA-Pyocin L1 regions.

For P28-BAM-Pyocin L2, 11,319 movies were collected at 53 e^-^/Å^2^ accumulated over 7.6 s exposure time at a dose rate of 4.55 e^−^ pixel^−1^ s^−1^ fractionated into 60 frames (**Figure S7**). For image processing, EPU optic groups were determined, and the resultant movies were dose-weighted and motion-corrected using CCSF Motioncor2 1.4.7 to output averages. The non-dose-weighted averages were used for estimating the contrast transfer function (CTF) parameter using CTFFIND 4.1.8 in RELION 5.0. 5,533,458 particles were picked with TOPAZ particle picker with the PAO1-BAM-Pyocin L1 trained model and extracted with a box size of 320 pixels and binned to 1.6 Å/pix. These particles were imported to cryoSPARC 4.5.3 and 2D-classified, retaining 1,350,291 particles. Ab-into reconstruction was done with 5 classes (no similarity), followed by 2D classification of all classes and selecting good 2D averages from each class. A second ab initio reconstruction with 2 classes (no similarity) resulted in one good class with 352,957 particles, which had the BAM subunits, LMNG micelle, and additional density corresponding to Pyocin L2. Two separate approaches for 2D classification to selectively obtain side-views and top/bottom views resulted in a combined 300,949 particles that were used for non-uniform refinement, generating a coulomb potential map of 3.56 Å global resolution. These particles were converted to the RELION-compatible file format using the pyem script and were re-extracted at 0.808 Å/pix. In cryoSPARC, ab initio reconstruction with 2 classes (no similarity) resulted in one incomplete map and one complete map. The better map was used for non-uniform refinement with 126,889 particles reaching 3.47 Å. In ChimeraX 1.8, a rough fit of P28-BamA-POTRA 5 with Pyocin L2 was used to generate a mask in cryoSPARC. This mask was used for local refinement, which yielded a coulomb potential map with a global resolution of 3.18 Å (gold standard FSC 0.143 criteria). Bayesian polishing of the particles did not improve the map quality and resolution.

### Molecular dynamics simulations

Classical atomistic molecular dynamics (MD) simulations were carried out on the two cryo-EM structures: PAO1-BAM-Pyocin L1 pre-inhibited complex (2.84 Å, pdb: 9PXI), and P28-BAM-Pyocin L2 fully inhibited complex (3.18 Å, pdb: 9PXJ). To keep the system size tractable and achieve sufficient simulation sampling, the model systems comprising pyocin (L1 or L2) and BamA subunits were constructed, in which the domains POTRA 1 to POTRA 4 (residue IDs 1 to 381 of BamA) were excluded. All titratable residues were set to their standard protonation states. The protein system was embedded into a lipid bilayer representing the composition of the outer membrane of gram-negative bacteria, with the inner leaflet consisting of 75% phosphoethanolamine (PPPE), 20 % phosphoglycerol (PVPG), and 5% cardiolipin (PVCL2) lipids, and the outer leaflet carrying lipopolysacharides (LPSA) without O antigen units (see below). To compensate for the negative charge of the lipid bilayer (LPSA: charge −10*e*; PVCL2: charge −2*e*; PVPG: charge −1*e*), the system was neutralised by Ca^2+^ ions, and solvated with 150 mM of Na^+^ and Cl^-^ ions. The placement of the protein-membrane model system in a periodic water box of 10 nm x 17 nm x 21 nm resulted in a total count of ∼390,000 atoms (Table S4).

For the construction and relaxation of lipid bilayers, a pure membrane system was constructed and relaxed by MD simulations. The G-OM membrane patch from gram-negative bacteria^82,83^ (size 8.1 nm x 8.1 nm) was replicated 8 times, yielding a larger patch of 24 nm x 24 nm. After immersing the larger membrane patch in a water box, and neutralising it with Ca2+ ions, and 150 mM of Na+ and Cl-ions, the system was minimised with a conjugate-gradient algorithm implemented in NAMD (v. 2.14)^84^ for 1000 steps with constraints on the phosphorous atoms (force constant 4.1 x 10^4^ kJ/(mol*nm2)). The same atoms were constrained in an additional energy minimisation in GROMACS (v. 2022.3)^85^ with a force constant of 2.0 x 10^4^ kJ/(mol*nm2) and force tolerance of 250 kJ/(mol*nm), followed by a 50-ns equilibration in an NPT ensemble. Finally, a production run of 100 ns produced a relaxed membrane patch, which was used for the construction of our protein-membrane systems (see above).

In order to study the conformational dynamics of BamA-pyocin complex, we constructed four different simulation setups based on the pre-inhibited and inhibited structures (see above), and in the presence and absence of pyocin (Table S4). To eliminate possible steric clashes, each protein-membrane setup was energy minimised using the conjugate-gradient algorithm in NAMD (v. 2.14)^84^ for 1000 steps with positional harmonic restraints (force constant 4.1 x 10^4^ kJ mol^-1^ nm^-2^ on lipid phosphorous atoms, and protein heavy atoms, excluding the modelled regions (Table S4, see also main text). This was followed by an additional energy minimization in GROMACS (v. 2022.3; force tolerance of 250 kJ mol^-1^ nm^-1^) with the same atoms constrained (force constant 2.0 x 10^4^ kJ mol^-1^ nm^-2^). Equilibration of the constructed model system was achieved with a sequence of MD simulation steps. First, a short 10-ns equilibration was performed (with the harmonic restraints on protein heavy atoms and lipid/LPS phosphorous atoms), allowing the system’s pressure and temperature to equilibrate to 1.0132 bar and 310 K (37°C), respectively, while maintaining the structural integrity of the protein and the membrane. After that, we released the constraints from lipids (keeping restraints on the protein heavy atoms) and performed a 100-ns equilibration, leading to the proper relaxation of the mixed lipid bilayer. The last step involved equilibration with the protein backbone restraints (force constant 2.0 x 10^4^ kJ mol^-1^ nm^-2^) for 15 ns, where during the last 5 ns, the restraints from the modelled regions were released. The production simulations were obtained in a set of 3 replicas, ∼ 1 μs each (**Table S4**), where replicas 2 and 3 were generated via extending the second equilibration step by 1 ns and 2 ns, respectively. CHARMM36m force field^86^ was used for protein, while lipids and ions were described by CHARMM36^87–89^. TIP3P model was applied to describe water solvent^90^. NBFIX corrections were applied to Na+, Cl-, and Ca2+ ions, as implemented in CHARMM36 force field^87,89,91^. Integration timestep of 2 fs was achieved via applying rigid constraints to all covalent bonds to hydrogen atoms with the LINCS algorithm^92^. Short-range non-bonded interactions were described by the Verlet^93^ cutoff scheme with the switching and cutoff distances of 1.0 nm and 1.2 nm, respectively, whereas long-range electrostatics was treated by the particle mesh Ewald method^94^. During the equilibration runs, temperature was maintained at 310 K using the V-rescale thermostat^95^, while a stable pressure of 1.0132 bar was achieved by the Berendsen barostat^96^. At the production phase, the thermostat was switched to Nose-Hoover^97,98^, and barostat to Parrinello-Rahman^99^.

All simulation trajectories were analysed with MDAnalysis (v. 2.7.0)^100^ and VMD (v. 1.9.4) visualisation tool. Hydrogen bonds were calculated using the Hbonds plugin (v. 1.2) implemented in VMD^101^.

### Cryo-electron microscopy structure and model visualization

The structures of this study were built using the crystal structure of Pyocin L1 (pdB:4LE7) and AlphaFold3 predictions of PAO1-BamA-E and P28-BamA. In ChimeraX 1.8, the models were docked into the Coulomb potential map and roughly adjusted using ISOLDE 1.8. Phenix 1.2, in combination with WinCoot 0.9.8.1 was used for building the structures and refinement. The CheckMyMetal web tool was used to determine the most likely metal for a density in the apo BAM subunit BamB. Model quality was assessed and confirmed with the PDB validation web tool. Images were created using ChimeraX 1.8. After structural models were built and refined with cryoSPARC-derived cryo-EM density maps, the AI map-enhancement tool EMReady was used exclusively for map visualisations of main-text figures^102^.

### RNA preparation form Pyocin L1 and darobactin-treated cells

RNA for sequencing was derived from four independent experiments, with three technical replicates performed for each condition in each experiment. An overnight culture of *P. aeruginosa* PAO1 was subcultured with a 1/100 dilution and grown for 2 hours and 15 min at 37 °C with shaking. The culture was then aliquoted into 2 mL samples, and darobactin, Pyocin L1 (N-terminal histagged cleaved using TEV protease) or Pyocin L2 (N-terminal histagged cleaved using TEV protease) was added at 1µg/mL (MIC) (or no antibiotic for untreated reference). 500 µL samples were taken at 15 min and 120 min timepoints and added to RNA Protect solution (Qiagen). The samples were left for 5 min at room temperature, spun down, supernatant removed, and then frozen at −80°C until the RNA was extracted. The untreated control was only sampled at the 15-minute timepoint. The Qiagen RNAprotect Bacteria Reagent Handbook protocol 4, protocol 7 and on-column DNase digestion using the Qiagen RNase-free DNase set were followed. 15 µL of each of the three technical replicates was combined to give 1 biological replicate (12 samples in total for sequencing).

### RNA sequencing and Bioinformatic analysis

RNA-seq libraries were sequenced at the Wellcome Trust Sanger Institute on an Illumina HiSeq 2500 using 2 × 100 bp paired-end runs. Reads containing ambiguous bases (“N”) or lacking all four nucleotides were discarded, and the remaining reads were aligned to the *P. aeruginosa* PAO1 genome (GenBank GCA_000006765.1) using BWA-SW v0.7.10^103^. For each read, the best alignment was retained based on mapping quality, and reads with multiple equally good hits were removed. Because the library preparation method captures essentially all transcripts, including small RNAs, read lengths were highly variable. We therefore quantified expression as the total number of nucleotides mapping to each gene (using the PseudoCAP GFF3 annotation^104^), normalized by gene length and total genic coverage, analogous to Transcripts per million^105^. Differential expression was assessed using edgeR^106^, with significance defined as Log_2_FC > 0.5 and −log_10_FDR > 1.3.

### Proteomic analysis of Pyocin L1 and darobactin-treated cells

The whole cell lysates were processed for proteomic analysis using an S-Trap (PROFIT) –based workflow. Protein samples were solubilized in 50 µL of SDS buffer containing 5% SDS and 50 mM triethylammonium bicarbonate (TEAB, pH 7.55), reduced with tris(2-carboxyethyl)phosphine (TCEP) to a final concentration of 10 mM, and incubated at 37 °C for 45 min. Alkylation was performed in the dark with iodoacetamide (IAA) at a final concentration of 55 mM under the same temperature and duration. Samples were acidified to approximately 2.5% phosphoric acid, and S-Trap binding buffer (90% methanol, 100 mM TEAB, pH 7.1) was added before loading onto S-Trap columns. Bound proteins were washed three times with binding buffer, then digested on-column overnight at 37 °C with trypsin (1:10 enzyme-to-protein ratio, prepared in 50 mM TEAB). Peptides were sequentially eluted with 40 µL of 50 mM TEAB, 0.2% aqueous formic acid, and 35 µL of 50% acetonitrile with 0.2% formic acid. The combined eluates were centrifuged, lyophilized, and reconstituted in 2% acetonitrile with 0.1% trifluoroacetic acid for subsequent LC-MS/MS analysis.LC-MS/MS Analysis: 20250204_IS-10197 Thermo Scientific Orbitrap Eclipse and 250404_IS-10601 Thermo Scientific Orbitrap Ascend.

For LC–MS/MS analysis, lyophilized peptide samples were analysed with a Thermo Scientific Orbitrap Eclipse Tribrid mass spectrometer coupled to an Ultimate 3000 RSLC nano-flow reversed-phase HPLC system (Dionex). The nano-LC was equipped with an Acclaim PepMap 100 C18 nano-trap column (100 Å pore size, 75 µm × 2 cm, Thermo Scientific) and an Acclaim PepMap RSLC C18 analytical column (100 Å pore size, 75 µm × 50 cm, Thermo Scientific). Mobile phases consisted of solvent A (0.1% formic acid in LC–MS–grade water) and solvent B (100% ACN with 0.1% formic acid). For each run, peptides were loaded onto the trap column at 6 µL/min in an isocratic solution of 2% ACN and 0.05% TFA for 6 min before being switched in-line with the analytical column. Peptides were separated over a 95 min gradient: 3% B at 0–6 min, increased to 4% B at 7 min, then to 25% B at 82 min, followed by a ramp to 40% B at 86 min and to 80% B at 87 min; the column was held at 80% B until 90 min, then re-equilibrated at 3% B until 95 min. The mass spectrometer was operated in positive ion mode with a spray voltage of 1,900 V, ion transfer tube temperature of 275 °C, and HCD collision gas pressure of 1 mTorr. Data were acquired in data-independent acquisition (DIA) mode with an initial full MS1 scan in the Orbitrap (m/z 350–1400; resolution 120,000 at m/z 200; AGC target 1 × 10⁶; maximum injection time 50 ms; profile mode) followed by DIA MS2 scans in the Orbitrap at a resolution of 30,000 (m/z 200–2000; AGC target 1 × 10⁶; maximum injection time 55 ms; centroid mode) using higher-energy collisional dissociation (HCD) at 30% normalized collision energy. Fifty sequential quadrupole isolation windows (13.7 m/z width, 1 m/z overlap) were used to cover the precursor m/z range of 361–1033, providing ∼9 points across chromatographic peaks with a total cycle time of 3 s.

#### Data Analysis

Global proteomics raw data were processed in Spectronaut (version 19.1.240806.62635, Huggins) using default BGS factory settings. Searches were performed against the *P. aeruginosa* PAO1 (October 2024 release, 26,816 protein entries). DIA (Data-Independent Acquisition) analysis employed maximum intensity for m/z extraction, with precursor and protein identifications filtered at a q-value threshold of 0.01 and a protein PEP cutoff of 0.75. Decoys were generated by mutation using a dynamic decoy limit strategy. Protein quantification was MS2-based with automatic and cross-run normalization, as well as interference correction. For peptide identification, the directDIA+ workflow was applied using Trypsin/P as the proteolytic enzyme, allowing up to two missed cleavages. Carbamidomethylation of cysteine was specified as a fixed modification, while oxidation of methionine and acetylation of protein N-termini were set as variable modifications. Peptide and protein identifications were controlled at a 1% false discovery rate (FDR). Differential abundance analysis was performed using Welch’s t-test (unpaired, unequal variances, no variance filtering or group-wise correction), with p-values adjusted using q-value confidence filtering (FDR threshold of 0.05). A log₂ fold-change cutoff of 0.58 was applied to define differentially abundant proteins. MSMS data have been deposited to the ProteomeXchange Consortium, via the PRIDE partner repository, with the dataset identifier PXD067326. (For review only purposes Username: Password: JGAaWcO9BayM)

### *P. aeruginosa* PAO1 transposon library preparation for TraDIS

A high-density *P. aeruginosa* PAO1 transposon mutant library was generated using biparental conjugation with *E. coli* S17-1 harbouring plasmid pUT-mini-Tn5-pro. Donor and recipient strains were streaked on LB agar and LB agar supplemented with 15 μg/mL gentamicin (LBA/Gent15), respectively, and incubated overnight at 37 °C. Overnight cultures were established in LB or LB/Gent15 and diluted to OD_600_= 0.1 for subculturing. After 7 h incubation at 37 °C with shaking, 200 μL of donor and 100 μL of recipient were each spread onto 35 pre-warmed 1.5% LBA or LBA/Gent15 plates using sterile glass beads and incubated overnight in humidified conditions (donor at 37 °C, recipient at 44.5 °C). The next day, VBMA (1.5% agar in Vogel & Bonne (VB) minimal medium) and VBMA/Gent30 (VBMA with 30 μg/mL gentamicin) plates were prepared by autoclaving agar in water (12 g in 720 mL), cooling, and supplementing with VBMA components (80 mL per 800 mL) and filter-sterilised gentamicin. Mating mixtures were established by scraping donor and recipient lawns from overnight plates using sterile loops and co-spotting onto fresh LB agar plates. Conjugations were set up in batches (28 total, labelled A–D), mixed for 10 min, and incubated for 2 h at 37 °C in humidified bags. Cells from conjugation spots were scraped, resuspended in LB, and plated onto VBMA and VBMA/Gent30 plates at multiple dilutions (10^-4^ to 10^-12^) for recovery of transconjugants. After 16 h at 37 °C, colonies were counted and pooled by conjugation set (A–D), resuspended in LB + 15% glycerol, and aliquoted into 1 mL cryovials for storage at –80 °C. CFU and OD₆₀₀ measurements were recorded, and mutant numbers were estimated to exceed 2 x 10^6^ total transformants.

### *P. aeruginosa* PAO1 Pyocin L1 treatment TraDIS experiment

Pyocin L1-supplemented TYE plates (10 g tryptone, 5 g yeast extract, 5 g NaCl, 0.8% agar per L H_2_O) were prepared by adding 14 nM (0.4 μg/mL) of Pyocin L1 to 40 mL TYE. Frozen aliquots of the mutant library and pyocin were thawed on ice for 15 min, diluted 1:1 in TYE, and 100 μL was spread onto prepared plates, which were dried under sterile conditions. For colony-forming unit (CFU) enumeration of the transposon library, serial 10-fold dilutions were plated in 20 μL spots. Bacterial lawns were harvested by plate scraping and DNA extraction was performed using the DNeasy Blood & Tissue Kit (Qiagen); cells (1 mL) were pelleted (10 min, 7,500 × g), resuspended in ATL buffer (180 μL) and Proteinase K (20 μL), vortexed, and incubated at 56 °C for 1.5 h with shaking. DNA was purified per the manufacturer’s protocol and eluted in nuclease-free water. Concentration and purity were assessed by Nanodrop (A260/280 and A260/230 ratios) and Qubit dsDNA HS Assay. DNA from technical and biological replicates was pooled (30 μL each; 120 μL total per condition) and normalised to ∼8.1 ng/μL before TraDIS library preparation and sequencing.

TraDIS sequencing libraries were prepared as described Barquist et al, 2016 ^40^, with an additional EcoRI digestion step performed following adapter ligation and prior to amplification to deplete vector DNA. Amplification of transposon-genomic DNA junctions was performed using the outward facing primer AATGATACGGCGACCACCGAGATCTACACAACTCTCTACTGTTTCTCCATACCC G and libraries were sequenced using primer GGCTAGGCCGCGGCCGCACTTGTGTA with a TraDIS-specific single-end MiSeq protocol ^40^. Reads were filtered for presence of the transposon tag sequence, mapped to the PAO1 genome, and assigned to genomic features using the bacteria-tradis pipeline, with multiple mapping reads excluded and reads mapping to the 3’ 10% of a gene excluded from each feature-wise count. Mutant abundance following selection was calculated using the tradis_comparison function of the Biotradis pipeline.

### Cryo-electron tomography

*P. aeruginosa* cells were treated with Pyocin L1 and darobactin as described before. At the desired treatment time, just before grid preparation, cultures were centrifuged and resuspended in ice-cold PBS buffer, and the OD_600_ was adjusted to ∼1. Untreated, pyocin- and darobactin-treated *P. aeruginosa* cultures were mixed in a 4:1 ratio with 10 nm colloidal gold beads that had been precoated with 1% BSA (Sigma-Aldrich, Australia). 4 µL of culture was applied to glow-discharged copper R2/2 Quantifoil holey carbon grids (Quantifoil Micro Tools GmbH, Jena, Germany). Depending on the sample, grids were blotted for 4-5 seconds (under 100% humidity conditions) with a blot force of 6 and plunged into liquid ethane using a Vitrobot Mark IV, FEI (Thermo Fisher Scientific). Tilt series were collected automatically with the FEI Tomography 5 software from −60° to +60° at 2° intervals with a defocus of −6 µm and a total electron fluence of 120 e^-^/Å^2^ at a pixel size of 3.4 Å using an FEI Titan Krios G4, 300 keV FEG transmission electron microscope (Thermo Fisher Scientific), equipped with a BioQuantum K3 Imaging Filter (slit width 20 eV), and a K3 direct electron detector (Gatan) in movie mode (4 dose fractions).

Tilt series movies were motion corrected and split into odd/even using motioncor3. Odd-even tilt series were pre-processed to bin4 (13.56 Å/pix), aligned using fiducial tracking, and reconstructed by weighted-back projection in IMOD (Version 5.1). Cryocare software was used to denoise tomograms using odd and even tomograms and a u-net 2.5 trained model (training was done using a subset of tomograms from each data collection). 2D slice Images of 3D tomograms were generated using 3dmod (IMOD, version 5.1) using n=4 slices. Denoised tomograms were binned further (bin8, 27.12 Å/pix) for processing by 3D segmentations. All these steps were done using the scipion3 framework. Visualisation and segmentation of tomograms were done using the Dragonfly software (https://www.theobjects.com/dragonfly/index.html). Tomograms were pre-processed using built-in histogram equalization filters. Neural network 5-class U-Net (with 2.5D input of 5 slices) was trained on tomogram slices to recognize *P. aeruginosa* inner- and outer membranes, ribosomes, flagella, T6SS, and cytoplasmic DNA using 5 rounds of optimisation. The software’s built-in tools for exporting 2D images and creating 3D movies were utilised. Movies were annotated and edited using DaVinci Resolve software.

### Bioinformatics analysis of protein sequences

Protein sequences (BAM and L-type pyocins) were aligned in MEGA11 v11.0.13 using the built-in ClustalW alignment tool (Multiple Sequence alignment: Gap Opening Penalty 10.00, Gap Extension Penalty 0.20, Divergent Cutoff 30% delayed). The software TreeViewer v2.2.0 was used for the generation of phylogenetic trees with neighbour-joining of an aligned sequence file (BLOSUM62 distance model and 1000 bootstrap replicates). The STRiNG webserver was used for analysis of TraDIS, RNA sequencing and proteomics dataset ^107^.

### Protein structure prediction

For the prediction of protein structures, AlphaFold2 multimer was used on the MASSIVE M3 High-Performance Computing cluster^108,109^. AlphaFold3 was used for the prediction of protein structures using the AlphaFold Server (https://alphafoldserver.com, state 22nd April 2025)^110^.

## Data Availability

Cryo-EM maps and atomic models generated in this paper have been deposited in the Protein Data Bank and the Electron Microscopy Data Bank (Accession codes: BAM_PAO1_ = 9PXG, EMD-71969; BAM_PAO1_-Pyocin L1 = 9PXI, EMD-71970; BAM_P28_-Pyocin L2 = 9PXJ EMD-71971). Data availability: Raw TraDIS reads are available from European Nucleotide Archive, study ERP106484. Individual sample accessions are provided in table S1. Individual sample accessions are provided in **Table S2**. Raw data from Proteomics experiments is available at: PXD067326. Raw data from Transcriptomics experiments will be available on final publication of the manuscript. Raw data from the BAM_PAO1_-Pyocin L1 molecular dynamics simulations is available on request. Raw data and numerical data for all experiments are provided with the manuscript.

## Acknowledgements

The authors acknowledge the use of electron microscopy and cryo-sample preparation facilities at the Ian Holmes Imaging Centre of the Bio21 Molecular Science & Biotechnology Institute, the University of Melbourne; in particular, Dr. Sepideh Valimehr for training and assistance, and at the Ramaciotti Centre for Cryo-Electron Microscopy of the Biomedicine Discovery Institute, Monash University. This research was supported by ARC LIEF grants (LE200100045, LE120100090) for the Titan Krios Gatan K3 Camera and the Titan Krios, and an ARC discovery project grant awarded to R.G. and G.K. (DP230102150). Non-CF bronchiectasis isolates were derived from samples in the David Serisier Research Biobank at Mater Misericordiae Ltd. O.Z. and V.S. acknowledge large-scale HPC resources from the Center for Scientific Computing (CSC), Finland – the IT Center for Science. F.M., M.D.J., D.G., and R.G are members of the Australian Research Council Industrial Transformation Training Centre for Cryo-Electron Microscopy of Membrane Proteins for Drug Discovery (IC200100052). C.A.M. was supported by NHMRC Ideas grant (2010400) and ARC Discovery Project (DP220100713). R.G. was funded by an NHMRC EL1 investigator grant (APP1197376). G.K. was supported by the Snow Medical Research Foundation (SMRF2021-276). L.C.M was funded by a Sir Henry Wellcome Fellowship from the Wellcome Trust 9106064/Z/14/29. D.G. is funded by an NHMRC EL1 grant (APP1196924) and a Cumming Global Centre for Pandemic Therapeutics Foundation grant, Peter Doherty Institute for Infection and Immunity (CGCPT-CGCPT00060); F.S. was funded by a Sir Henry Wellcome Fellowship (106063/A/14/Z) and an ARC DECRA fellowship (DE200101524). TraDIS sequencing was supported by Wellcome (206194), O.Z. is supported by the Research Council of Finland. V.S. acknowledges research funding from the Research Council of Finland, the Sigrid Jusélius Foundation and the Jane and Aatos Erkko Foundation. This research was supported by The University of Melbourne’s Research Computing Services and the Petascale Campus Initiative. The authors acknowledge the use of the facilities at the Melbourne Protein Characterisation Facility, of the Bio21 Molecular Science & Biotechnology Institute, the University of Melbourne and thank Belinda Michell and Troy Attard for training and assistance. We thank Kim Lewis (Antimicrobial Discovery Center, Department of Biology, Northeastern University, Boston, MA, USA) for kindly providing us with the darobactin used in this study.

## Author Contributions

Conceptualization: R.G., F.M., L.M.,

Data curation: F.M., R.G., M.J., I.S., L.C.M., O.Z., L.Z. E.P.P, D.S.S, S.V., F.S., J.P.R.C.,

Formal analysis: F.M., R.G., M.J., I.S., L.C.M., O.Z., L.Z., C.A.M., E.P.P, D.S.S, S.V., F.S., J.P.R.C.

Funding acquisition: R.G., G.K., J.P.R.C., L.C.M., O.Z., V.S., D.G.

Investigation: F.M., R.G., M.J., I.S., L.C.M., O.Z., C.W., A.K., L.Z., E.P.P., D.S.S, H.V., F.S., J.P.R.C., R.G.

Methodology: F.M, R.G., M.J., I.S., L.C.M., O.Z., A.K., L.Z., C.A.M. E.P.P., D.S.S., S.V., H.V., V.S., M.T.D., F.S., D.G., J.P.R.C., G.K.

Project administration: R.G., G.K., J.P.R.C., L.C.M., V.S., D.G.

Resources: L.C.M., E.P.P., D.S.S, H.V., V.S., F.S., D.G., J.P.R.C., G.K., R.G. Supervision: R.G., D.G., C.A.M., M.T.D., V.S., S.V.

Validation: F.M., M.J., I.S., L.C.M., O.Z., E.P.P., D.S.S, S.V., H.V., V.S., M.T.D., F.S., D.G., J.P.R.C., G.K., R.G.

Visualization: F.M., R.G.

Writing – original draft: F.M., R.G.

Writing – reviewing & editing: all authors

## Supplemental Figures

**Figure S1:**
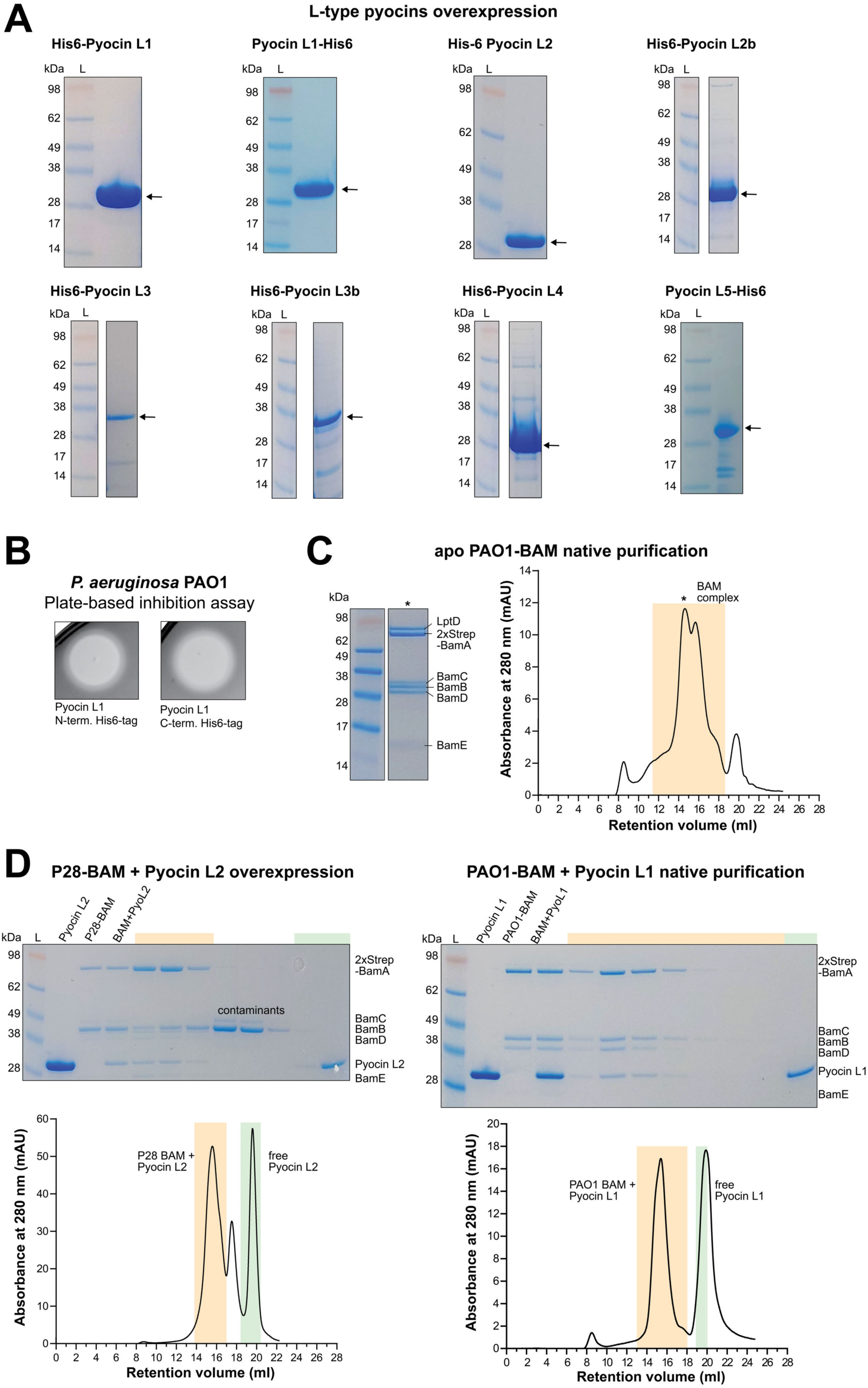
Representative purified protein samples used in this study. (A) Coomassie-stained SDS-PAGE gels of Pyocin L1, L2, L2b, L3, L3b, L4, and L5. The black arrow indicates the protein band corresponding to the expected molecular weight of the pyocin. All L-type pyocins were purified to high quality and purity. (B) Plate-based inhibition assay of N-terminal His6-tagged Pyocin L1 and C-terminally His6-tagged Pyocin L1 showing comparable inhibition activity. Both Pyocin L1 versions exhibit a similar activity, but other L-type pyocins are less active or inactive with C-terminal His6-tags. (C,D) Coomassie-stained SDS-PAGE gels and corresponding size exclusion chromatograms for BAM purification and BAM + Pyocin L-type pyocins. Fractions of the chromatogram and corresponding SDS-PAGE samples are colour coded in orange (BAM) and L-type pyocin (green).

**Figure S2:**
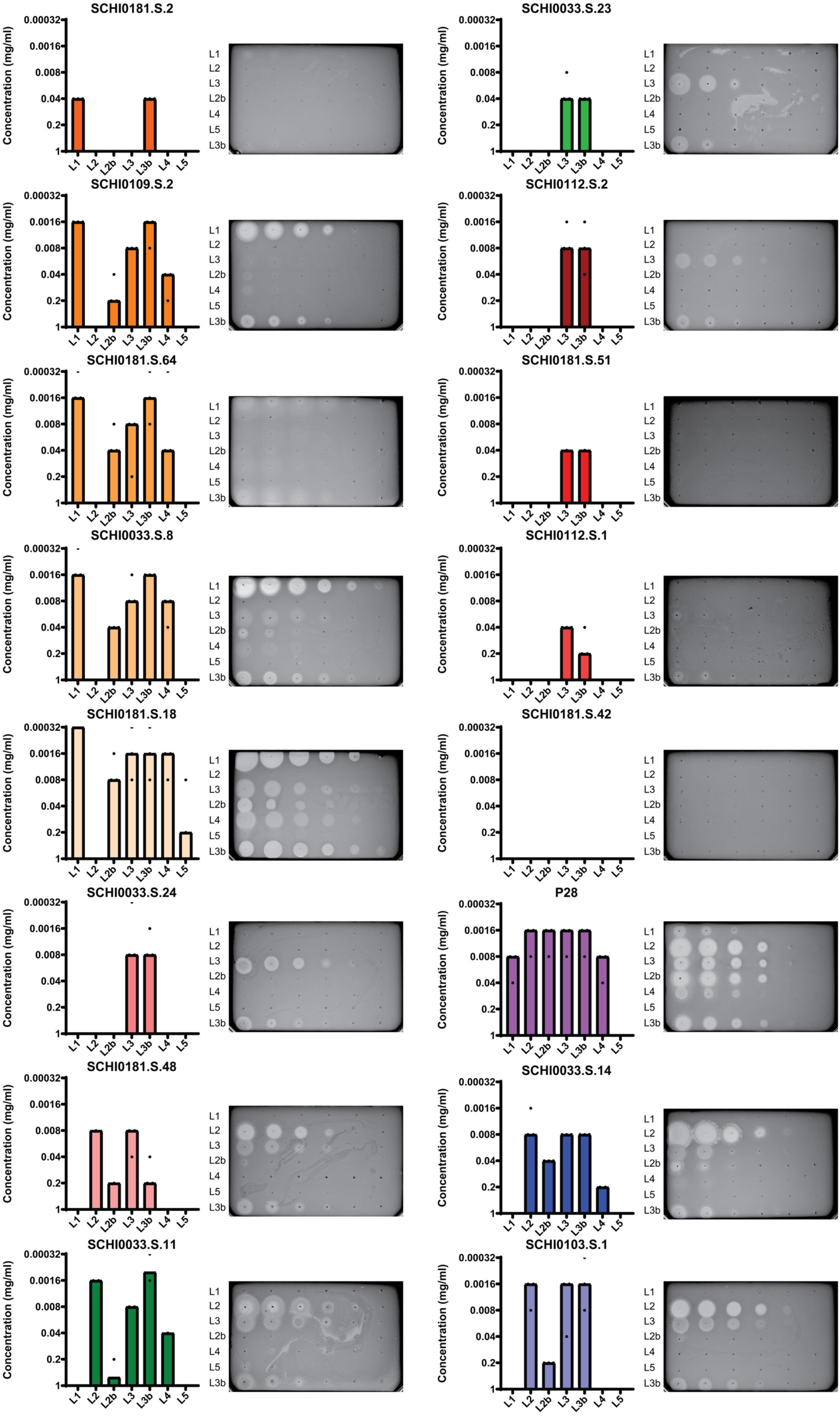
Activity of L-type pyocins against *P. aeruginosa* clinical isolates. Plate-based inhibition screening of 7 L-type pyocins against 17 *P. aeruginosa* strains. Bar graphs (left) of three replicates and their lowest effective concentration. Representative inhibition plates (right) showing inhibition zones. The inhibition screen of *P. aeruginosa* PAO1 is shown in Fig. 1E. Plate colours have been changed to Greyscale.

**Figure S3.**
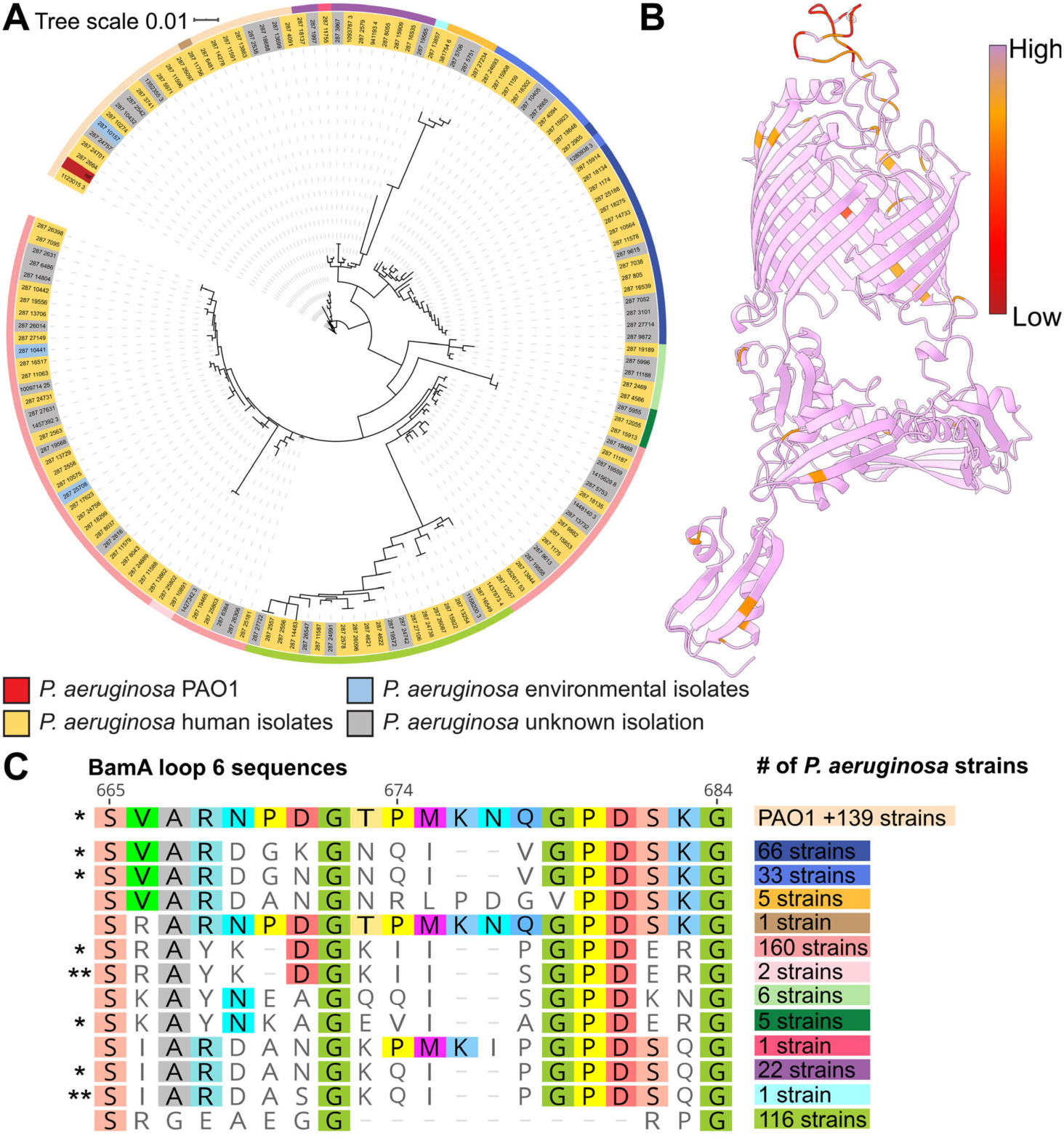
Conservation of *bamA* in *P. aeruginosa* species. (A) Phylogenetic analysis depicting sequence diversity across 151 unique *bamA* nucleotide sequences from 557 publicly available *P. aeruginosa* genomes that encoded a *bamA* gene. Human and environmental isolates are as indicated, and the reference PAO1 sequence is indicated in red. Sequences of BamA loop 6 are denoted in the outer ring with colours corresponding to C. (B) The amino acid conservation of the BamA multiple sequence alignment mapped onto the Alphafold3 model of PAO1-BamA. (C) Sequence alignment of BamA loop 6 regions in 557 *P. aeruginosa* strains. A single asterisk (*) indicates this sequence can encoded by one of the 17 experimentally tested loop 6 variations, and two asterisks (**) indicates a single amino acid change compared to one of the 17 experimentally tested loop 6 variations.

**Figure S4:**
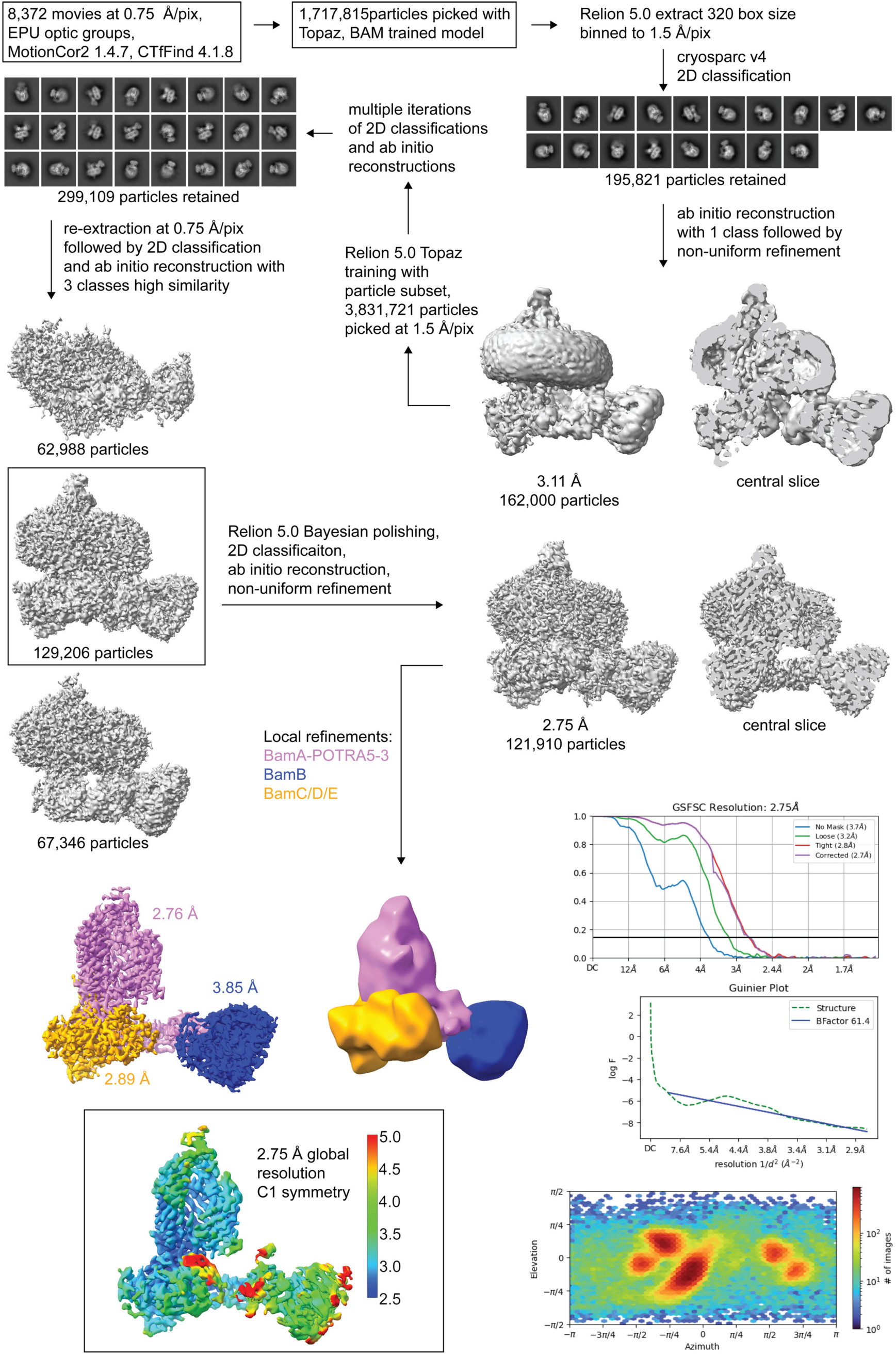
Cryo-EM SPA processing workflow of apo PAO1-BAM. Workflow of cryo-EM data processing of apo BAM from *P. aeruginosa* PAO1. Detailed information can be found in the method section.

**Figure S5:**
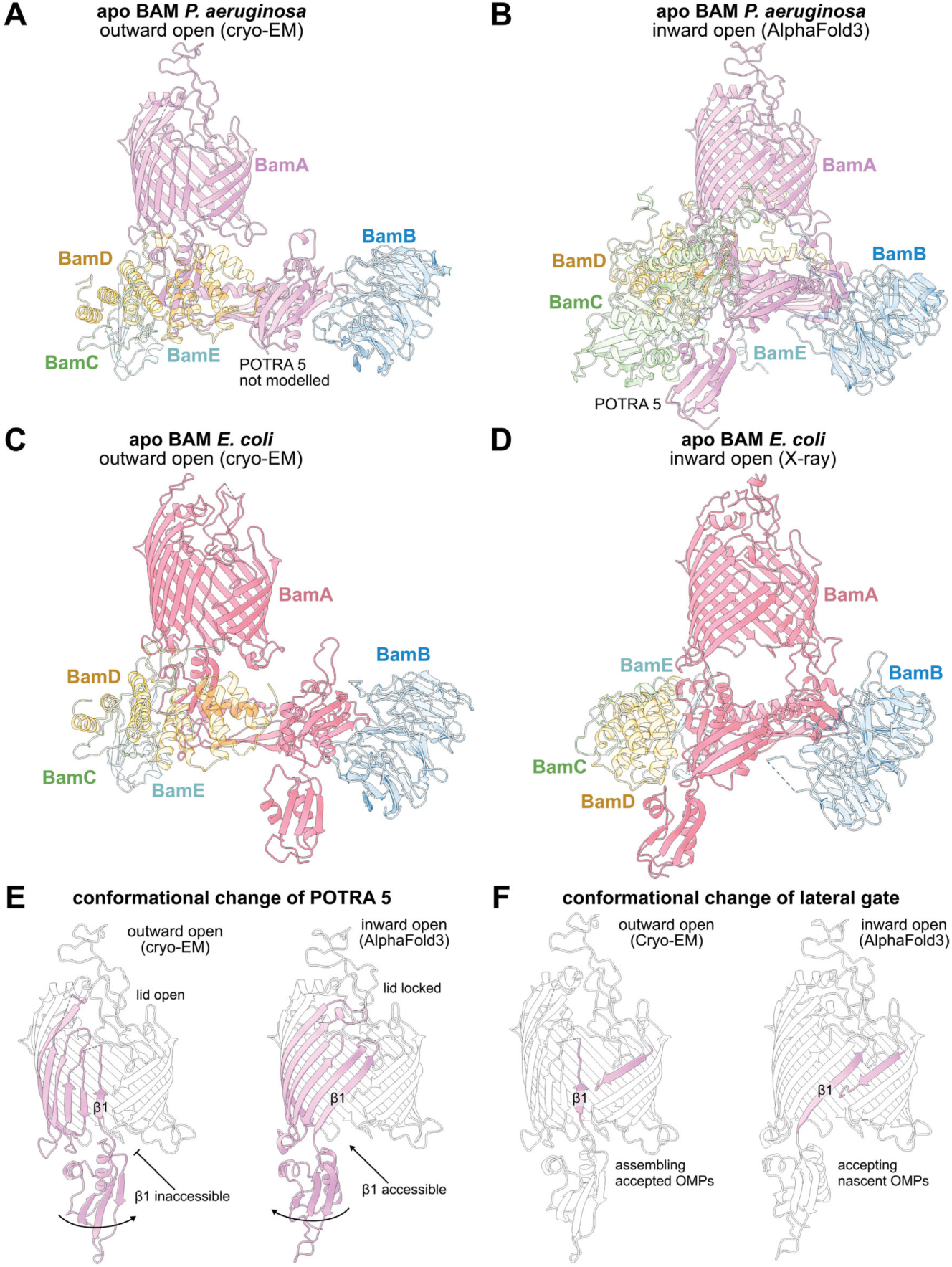
Comparison of conformational changes of P. aeruginosa BAM and E. coli BAM. (A) Cryo-EM structure of the BAM complex from *P. aeruginosa* PAO1 (pdb: 9PXG) in the lateral-gate open conformation. The POTRA 1 domain of BamA (pink) was not modelled due to poor density. BamB (blue), BamD (orange), and BamE (cyan) were built to high completeness, and BamC (green) is missing both α-grip domains due to high flexibility. (B) AlphaFold3 structure prediction of the BAM complex from *P. aeruginosa* PAO1 in the inward open conformation. Here, full-length BamC (green) is shown. (C) Cryo-EM structure of apo BAM from *E. coli* (pdb: 7NBX) in the lateral-gate open conformation with high similarity and structure completeness to *P. aeruginosa* BAM. (D) Crystal structure of apo BAM from *E. coli* in the inward open conformation. (E) Differences between the lateral-gate open (left) and inward open (right) conformations of BamA (POTRA 1-4 not shown). The N-terminal side of the β-barrel is flexible and moves POTRA 5 as it changes conformation, either blocking or allowing access to β-strand 1, while the C-terminal side of the β-barrel is fixed. (F) Differences in β-barrel N/C-terminal contacts between both conformations (POTRA 1-4 not shown). In the lateral-gate open conformation, nascent OMPs that had been accepted can be assembled at β-strand 1 (left), while nascent OMPs can be delivered to β-strand 1 in the inward open conformation (right).

**Figure S6:**
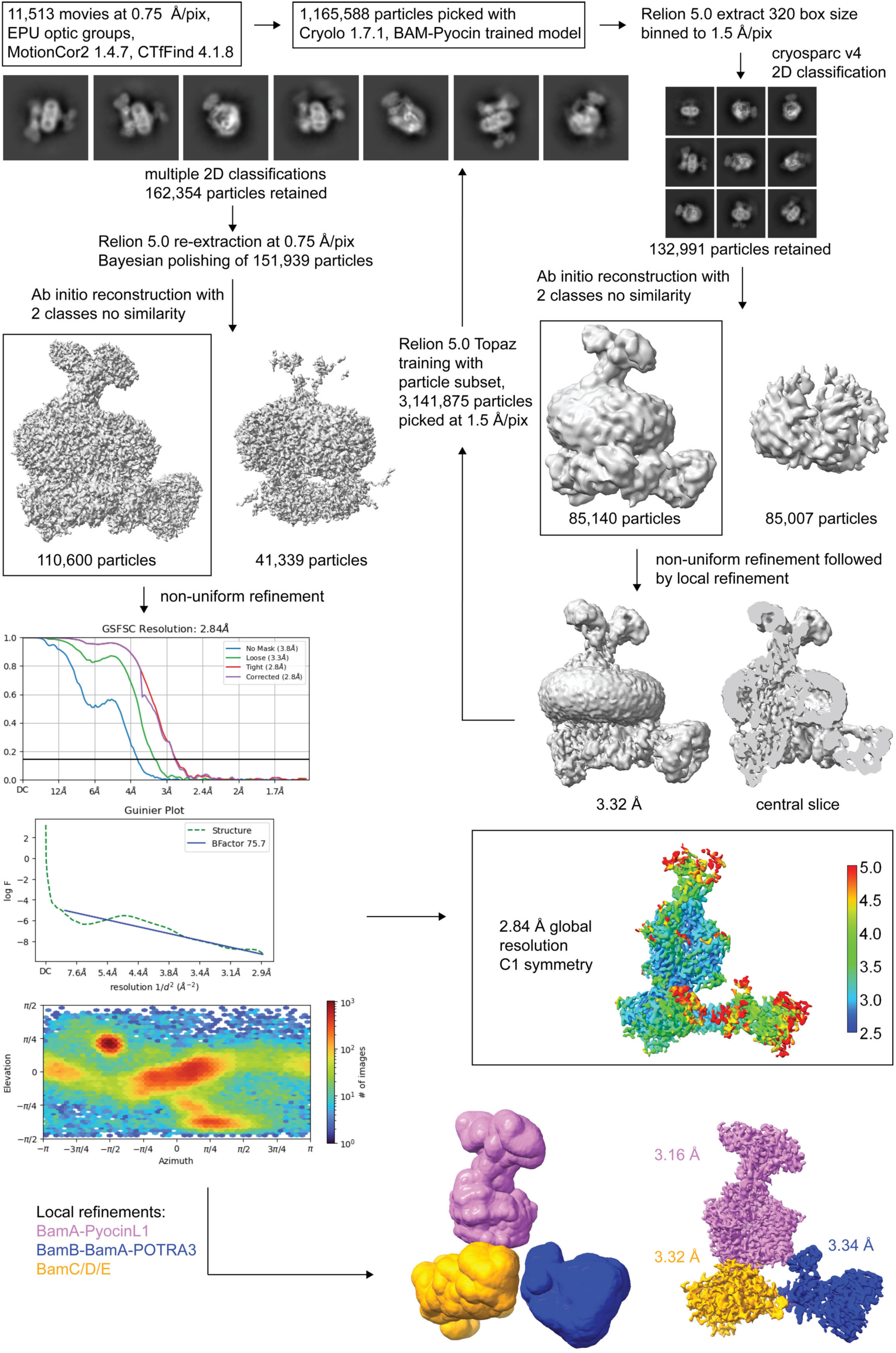
Cryo-EM SPA processing workflow of PAO1-BAM-Pyocin L1. Workflow of cryo-EM data processing of BAM from *P. aeruginosa* PAO1 in complex with Pyocin L1. Detailed information can be found in the method section.

**Figure S7:**
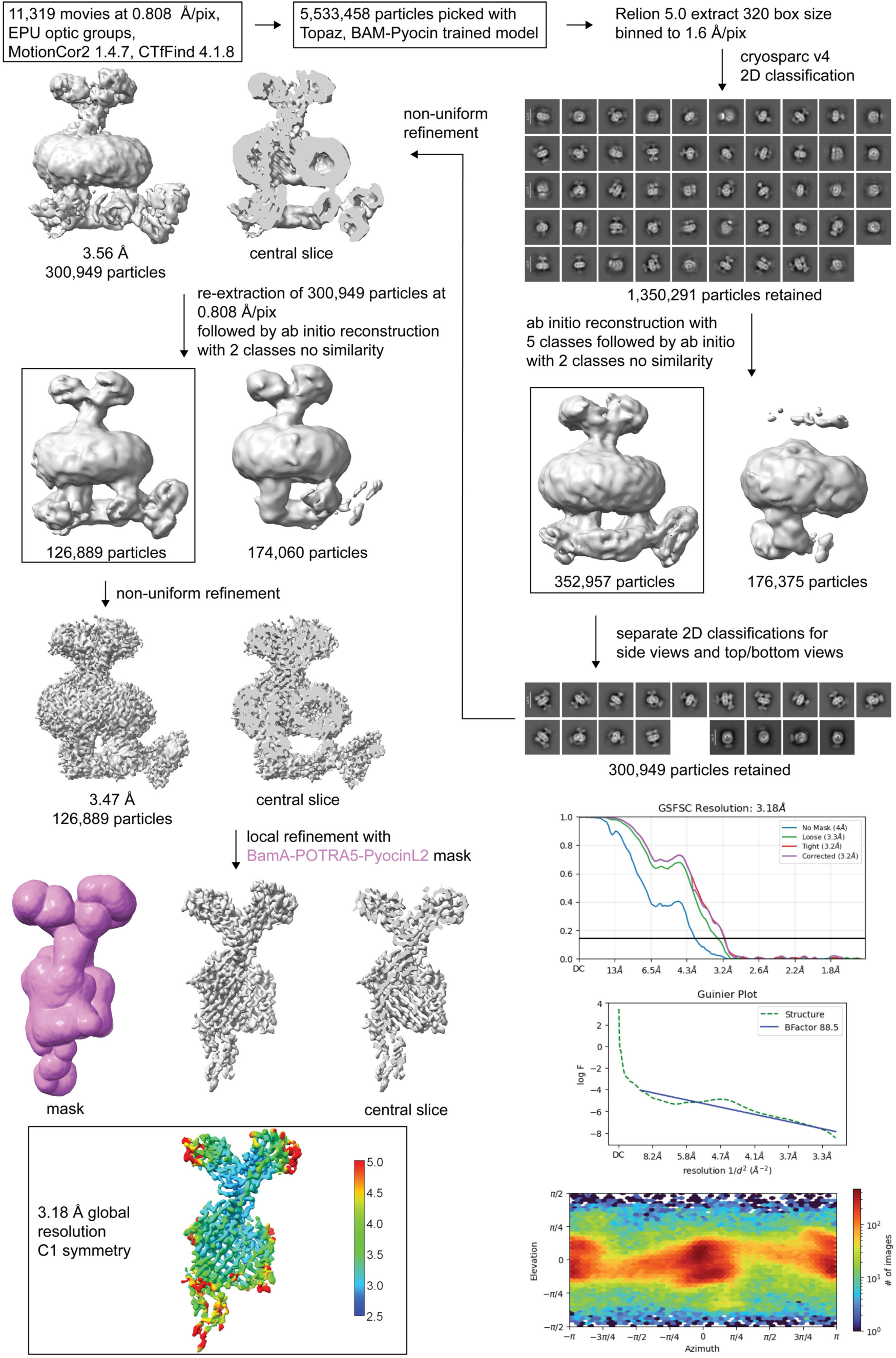
Cryo-EM SPA processing workflow of P28-BAM-Pyocin L2. Workflow of cryo-EM data processing of BAM from *P. aeruginosa* PAO1 with BamA loop 6 from *P. aeruginosa* P28 in complex with Pyocin L2. Detailed information can be found in the method section.

**Figure S8:**
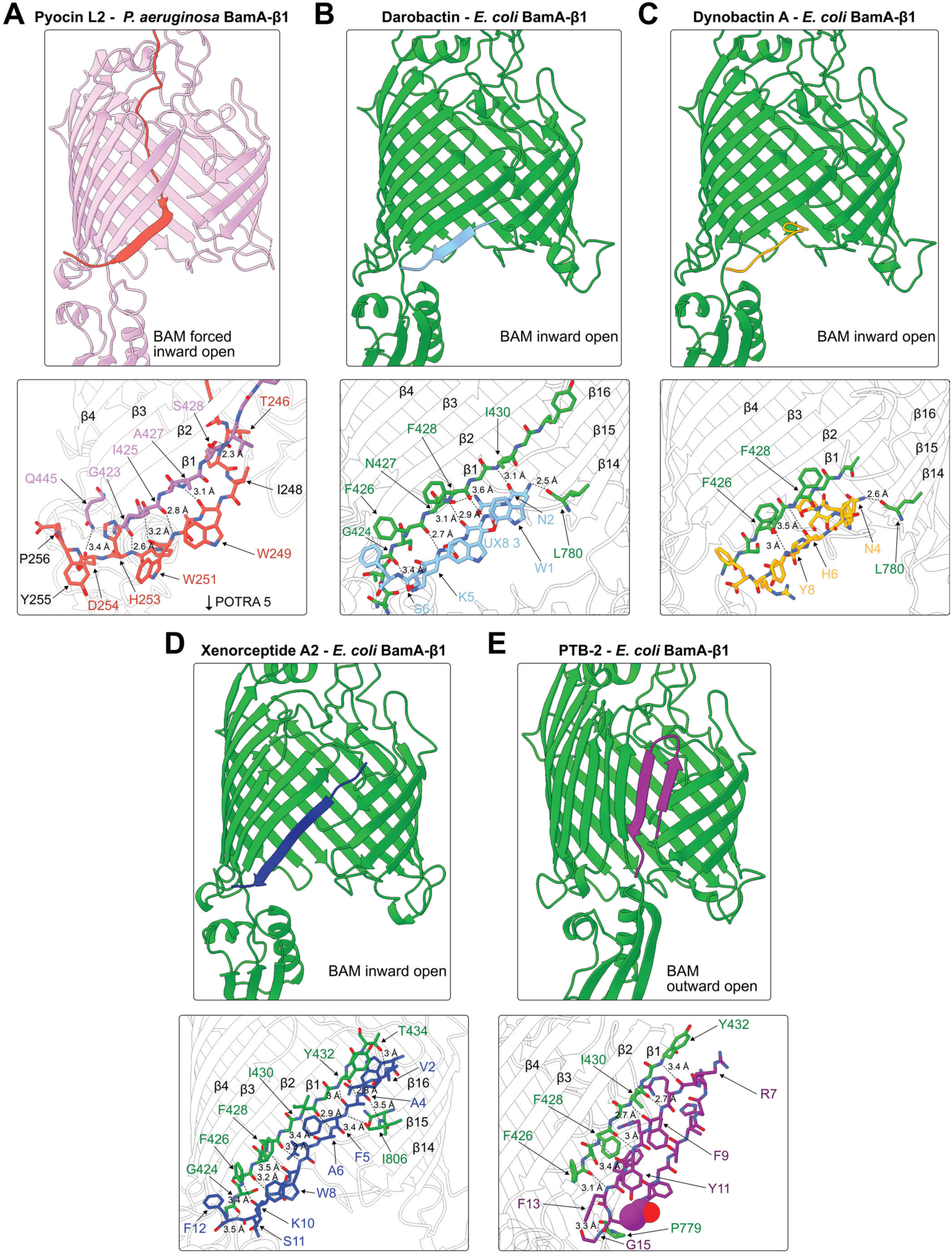
Interactions of Pyocin L2 at BamA β-strand 1 in comparison to bamabactins. (A) The C-terminal peptide of Pyocin L2 interacts with β-strand 1, forming a hybrid β-sheet with the BamA β-barrel. BamA is in the inward open conformation, but the C-terminal peptide prevents formation of contacts between the sides of the β-barrel and cannot close the extracellular loops 1-3 to prevent access to the BamA lumen. (B) The darobactin interacts with β-strand 1, forming a hybrid β-sheet with the BamA β-barrel. Here, the β-barrel forms contacts between both sides, and BamA is in an inward open conformation. (C) Dynobactin A interacts with β-strand 1 of BamA in the inward open conformation. (D) Xenorceptide A2 interacts with β-strand 1, forming a hybrid β-sheet with the BamA β-barrel and preventing contacts between both sides of the β-barrel. Xenorceptide A2 does not prevent the closing of the extracellular loops 1-3. (E) PTB-2 interacts with β-strand 1 of BamA in the lateral-gate open conformation, forming two additional β-strands of the BamA β-barrel.

**Figure S9:**
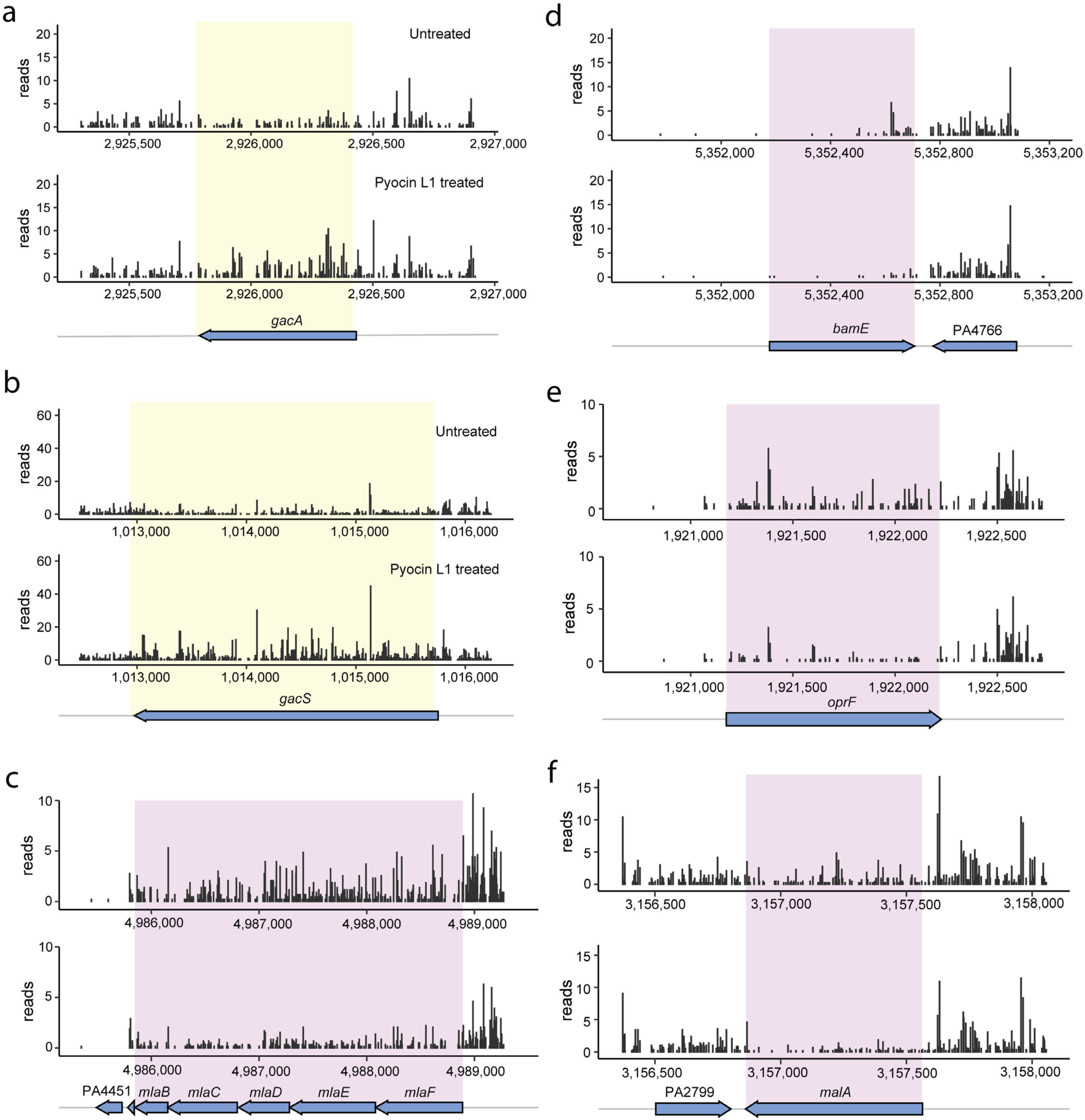
TraDIS insertion maps from *P. aeruginosa* PAO1 treated with Pyocin L1. TraDIS insertion plots of regions of the *P. aeruginosa* PAO1 genome, showing the frequency of transposon insertions by the size of the black ‘reads’ bar at that position, for untreated and Pyocin L1-treated cells from the transposon insertion library. Regions with a higher frequency of transposon insertion in the Pyocin L1-treated cells are highlighted in yellow, while those with a lower frequency of insertion are highlighted in purple. Genes encoded in these regions are shown below the insertion plots.

**Figure S10:**
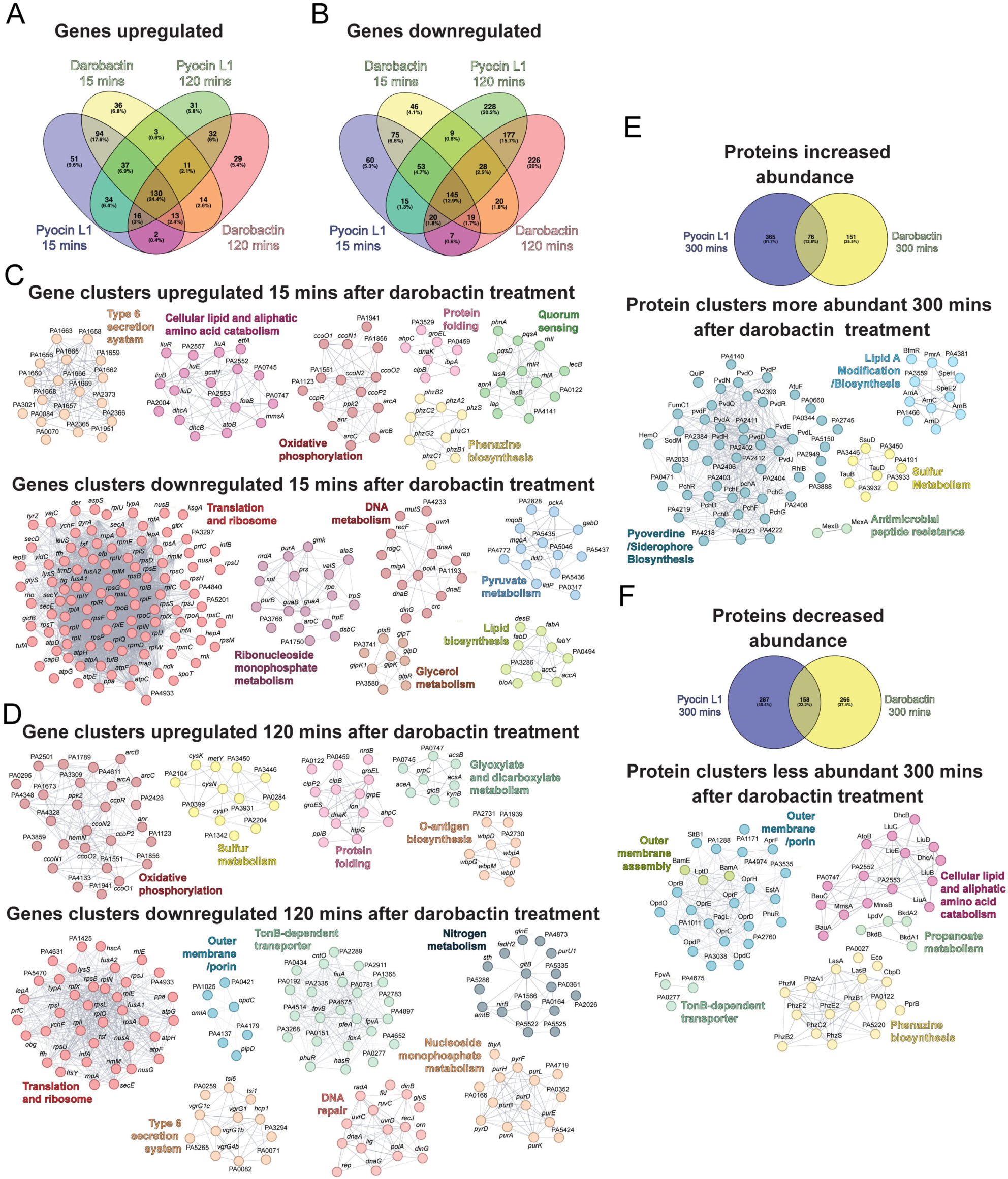
Transcriptomic and proteomic response of *P. aeruginosa* PAO1 to darobactin. (A) Venn diagrams showing the percentages of shared genes differentially expressed in *P. aeruginosa* PAO1 treated with either Pyocin L1 or darobactin at 15- and 120-minutes post-treatment. (C) and (D) STRING functional clustering analysis of genes with increased and decreased expression 15 and 120 minutes after darobactin treatment, respectively. (G) Venn diagrams showing the percentages of shared proteins with differential abundance in *P. aeruginosa* PAO1 treated with either Pyocin L1 or darobactin 300 minutes post-treatment, and STRING functional clustering analysis of proteins with increased and decreased abundance 300 minutes after darobactin treatment.

**Figure S11:**
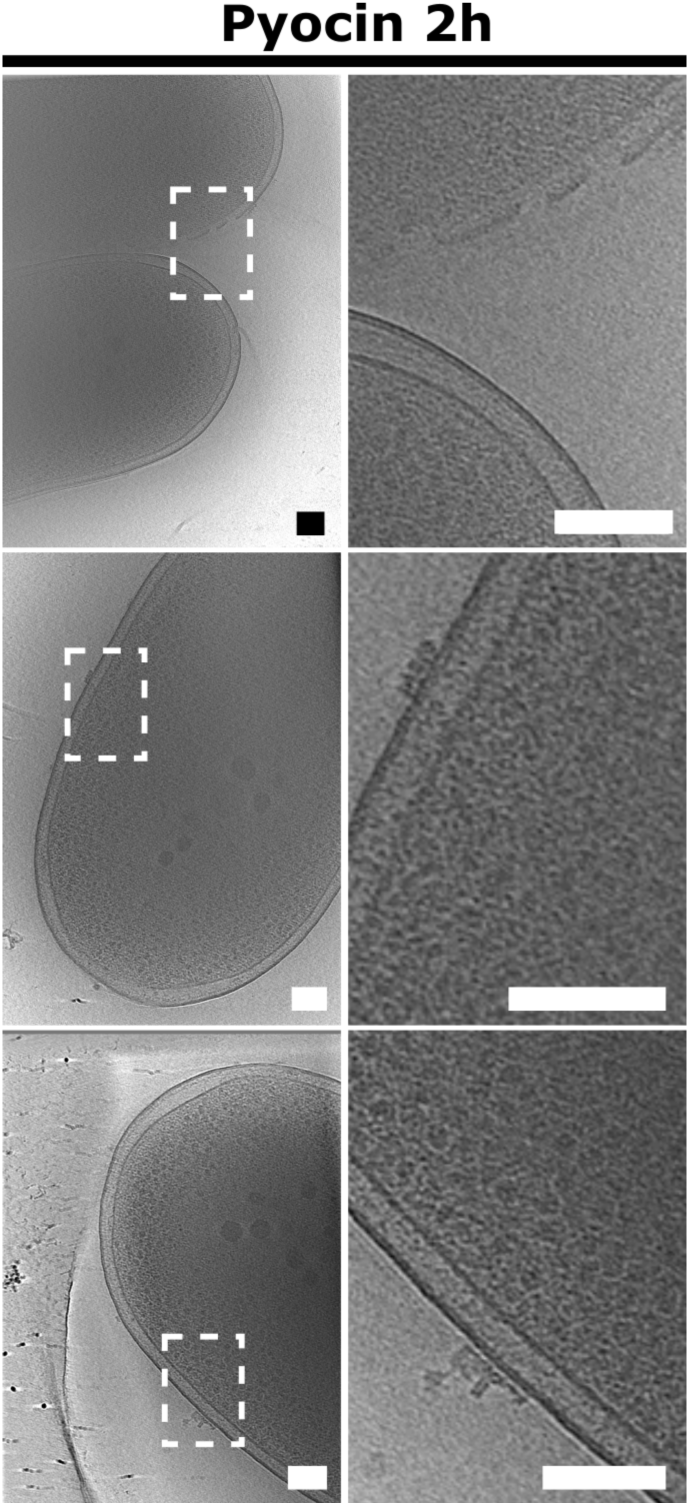
Example tomograms of *P. aeruginosa* cells treated with Pyocin L1 120 min post-treatment. 2D-slices of 3D reconstructed tomograms of three representative cells treated with Pyocin for 120 min. Zoomed regions from the left panels (white dashed box) are shown on the panels. Some cells exhibited outer membrane defects, but the majority did not display an obvious phenotype. Scale bars represent 100 nm.

**Figure S12:**
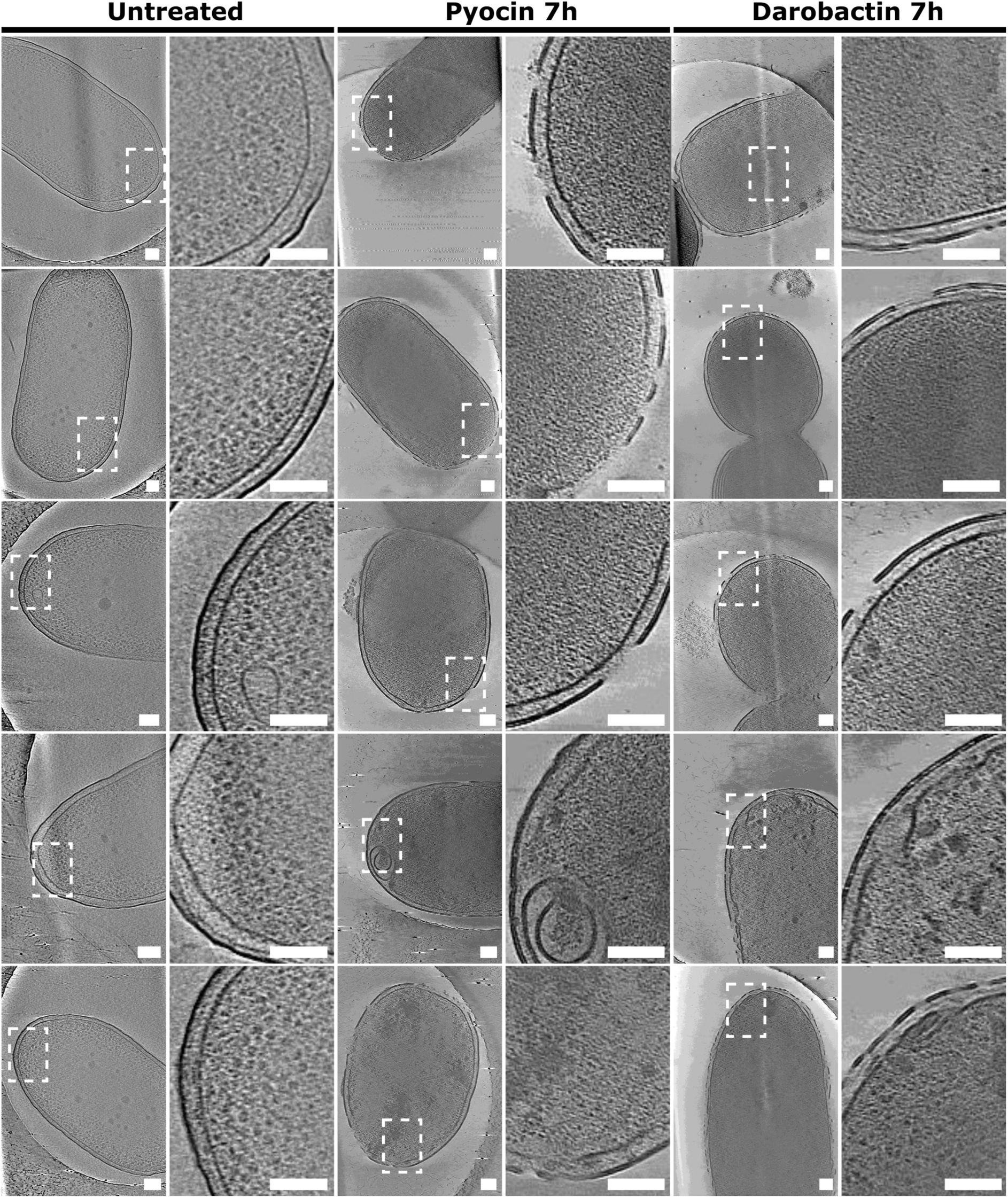
Examples of tomograms of *P. aeruginosa* cells. 2D-slices of 3D reconstructed tomograms of five representative untreated *P. aeruginosa* PAO1 cells, which show intact inner and outer membranes and an abundance of ribosomes; and cells treated with Pyocin L1 or darobactin 420 min, which show severe outer membrane perturbations and a relative paucity of ribosomes. Zoomed regions from the left panels (white dashed box) are shown on the panels and scale bars represent 100 nm.

**Figure S13:**
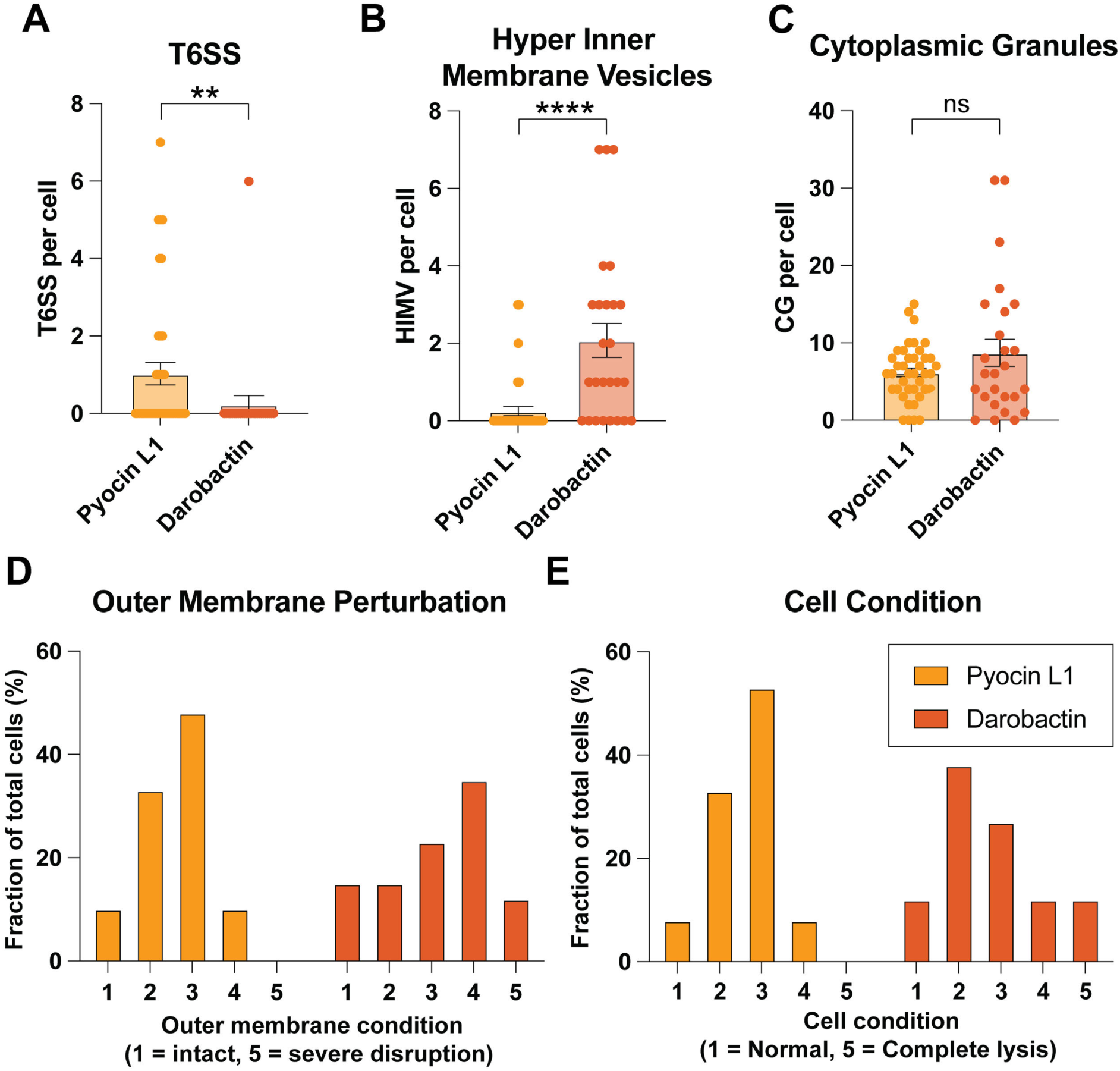
Quantification of Pyocin L1 and darobactin-treated cell features in tomograms 420 min post-treatment. Plots of the number of T6SS apparatuses (A), Hyper inner membrane vesicles (B), and Cytoplasmic granules (C) per cell for *P. aeruginosa* PAO1 treated with Pyocin L1 or darobactin. Qualitative assessment of the outer membrane perturbation (D) and general cell condition (E) for *P. aeruginosa* PAO1 treated with Pyocin L1 or darobactin. Pyocin L1-treated cells N = 40, darobactin-treated cells N = 26. Error bars = S.E.M. Significance was determined using a Mann-Whitney t-test, ** = P-value <0.01, **** <0.0001, ns = nonsignificant.

## Supplemental Tables

**Table S1:** Cryo-EM data collection, refinement, and validation statistics for apo PAO1-BAM.

|  | apo PAO1-BAM<br>9PXG | BAM<sub>PAO1</sub>-Pyocin L1<br>9PXI | BAM<sub>P28</sub>-Pyocin L2<br>9PXJ |
| --- | --- | --- | --- |
| **Data collection and processing** |  |  |  |
| Magnification | 105,000X | 105,000X | 96,000 |
| Voltage (kV) | 300 | 300 | 300 |
| Electron exposure (e-/Å<sup>2</sup>) | 60 | 60 | 53 |
| Defocus range (μm) | 0.4-1.2 | 0.4-1.2 | 0.6-1.2 |
| Pixel size (Å) | 0.75 | 0.75 | 0.808 |
| Symmetry imposed | C1 | C1 | C1 |
| Initial particle images (no.) | 3,831,721 | 3,144,875 | 5,533,458 |
| Final particle images (no.) | 121,910 | 110,600 | 126,889 |
| Map resolution (Å) | 2.75 | 2.84 | 3.18 |
| FSC threshold | 0.143 | 0.143 | 0.143 |
| Map resolution range (Å) | 2.72 – 2.98 | 2.83 – 3.17 | 3.12 – 3.18 |
| **Refinement** |  |  |  |
| Initial model used | AlphaFold3 of<br>*P. aeruginosa* BAM | apo PAO1-BAM<br>(9PXG)<br>Pyocin L1<br>(4LEA) | PAO1-BAM<br>(9PXI)<br>AlphaFold3 of<br>Pyocin L2 |
| Model resolution (Å) | 2.75 | 2.84 | 3.18 |
| FSC threshold | 0.143 | 0.143 | 0.143 |
| Model resolution range (Å) | 2.73 – 2.98 | 2.83 – 3.17 | 3.12 – 3.18 |
| Map sharpening *B* factor (Å<sup>2</sup>) | 61.4 | 75.7 | 85.5 |
| **Model composition** |  |  |  |
| Non-hydrogen atoms | 10,841 | 12,801 | 5,433 |
| Protein residues | 1,389 | 1,640 | 696 |
| Ligands | 1 | 0 | 0 |
| ***B* factors (Å<sup>2</sup>)** |  |  |  |
| Protein | 23.69 – 302.91 | 25.66 – 328.85 | 70.41 – 245.28 |
| Ligand | 73.62 | / | / |
| **R.m.s. deviations** |  |  |  |
| Bond lengths (Å) | 0.3 | 0.28 | 0.29 |
| Bond angles (°) | 0.52 | 0.51 | 0.55 |
| **Validation** |  |  |  |
| MolProbity score | 1.78 | 1.72 | 1.91 |
| Clashscore | 8 | 7 | 10 |
| Poor rotamers (%) | 0 | 0 | 0 |
| **Ramachandran plot** |  |  |  |
| Favored (%) | 95 | 96 | 93 |
| Allowed (%) | 5 | 4 | 7 |
| Disallowed (%) | 0 | 0 | 0 |

**Table S2: TraDIS data table (provided as Excel sheet)**

**Table S3:**
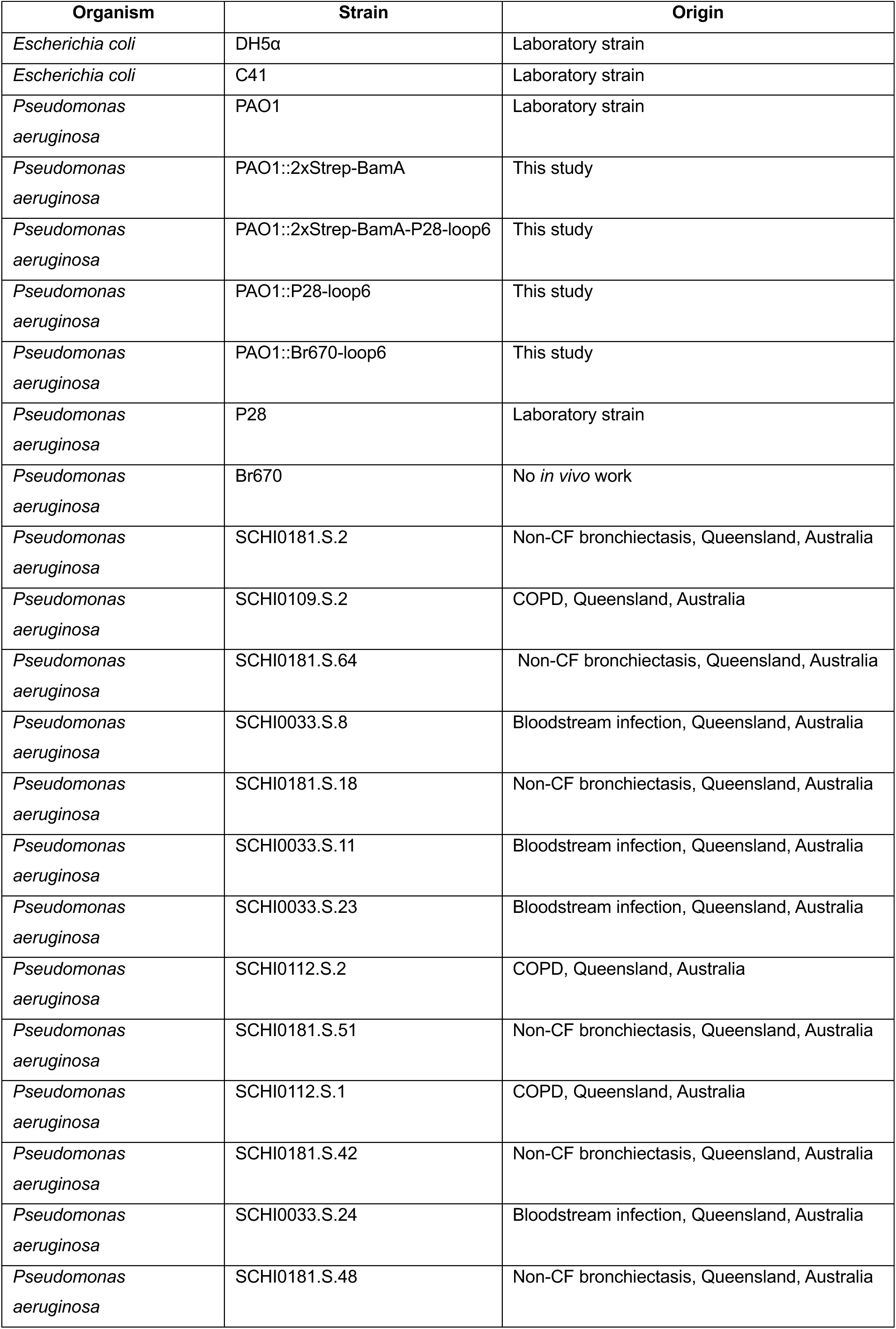
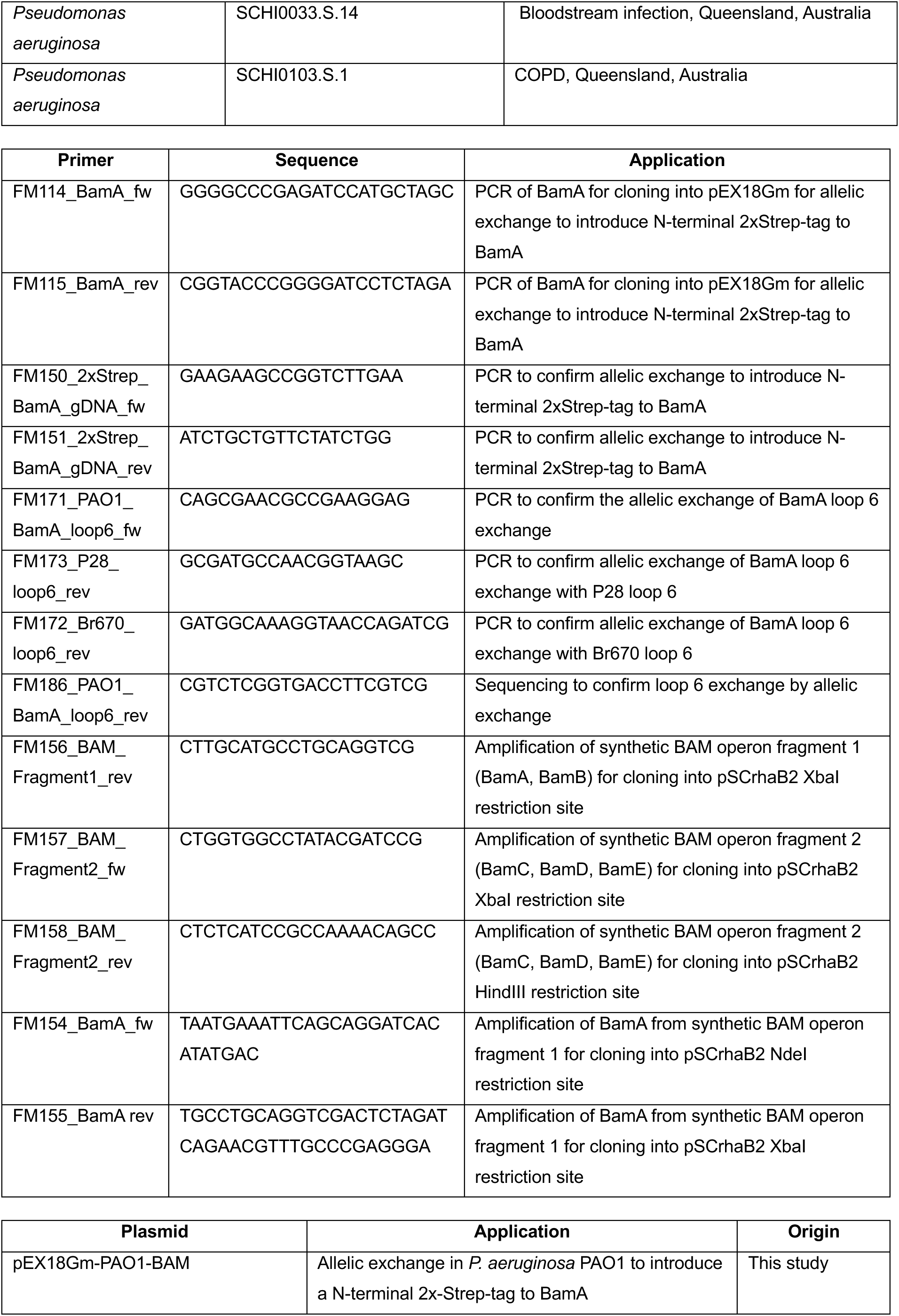
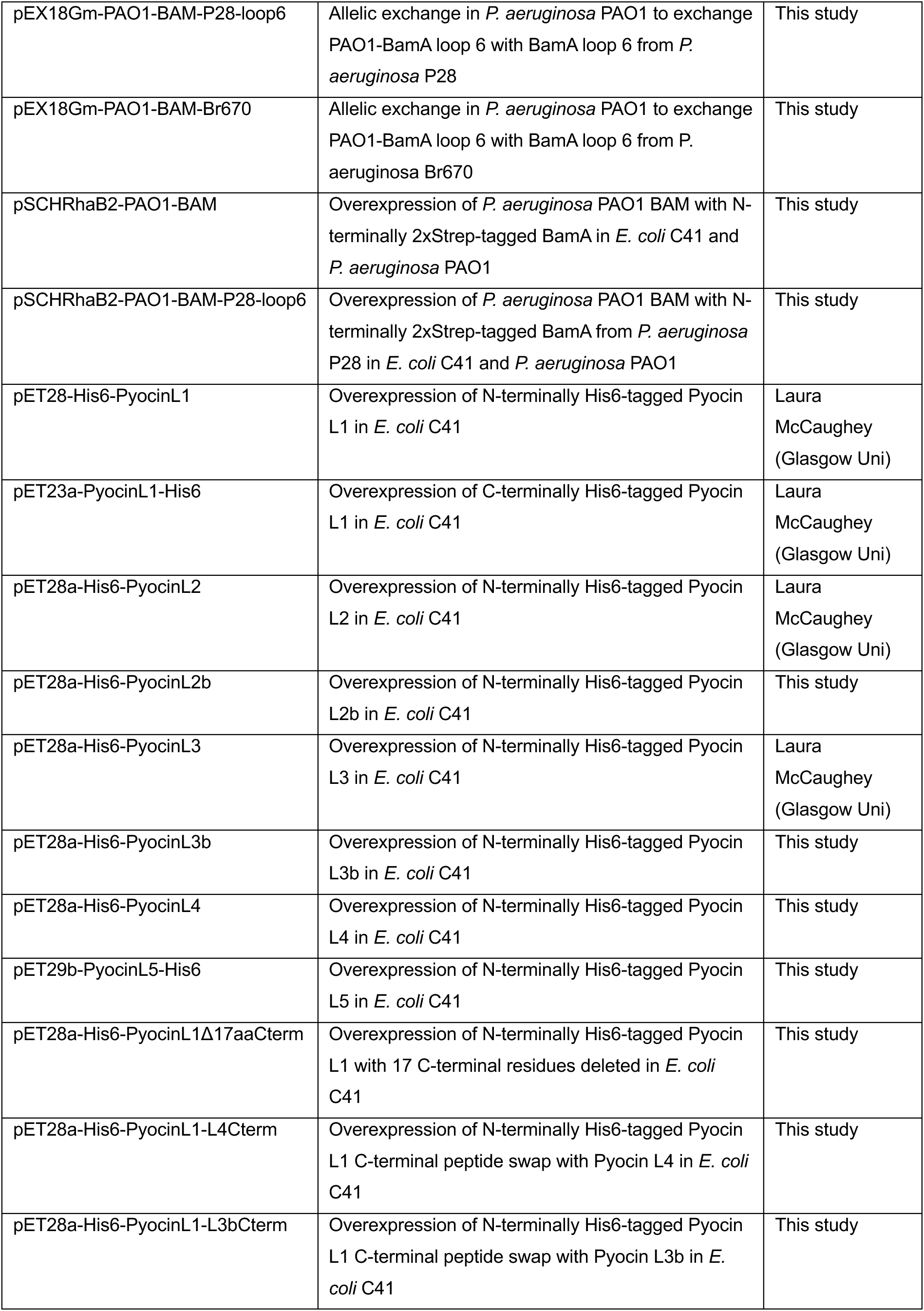
Strains, primers, and plasmids used in this study.

**Table S4:** Molecular dynamics simulation setups.

| Setup | State | Starting structure | Presence of pyocin | Modelled regions, residue IDs | Number of atoms | Simulation time, ~μs |  |  |
| --- | --- | --- | --- | --- | --- | --- | --- | --- |
|  |  |  |  |  |  | Replica 1 | Replica 2 | Replica 3 |
| 1 | Pre-inhibited | 9PXI | + | BamA: 469-472<br>Pyocin: 261-276 | 386,051 | 1 | 1.01 | 1 |
| 2 | Pre-inhibited | 9PXI | - | BamA: 469-472 | 387,583 | 0.95 | 1.06 | 1.07 |
| 3 | Fully inhibited | 9PXJ | + | BamA: 382-453, 701-729, 746-756 | 384,915 | 1.02 | 1.14 | 1.13 |
| 4 | Fully inhibited | 9PXJ | - |  | 386,399 | 1.10 | 1.11 | 1.07 |

## Supplementary Movies

**Movie S1: A morph of conformational changes between the Cryo-EM structure and AlphaFold2 mode of BAM_PAO1_**

**Movie S2: Key elements of the structure of the BAM_PAO1_-Pyocin L1 complex**

**Movie S3: Molecular dynamics simulation of the pre-inhibited PAO1-BAM-Pyocin L1 complex with modelled C-terminal tail**

**Movie S4: Key elements of the structure of the BAM_PAO1_-Pyocin L2 complex**

**Movie S5: Molecular dynamics simulation of the fully inhibited P28-BAM-Pyocin L2 complex**

**Movie S6: A morph capturing the conformational changes of Pyocin L1 and BAM_PAO1_ during the formation of the BAM_PAO1_-Pyocin L1 inhibitor complex.**

**Movie S7: Tomography analysis of untreated *P. aeruginosa* PAO1 cells after 420 minutes of growth in LB broth.**

**Movie S8: Tomography analysis of *P. aeruginosa* PAO1 treated with Pyocin L1 after 420 minutes of growth in LB broth.**

**Movie S9: Tomography analysis of *P. aeruginosa* PAO1 treated with darobactin after 420 minutes of growth in LB broth.**

**Supplementary dataset 1: *P. aeruginosa* BamA sequences**

**Supplementary dataset 2: Processed TraDIS data and analysis**

**Supplementary dataset 3: Processed transcriptomics data and STRING analysis**

**Supplementary dataset 4: Processed proteomics data and STRING analysis**

## References

1 Qin, S. et al. Pseudomonas aeruginosa: pathogenesis, virulence factors, antibiotic resistance, interaction with host, technology advances and emerging therapeutics. Signal transduction and targeted therapy 7, 199 (2022).

2 Abdulhak, A. et al. Multidrug-resistant Pseudomonas aeruginosa in immunocompromised cancer patients: epidemiology, antimicrobial resistance, and virulence factors. BMC Infectious Diseases 25, 804 (2025).

3 Reynolds, D. & Kollef, M. The epidemiology and pathogenesis and treatment of Pseudomonas aeruginosa infections: an update. Drugs 81, 2117–2131 (2021).

4 Pang, Z., Raudonis, R., Glick, B. R., Lin, T.-J. & Cheng, Z. Antibiotic resistance in Pseudomonas aeruginosa: mechanisms and alternative therapeutic strategies. Biotechnology advances 37, 177–192 (2019).

5 Lister, P. D., Wolter, D. J. & Hanson, N. D. Antibacterial-resistant Pseudomonas aeruginosa: clinical impact and complex regulation of chromosomally encoded resistance mechanisms. Clinical microbiology reviews 22, 582–610 (2009).

6 Organization, W. H. WHO bacterial priority pathogens list, 2024: bacterial pathogens of public health importance, to guide research, development, and strategies to prevent and control antimicrobial resistance. (World Health Organization, 2024).

7 Storek, K. M., Sun, D. & Rutherford, S. T. Inhibitors targeting BamA in gram-negative bacteria. Biochimica et Biophysica Acta (BBA)-Molecular Cell Research 1871, 119609 (2024).

8 Pahil, K. S. et al. A new antibiotic traps lipopolysaccharide in its intermembrane transporter. Nature 625, 572–577 (2024). 10.1038/s41586-023-06799-7

9 Braun, M. & Silhavy, T. J. Imp/OstA is required for cell envelope biogenesis in *Escherichia coli*. Molecular microbiology 45, 1289–1302 (2002).

10 Tashiro, Y. et al. Opr86 Is Essential for Viability and Is a Potential Candidate for a Protective Antigen against Biofilm Formation by *Pseudomonas aeruginosa*. Journal of Bacteriology 190, 3969–3978 (2008). 10.1128/jb.02004-07

11 Voulhoux, R., Bos, M. P., Geurtsen, J., Mols, M. & Tommassen, J. Role of a highly conserved bacterial protein in outer membrane protein assembly. Science 299, 262–265 (2003).

12 McCaughey, L. C. et al. Lectin-like bacteriocins from *Pseudomonas* spp. utilise D-rhamnose containing lipopolysaccharide as a cellular receptor. PLoS pathogens 10, e1003898 (2014).

13 Ghequire, M. G., Swings, T., Michiels, J., Buchanan, S. K. & De Mot, R. Hitting with a BAM: selective killing by lectin-like bacteriocins. MBio 9, 10.1128/mbio.02138-02117 (2018).

14 Ghequire, M. G., Öztürk, B. & De Mot, R. Lectin-like bacteriocins. Frontiers in Microbiology 9, 2706 (2018).

15 McCaughey, L. C., Ritchie, N. D., Douce, G. R., Evans, T. J. & Walker, D. Efficacy of species-specific protein antibiotics in a murine model of acute Pseudomonas aeruginosa lung infection. Scientific reports 6, 30201 (2016).

16 Ghequire, M. G. K. et al. Structural Determinants for Activity and Specificity of the Bacterial Toxin LlpA. PLOS Pathogens 9, e1003199 (2013). 10.1371/journal.ppat.1003199

17 Ghequire, M. G. et al. O serotype-independent susceptibility of Pseudomonas aeruginosa to lectin-like pyocins. Microbiologyopen 3, 875–884 (2014). 10.1002/mbo3.210

18 Noinaj, N. et al. Structural insight into the biogenesis of β-barrel membrane proteins. Nature 501, 385–390 (2013).

19 Struyvé, M., Moons, M. & Tommassen, J. Carboxy-terminal phenylalanine is essential for the correct assembly of a bacterial outer membrane protein. Journal of molecular biology 218, 141–148 (1991).

20 Wu, T. et al. Identification of a multicomponent complex required for outer membrane biogenesis in Escherichia coli. Cell 121, 235–245 (2005).

21 Shen, C. et al. Structural basis of BAM-mediated outer membrane β-barrel protein assembly. Nature 617, 185–193 (2023).

22 Karruli, A. et al. Evidence-Based Treatment of Pseudomonas aeruginosa Infections: A Critical Reappraisal. Antibiotics (Basel) 12 (2023). 10.3390/antibiotics12020399

23 Mackay, B., Parcell, B. J., Shirran, S. L. & Coote, P. J. Carbapenem-Only Combination Therapy against Multi-Drug Resistant Pseudomonas aeruginosa: Assessment of In Vitro and In Vivo Efficacy and Mode of Action. Antibiotics (Basel) 11 (2022). 10.3390/antibiotics11111467

24 Farrell, D. J., Flamm, R. K., Sader, H. S. & Jones, R. N. Antimicrobial activity of ceftolozane-tazobactam tested against Enterobacteriaceae and Pseudomonas aeruginosa with various resistance patterns isolated in U.S. Hospitals (2011-2012). Antimicrob Agents Chemother 57, 6305–6310 (2013). 10.1128/aac.01802-13

25 Nichols, W. W. et al. Ceftazidime-Avibactam Susceptibility Breakpoints against Enterobacteriaceae and Pseudomonas aeruginosa. Antimicrob Agents Chemother 62 (2018). 10.1128/aac.02590-17

26 Madden, D. E. et al. Keeping up with the pathogens: improved antimicrobial resistance detection and prediction from Pseudomonas aeruginosa genomes. Genome Medicine 16, 78 (2024). 10.1186/s13073-024-01346-z

27 Ghequire, M. G., Li, W., Proost, P., Loris, R. & De Mot, R. Plant lectin-like antibacterial proteins from phytopathogens Pseudomonas syringae and Xanthomonas citri. Environmental microbiology reports 4, 373–380 (2012).

28 Parret, A. H., Wyns, L., De Mot, R. & Loris, R. Overexpression, purification and crystallization of bacteriocin LlpA from Pseudomonas sp. BW11M1. Biological Crystallography 60, 1922–1924 (2004).

29 Iadanza, M. G. et al. Lateral opening in the intact β-barrel assembly machinery captured by cryo-EM. Nature Communications 7, 12865 (2016). 10.1038/ncomms12865

30 Gu, Y. et al. Structural basis of outer membrane protein insertion by the BAM complex. Nature 531, 64–69 (2016). 10.1038/nature17199

31 Doyle, M. T. & Bernstein, H. D. Bacterial outer membrane proteins assemble via asymmetric interactions with the BamA β-barrel. Nature Communications 10, 3358 (2019). 10.1038/s41467-019-11230-9

32 Doyle, M. T. et al. Cryo-EM structures reveal multiple stages of bacterial outer membrane protein folding. Cell 185, 1143–1156.e1113 (2022). 10.1016/j.cell.2022.02.016

33 Tomasek, D. et al. Structure of a nascent membrane protein as it folds on the BAM complex. Nature 583, 473–478 (2020).

34 Fenn, K. L. et al. Outer membrane protein assembly mediated by BAM-SurA complexes. Nature Communications 15, 7612 (2024). 10.1038/s41467-024-51358-x

35 Kaur, H. et al. The antibiotic darobactin mimics a β-strand to inhibit outer membrane insertase. Nature 593, 125–129 (2021). 10.1038/s41586-021-03455-w

36 Miller, R. D. et al. Computational identification of a systemic antibiotic for Gram-negative bacteria. Nature Microbiology 7, 1661–1672 (2022). 10.1038/s41564-022-01227-4

37 Modaresi, S. M. et al. Antibiotics that Kill Gram-negative Bacteria by Restructuring the Outer Membrane Protein BamA. bioRxiv, 2024.2012.2016.628070 (2024). 10.1101/2024.12.16.628070

38 Imai, Y. et al. A new antibiotic selectively kills Gram-negative pathogens. Nature 576, 459–464 (2019). 10.1038/s41586-019-1791-1

39 Lehner, P. A. et al. Architecture and conformational dynamics of the BAM-SurA holo insertase complex. Science Advances 11, eads6094 (2025). doi:10.1126/sciadv.ads6094

40 Barquist, L. et al. The TraDIS toolkit: sequencing and analysis for dense transposon mutant libraries. Bioinformatics 32, 1109–1111 (2016).

41 Short, F. L. et al. Transposon insertion sequencing reveals novel hypermutator genes in Acinetobacter baumannii. mBio, e00966–00925 (2025).

42 Song, H., Li, Y. & Wang, Y. Two-component system GacS/GacA, a global response regulator of bacterial physiological behaviors. Engineering Microbiology 3, 100051 (2023).

43 Khandekar, S. et al. The putative de-N-acetylase DnpA contributes to intracellular and biofilm-associated persistence of Pseudomonas aeruginosa exposed to fluoroquinolones. Frontiers in microbiology 9, 1455 (2018).

44 Ruiz, N., Davis, R. M. & Kumar, S. YhdP, TamB, and YdbH are redundant but essential for growth and lipid homeostasis of the Gram-negative outer membrane. MBio 12, e02714–02721 (2021).

45 Thong, S. et al. Defining key roles for auxiliary proteins in an ABC transporter that maintains bacterial outer membrane lipid asymmetry. Elife 5, e19042 (2016).

46 Abellón-Ruiz, J. et al. Structural basis for maintenance of bacterial outer membrane lipid asymmetry. Nature Microbiology 2, 1616–1623 (2017). 10.1038/s41564-017-0046-x

47 Nikaido, H. Molecular basis of bacterial outer membrane permeability revisited. Microbiology and molecular biology reviews 67, 593–656 (2003).

48 Paulsson, M. et al. Peptidoglycan-Binding Anchor Is a Pseudomonas aeruginosa OmpA Family Lipoprotein With Importance for Outer Membrane Vesicles, Biofilms, and the Periplasmic Shape. Frontiers in Microbiology Volume 12 **-** 2021 (2021). 10.3389/fmicb.2021.639582

49 Lim Jr, A., et al. Molecular and immunological characterization of OprL, the 18 kDa outer-membrane peptidoglycan-associated lipoprotein (PAL) of Pseudomonas aeruginosa. Microbiology 143, 1709–1716 (1997).

50 Kadokura, H., Katzen, F. & Beckwith, J. Protein disulfide bond formation in prokaryotes. Annual review of biochemistry 72, 111–135 (2003).

51 Schurr, M., Yu, H., Martinez-Salazar, J., Boucher, J. & Deretic, V. Control of AlgU, a member of the sigma E-like family of stress sigma factors, by the negative regulators MucA and MucB and Pseudomonas aeruginosa conversion to mucoidy in cystic fibrosis. Journal of bacteriology 178, 4997–5004 (1996).

52 Wood, L. F. & Ohman, D. E. Use of cell wall stress to characterize σ22 (AlgT/U) activation by regulated proteolysis and its regulon in Pseudomonas aeruginosa. Molecular microbiology 72, 183–201 (2009).

53 LaBauve, A. E. & Wargo, M. J. Growth and laboratory maintenance of Pseudomonas aeruginosa. Curr Protoc Microbiol Chapter 6, Unit 6E.1. (2012). 10.1002/9780471729259.mc06e01s25

54 Silale, A. & van den Berg, B. TonB-Dependent Transport Across the Bacterial Outer Membrane. Annu Rev Microbiol 77, 67–88 (2023). 10.1146/annurev-micro-032421-111116

55 Chevalier, S. et al. Structure, function and regulation of Pseudomonas aeruginosa porins. FEMS Microbiology Reviews 41, 698–722 (2017). 10.1093/femsre/fux020

56 Basler, M., Ho, B. T. & Mekalanos, J. J. Tit-for-tat: type VI secretion system counterattack during bacterial cell-cell interactions. Cell 152, 884–894 (2013).

57 George, M., Narayanan, S., Tejada-Arranz, A., Plack, A. & Basler, M. Initiation of H1-T6SS dueling between Pseudomonas aeruginosa. mBio 15, e00355–00324 (2024). 10.1128/mbio.00355-24

58 Sun, D. et al. The discovery and structural basis of two distinct state-dependent inhibitors of BamA. Nature Communications 15, 8718 (2024). 10.1038/s41467-024-52512-1

59 Zgurskaya, H. I., López, C. A. & Gnanakaran, S. Permeability barrier of Gram-negative cell envelopes and approaches to bypass it. ACS infectious diseases 1, 512–522 (2015).

60 Maher, C. & Hassan, K. A. The Gram-negative permeability barrier: tipping the balance of the in and the out. MBio 14, e01205–01223 (2023).

61 Bryant, J. A. et al. (eLife Sciences Publications, Ltd, 2024).

62 Hews, C. L., Cho, T., Rowley, G. & Raivio, T. L. Maintaining integrity under stress: envelope stress response regulation of pathogenesis in gram-negative bacteria. Frontiers in cellular and infection microbiology 9, 313 (2019).

63 Davies, M. R. et al. Atlas of group A streptococcal vaccine candidates compiled using large-scale comparative genomics. Nat Genet 51, 1035–1043 (2019). 10.1038/s41588-019-0417-8

64 Sievers, F. et al. Fast, scalable generation of high-quality protein multiple sequence alignments using Clustal Omega. Molecular Systems Biology 7, 539 (2011). 10.1038/msb.2011.75

65 Edgar, R. C. MUSCLE: multiple sequence alignment with high accuracy and high throughput. Nucleic Acids Res 32, 1792–1797 (2004). 10.1093/nar/gkh340

66 Lemoine, F., et al. NGPhylogeny.fr: new generation phylogenetic services for non-specialists. Nucleic Acids Res 47, W260-W265 (2019). 10.1093/nar/gkz303

67 Dress, A. W. et al. Noisy: identification of problematic columns in multiple sequence alignments. Algorithms Mol Biol 3, 7 (2008). 10.1186/1748-7188-3-7

68 Minh, B. Q. et al. IQ-TREE 2: New Models and Efficient Methods for Phylogenetic Inference in the Genomic Era. Mol Biol Evol 37, 1530–1534 (2020). 10.1093/molbev/msaa015

69 Letunic, I. & Bork, P. Interactive Tree of Life (iTOL) v6: recent updates to the phylogenetic tree display and annotation tool. Nucleic Acids Res 52, W78–W82 (2024). 10.1093/nar/gkae268

70 Pettersen, E. F. et al. UCSF ChimeraX: Structure visualization for researchers, educators, and developers. Protein Sci 30, 70–82 (2021). 10.1002/pro.3943

71 Motohashi, K. A simple and efficient seamless DNA cloning method using SLiCE from Escherichia coli laboratory strains and its application to SLiP site-directed mutagenesis. BMC Biotechnology 15, 47 (2015). 10.1186/s12896-015-0162-8

72 Hmelo, L. R. et al. Precision-engineering the Pseudomonas aeruginosa genome with two-step allelic exchange. Nature Protocols 10, 1820–1841 (2015). 10.1038/nprot.2015.115

73 Munder, F., Voutsinos, M., Hantke, K., Venugopal, H. & Grinter, R. High-affinity PQQ import is widespread in Gram-negative bacteria. Science Advances 11, eadr2753 (2025). doi:10.1126/sciadv.adr2753

74 Zheng, S. Q. et al. MotionCor2: anisotropic correction of beam-induced motion for improved cryo-electron microscopy. Nature methods 14, 331–332 (2017).

75 Rohou, A. & Grigorieff, N. CTFFIND4: Fast and accurate defocus estimation from electron micrographs. Journal of structural biology 192, 216–221 (2015).

76 Burt, A. et al. An image processing pipeline for electron cryo-tomography in RELION-5. FEBS Open Bio 14, 1788–1804 (2024).

77 Bepler, T. et al. Positive-unlabeled convolutional neural networks for particle picking in cryo-electron micrographs. Nature methods 16, 1153–1160 (2019).

78 Punjani, A., Rubinstein, J. L., Fleet, D. J. & Brubaker, M. A. cryoSPARC: algorithms for rapid unsupervised cryo-EM structure determination. Nature methods 14, 290–296 (2017).

79 Asarnow, D., Palovcak, E. & Cheng, Y. UCSF pyem v0. 5. Zenodo 10.5281/zenodo 3576630, 2019 (2019).

80 Zivanov, J., Nakane, T. & Scheres, S. H. A Bayesian approach to beam-induced motion correction in cryo-EM single-particle analysis. IUCrJ 6, 5–17 (2019).

81 Meng, E. C. et al. UCSF ChimeraX: Tools for structure building and analysis. Protein Science 32, e4792 (2023).

82 Pogozheva, I. D. et al. Comparative Molecular Dynamics Simulation Studies of Realistic Eukaryotic, Prokaryotic, and Archaeal Membranes. Journal of Chemical Information and Modeling 62, 1036–1051 (2022). 10.1021/acs.jcim.1c01514

83 Jo, S., Kim, T., Iyer, V. G. & Im, W. CHARMM-GUI: A web-based graphical user interface for CHARMM. Journal of Computational Chemistry 29, 1859–1865 (2008). 10.1002/jcc.20945

84 Phillips, J. C. et al. Scalable molecular dynamics on CPU and GPU architectures with NAMD. J Chem Phys 153, 044130 (2020). 10.1063/5.0014475

85 Abraham, M. J. et al. GROMACS: High performance molecular simulations through multi-level parallelism from laptops to supercomputers. SoftwareX 1-2, 19–25 (2015). 10.1016/j.softx.2015.06.001

86 Huang, J. et al. CHARMM36m: an improved force field for folded and intrinsically disordered proteins. Nature Methods 14, 71–73 (2017). 10.1038/nmeth.4067

87 Klauda, J. B. et al. Update of the CHARMM all-atom additive force field for lipids: validation on six lipid types. J Phys Chem B 114, 7830–7843 (2010). 10.1021/jp101759q

88 Beglov, D. & Roux, B. Finite representation of an infinite bulk system: Solvent boundary potential for computer simulations. The Journal of Chemical Physics 100, 9050–9063 (1994). 10.1063/1.466711

89 Best, R. B. et al. Optimization of the additive CHARMM all-atom protein force field targeting improved sampling of the backbone φ, ψ and side-chain χ(1) and χ(2) dihedral angles. J Chem Theory Comput 8, 3257–3273 (2012). 10.1021/ct300400x

90 Jorgensen, W. L., Chandrasekhar, J., Madura, J. D., Impey, R. W. & Klein, M. L. Comparison of simple potential functions for simulating liquid water. The Journal of Chemical Physics 79, 926–935 (1983). 10.1063/1.445869

91 Venable, R. M., Luo, Y., Gawrisch, K., Roux, B. & Pastor, R. W. Simulations of anionic lipid membranes: development of interaction-specific ion parameters and validation using NMR data. J Phys Chem B 117, 10183–10192 (2013). 10.1021/jp401512z

92 Hess, B., Bekker, H., Berendsen, H. J. C. & Fraaije, J. G. E. M. LINCS: A linear constraint solver for molecular simulations. Journal of Computational Chemistry 18, 1463–1472 (1997). 10.1002/(SICI)1096-987X(199709)18:12<1463::AID-JCC4>3.0.CO;2-H

93 Verlet, L. Computer "Experiments" on Classical Fluids. I. Thermodynamical Properties of Lennard-Jones Molecules. Physical Review 159, 98–103 (1967). 10.1103/physrev.159.98

94 Darden, T., York, D. & Pedersen, L. Particle mesh Ewald: An *N* log *N* method for Ewald sums in large systems. The Journal of Chemical Physics 98, 10089–10092 (1993). 10.1063/1.464397

95 Bussi, G., Donadio, D. & Parrinello, M. Canonical sampling through velocity rescaling. The Journal of Chemical Physics 126, 014101 (2007). 10.1063/1.2408420

96 Berendsen, H. J. C., Postma, J. P. M., Van Gunsteren, W. F., Dinola, A. & Haak, J. R. Molecular dynamics with coupling to an external bath. The Journal of Chemical Physics 81, 3684–3690 (1984). 10.1063/1.448118

97 Nosé, S. A unified formulation of the constant temperature molecular dynamics methods. The Journal of Chemical Physics 81, 511–519 (1984). 10.1063/1.447334

98 Hoover, W. G. Canonical dynamics: Equilibrium phase-space distributions. Physical Review A 31, 1695–1697 (1985). 10.1103/PhysRevA.31.1695

99 Parrinello, M. & Rahman, A. Polymorphic transitions in single crystals: A new molecular dynamics method. Journal of Applied Physics 52, 7182–7190 (1981). 10.1063/1.328693

100 Gowers, R. et al. 98–105 (SciPy).

101 Humphrey, W., Dalke, A. & Schulten, K. VMD: Visual molecular dynamics. Journal of Molecular Graphics 14, 33–38 (1996). 10.1016/0263-7855(96)00018-5

102 He, J., Li, T. & Huang, S.-Y. Improvement of cryo-EM maps by simultaneous local and non-local deep learning. Nature communications 14, 3217 (2023).

103 Li, H. & Durbin, R. Fast and accurate long-read alignment with Burrows–Wheeler transform. Bioinformatics 26, 589–595 (2010). 10.1093/bioinformatics/btp698

104 Winsor, G. L. et al. Pseudomonas Genome Database: improved comparative analysis and population genomics capability for Pseudomonas genomes. Nucleic Acids Research 39, D596–D600 (2011). 10.1093/nar/gkq869

105 Wagner Victoria, E., Bushnell, D., Passador, L., Brooks Andrew, I. & Iglewski Barbara, H. Microarray Analysis of Pseudomonas aeruginosa Quorum-Sensing Regulons: Effects of Growth Phase and Environment. Journal of Bacteriology 185, 2080–2095 (2003). 10.1128/jb.185.7.2080-2095.2003

106 Robinson, M. D., McCarthy, D. J. & Smyth, G. K. edgeR: a Bioconductor package for differential expression analysis of digital gene expression data. Bioinformatics 26, 139–140 (2010). 10.1093/bioinformatics/btp616

107 Szklarczyk, D. et al. The STRING database in 2023: protein–protein association networks and functional enrichment analyses for any sequenced genome of interest. Nucleic Acids Research 51, D638–D646 (2022). 10.1093/nar/gkac1000

108 Jumper, J. et al. Highly accurate protein structure prediction with AlphaFold. Nature 596, 583–589 (2021).

109 Evans, R. et al. Protein complex prediction with AlphaFold-Multimer. bioRxiv, 2021.2010.2004.463034 (2022). 10.1101/2021.10.04.463034

110 Abramson, J. et al. Accurate structure prediction of biomolecular interactions with AlphaFold 3. Nature 630, 493–500 (2024).

